# SH3BP2 regulates the organization of the neuromuscular synapses through protein-driven phase separation

**DOI:** 10.1101/2024.05.23.595491

**Authors:** Bhola Shankar Pradhan, Krzysztof Bernadzki, Yauhen Bandaruk, Anna Siudzińska, Teresa De Cicco, Marta Gawor, Joanna Krzemień, Grzegorz Chodaczek, Said Hashemolhosseini, Tomasz J. Prószyński

## Abstract

The molecular mechanisms underlying the development and maintenance of the neuromuscular junction are poorly understood, even though the malfunction of these specialized synapses is associated with severe genetic and autoimmune disorders. The identity of factors controlling the maintenance mechanisms of postsynaptic acetylcholine receptors (AChR) in high-density has been elusive and is of great interest to the pharma industry, searching for possible new targets for disease interventions. Here, we report the identification of a scaffold protein SH3BP2, which exhibits polyvalent interaction with the dystrophin-glycoprotein complex (DGC) and AChR pentamers, promoting AChR clustering through phase separation. Muscle-specific SH3BP2 deletion in mice leads to impaired organization of the neuromuscular synapses, defects in synaptic transmission, and reduced muscle strength. Our studies identified a novel regulator of the postsynaptic machinery involved in clustering AChR and linking it to the DGC.

## Introduction

Formation of the postsynaptic machinery at the neuromuscular junction (NMJ) is a complex process that leads to the clustering of acetylcholine receptors (AChR) in high density on the surface of muscle fibers underneath nerve termini^1–4^. Decades of studies have established the critical role of scaffold protein rapsyn and the dystrophin-glycoprotein complex (DGC) in this process^5–7^. DGC is a macromolecular transmembrane complex that helps to stabilize membrane and peripheral proteins (including postsynaptic components) by linking them to the extracellular matrix (ECM) and cytoskeleton^8^. Mutations in several DGC proteins often cause severe muscular dystrophies characterized by muscle weakness and muscle mass loss and are often fatal^9–12^. However, the molecular functions of DGC components are not fully understood.

Rapsyn is a peripheral membrane protein that interacts with the DGC and is well-recognized for its function in AChR clustering. The process of AChR aggregation involves direct binding to AChR subunits α and β and self-aggregation of rapsyn through its seven tetratricopeptide repeats (TPR)^13,14^. Consistently with rapsyn pivotal function in AChR aggregation and organization of the postsynaptic machinery, mutant mice lacking rapsyn expression die at birth due to the complete absence of the postsynaptic machinery and respiratory problems^15^. Similarly, patients with mutations affecting rapsyn expression or functions develop congenital myasthenic syndrome^16^. Recent discoveries provided more comprehensive information on the novel functions of rapsyn in the postsynaptic machinery. First, Li et al. demonstrated that the rapsyn RING domain possesses enzymatic activity functioning as an E3 ubiquitin ligase that regulates AChR neddylation and possibly posttranslational modification of other postsynaptic components^17^. Neddylation of AChR was shown to be important for AChR clustering. Second, rapsyn, similarly to the DGC, serves as a scaffold for the attachment of all three types of cytoskeleton – actin, intermediate cytoskeleton, and microtubules^18–20^. Therefore, the current view is that rapsyn plays many important functions in NMJ development besides aggregating AChR, which could contribute to the lethal phenotype in the absence of rapsyn. Additionally, it was recently shown that rapsyn-mediated aggregation of the AChR involves phase separation, a process in which polyvalent protein interaction leads to the formation of a separate domain with unique biophysical properties capable of recruiting other molecules^16^. Phase separation emerged recently as a key process regulating protein interactions in macromolecular assemblies, and transmembrane ECM receptors were proposed to serve a role in orchestrating protein condensates at the cell periphery on the cytoplasmic side^21^. Collectively, these findings, together with other unexpected discoveries, recapitulate our understanding of the principles behind the NMJ organization and highlight the need for further studies^22^.

Our research on the DGC was focused on the α-dystrobrevin-1 (αDB1), a cytoplasmic component of the DGC. This protein plays an important role in the organization of AChR at postsynaptic sites and NMJ development^23^. It has been established that synaptic functions of αDB1 depend on phosphorylation of the C-terminally located tyrosine residues^24–26^. However, the molecular mechanisms through which αDB1 and its phosphorylation exert synaptic functions are unclear, although it was postulated that tyrosine phosphorylation generates new sites for interactions with downstream components.

Here, we report that αDB1 phosphorylation at Y713 recruits a scaffold protein, SH3BP2. This protein has been previously implicated in signaling regulation in immune and bone cells^27–35^, but its function in skeletal muscles has not been reported. We show that SH3BP2 is enriched in muscle postsynaptic machinery and interacts with several DGC components. At the same time, SH3BP2 forms polyvalent interactions with AChR pentamers, promoting their clustering in a process involving phase separation. Muscle-specific knockout of SH3BP2 in mice leads to abnormal NMJ organization and function. Mutant mice have impairments linked to synaptic transmission, muscle strength, and general fitness.

## Results

### SH3BP2 interacts with α-dystrobrevin-1 in a phospho-dependent manner

We have previously reported the interaction of a scaffold protein SH3BP2 (and its interacting adapter protein Grb2) with phosphorylated Y713 of αDB1 (**Fig. 1, A**)^23^. SH3BP2 contains an N-terminal pleckstrin homology (PH) domain, a centrally located SH3-binding proline-rich region, and a C-terminal Src homology 2 (SH2) domain (**Fig. 1, B**). It also has a predicted intrinsic disorder (ID) domain, suggesting a possible interaction with other proteins and selfinteraction. The functions of SH3BP2 are poorly described, and its role in skeletal muscles has not been investigated before.

**Fig. 1.**
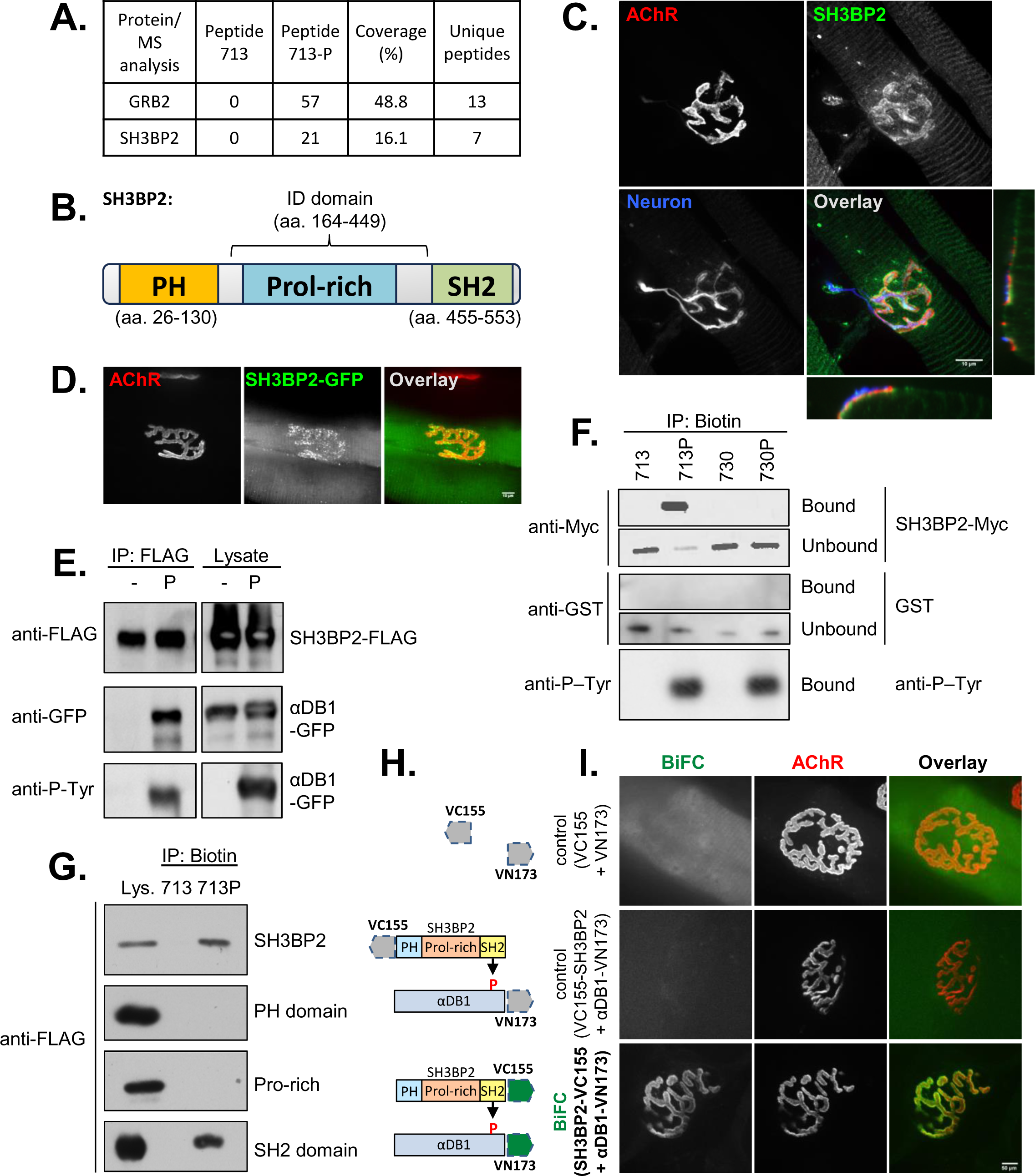
SH3BP2 interacts with phosphorylated αDB1 and localizes to the NMJ. (A) Mass spectrometry identifies SH3BP2 as an interactor of the phospho-Y713 peptide of αDB1^23^. MS/MS analysis shows the spectra identified for SH3BP2 and GRB2 for phosphorylated Y713-P and control Y713 peptides. (B) Schematic illustration of SH3BP2 domains. ID – intrinsically disordered domain. (C) SH3BP2 was localized to the pre- and postsynaptic compartments of the NMJ. XZ and YZ side views are also presented for better visualization of SH3BP2 localization at NMJ. The blue channel shows the motor neuron terminal (Neuron) visualized as a combined neurofilament and synaptophysin immunofluorescence. (D) Localization of SH3BP2-GFP at the NMJ. (E) SH3BP2-FLAG coprecipitates (IP) phosphorylated αDB1-GFP from Hek293 cell lysate in the presence of pervanadate (P). (F) Phosphorylated Y713 (Y713P) αDB1 peptide precipitates purified SH3BP2-MYC, suggesting a direct interaction. Unphosphorylated Y713, Y730, and phosphorylated 730-P peptides of αDB1 were used as negative controls. Purified GST provides unrelated protein control. Precipitation of anti-P-Tyr antibody was used to confirm peptide phosphorylation. (G) 713-P αDB1 peptide precipitates full-length SH3BP2-FLAG and the FLAG-tagged SH2 domain of SH3BP2 protein but not tagged PH domain or proline-rich domain (Pro-rich) of SH3BP2 from Hek293 cell extract. (H and I) A BiFC signal of SH3BP2–VC155 with αDB1–VN173 at the NMJ. Fluorescence complementation was observed when the two Venus fragments were fused C’-terminally αDB1 and SH3BP2, close to the interaction sites. No BiFC was observed when the VC155 fragment was located at the N’-terminal of SH3BP2, demonstrating BiFC signal specificity. Scale bar 10 µm.

To study the localization of SH3BP2 in skeletal muscles, we immuno-stained teased muscle fibers from the Tibialis anterior muscles. Strong SH3BP2 immunoreactivity was detected at the NMJ postsynaptic machinery, which overlapped with AChR (**Fig. 1, C**). This pattern was very similar to the one reported previously for αDB1^23^. To verify the localization of SH3BP2, we electroporated the Tibialis muscle with plasmids encoding GFP-SH3BP2 or GFP alone as a control. As expected, GFP did not localize to the NMJ (**Fig. S1, A**), but GFP-SH3BP2 was strongly enriched at the junctions, although in a bit punctate pattern (**Fig. 1, D**). Also, the fluorescence in a striated pattern along the entire fiber was detected. However, it was weaker than in the immunostaining experiment, possibly due to the elevated background in the overexpression experiment. To investigate this periodic localization of SH3BP2, we co-stained teased muscle fibers with antibodies against Titin, which connects the Z disc to the M line in the sarcomere, α-actinin, which is an actin-binding protein that anchors myofibrillar actin thin filaments and Titin to Z-disc, and ryanodine receptors (Ryr) that are located in sarcoplasmic reticulum abutting Z-discs (**Fig. S1, B**). The co-localization of SH3BP2 was observed for α-actinin, a protein associated with Z-discs.

To validate the interaction between SH3BP2 and phosphorylated αDB1, we performed a coimmunoprecipitation experiment of SH3BP2-FLAG and GFP-αDB1 overexpressed in Hek293 cells. To induce αDB1 phosphorylation, we co-transfected cells with a dominant active mutant of Src kinase and treated cells with phosphatase inhibitor pervanadate, as described previously^23^. As expected, SH3BP2-FLAG co-precipitated GFP-αDB1 from pervanadate-treated cells extract (**Fig. 1E**). Next, we used biotinylated synthetic αDB1 peptides containing centrally located tyrosine Y713 in phosphorylated or unphosphorylated states. The peptides were used to coat magnetic beads and tested for their ability to precipitate purified SH3BP2-MYC protein. Only phosphorylated Y713 peptide efficiently precipitated SH3BP2-MYC (**Fig. 1, F**). As a control, we performed a similar experiment with αDB1 peptides corresponding to the Y730 site, but no affinity for SH3BP2-MYC was observed (**Fig. 1, F**). None of the peptides precipitated purified GST protein, which was used as an additional negative control. On the other hand, both Y713-P and Y730-P could precipitate an anti-phosphotyrosine antibody, demonstrating that both peptides were phosphorylated (**Fig. 1, F**). These experiments demonstrate that SH3BP2 interacts directly and specifically with the Y713 in its phosphorylated state (**Fig. 1, F**).

To examine which of the SH3BP2 domains is responsible for interaction with phosphorylated αDB1, we expressed full-length SH3BP2-FLAG and three domains of SH3BP2 (i.e., PH domain, a proline-rich domain, and SH2 domain) fused to FLAG individually in Hek293 cells and tested for their interaction with either phosphorylated and unphosphorylated Y713 αDB1 peptides. Only the SH2 domain with FLAG was precipitated by the Y713-P peptide of αDB1 (**Fig. 1, G**). This observation is in agreement with the known function for SH2 domains that mediate interactions with phosphorylated tyrosine residues in a sequence-specific manner. To confirm that the interaction between SH3BP2 and αDB1 also occurs in skeletal muscles, we performed bimolecular fluorescence complementation (BiFC) experiments. For this, we electroporated Tibialis muscle with plasmids expressing SH3BP2 and αDB1 fused nonfluorescent portions of the Venus protein. If two proteins of interest interact and bring split Venus sequences in close proximity, then the functional protein can fold and become fluorescent due to bimolecular complementation^36^. As a control, we expressed fragments of Venus alone, but no BiFC signal was observed at the NMJ (**Fig. 1, H and I**). Similarly, no Venus fluorescence was detected when the VC155 tag was placed at the N-terminal of SH3BP2. In contrast, co-electroporation of SH3BP2 with the C-terminally attached VC155 and αDB1 with VN173 attached to the C-terminal side containing phosphorylated tyrosine residues led to strong BiFC signal at the NMJ, which overlapped with AChR (**Fig. 1, H and I**). Therefore, SH3BP2 and αDB1 appear to interact at the NMJ.

To determine the SH3BP2 protein domains involved in NMJ localization, we electroporated the Tibialis muscle of WT mice with plasmids expressing GFP-tagged PH, Proline-rich, and SH2 domain or GFP-tagged truncation constructs missing the N-terminal PH domain or C’-terminal SH2 domain. Both truncation constructs were strongly enriched at the NMJ postsynaptic machinery (**Fig. 2**). However, the protein missing the PH domain was more evenly distributed at the postsynaptic machinery, and the construct lacking the SH2 domain had more punctate distribution. Additionally, fibers expressing the PH-proline-rich construct contained protein aggregates (**Fig. 2**). Surprisingly, all three GFP-tagged individual domains of SH3BP2 had also concentrated at the NMJ. The proline-rich domain had the most even staining, the PH domain the most punctate, and the SH2 domain had a relatively weak signal at the NMJ. Our results suggest that all three domains of the SH3BP2 adapter protein must interact with postsynaptic components that mediate their recruitment to the junction.

**Fig. 2.**
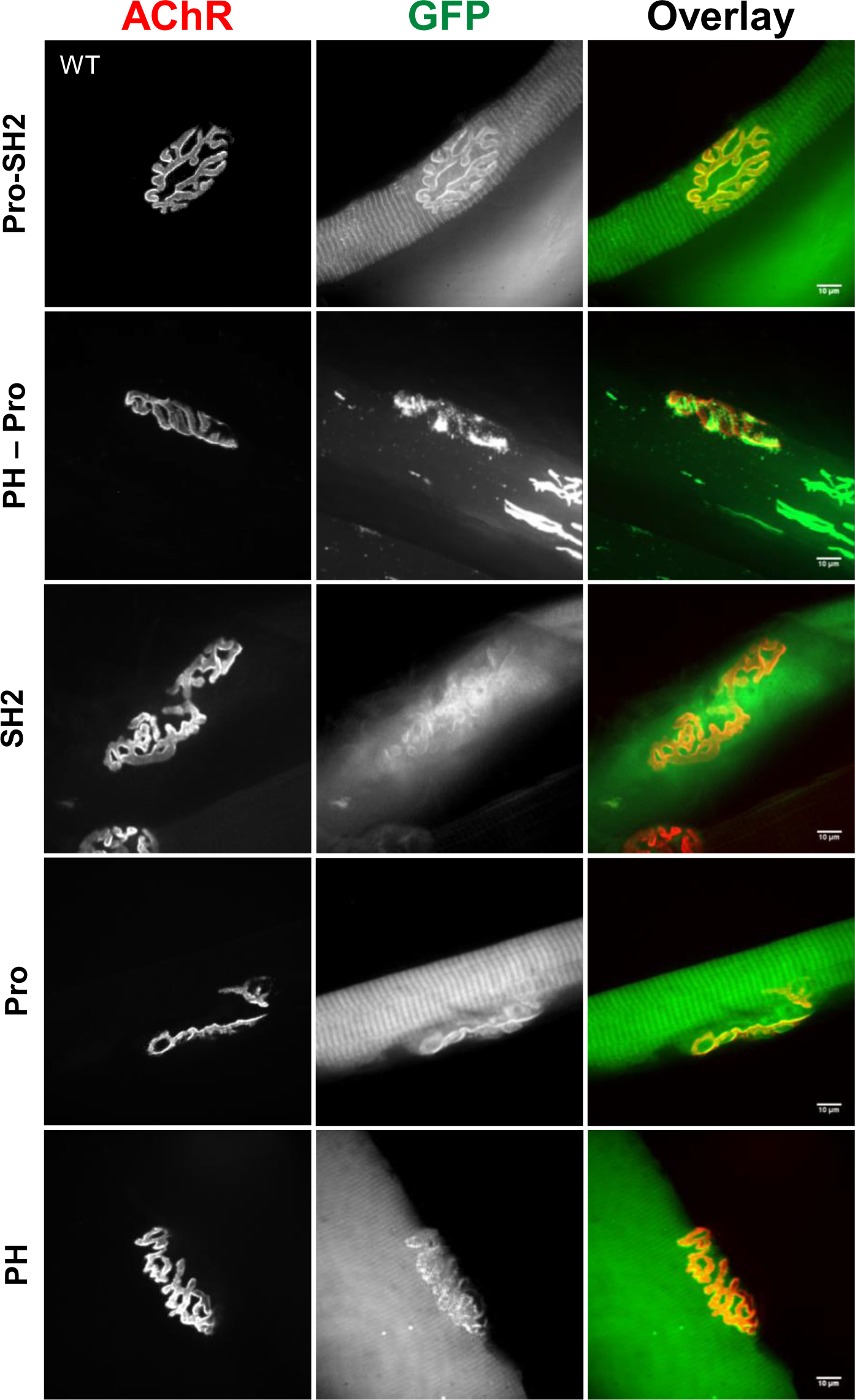
NMJ localization of SH3BP2 truncation proteins. Each domain of SH3BP2 fused to GFP can be targeted to the junction. The Tibialis anterior muscles were electroporated with plasmids expressing GFP-tagged PH domain (PH), proline-rich domain (Pro), SH2 domain (SH2), or a truncated version of SH3BP2 protein missing either the SH2 domain (PH–Pro) or the PH domain (Pro–SH2). Scale bar 10 µm.

### Depletion of SH3BP2 impairs AChR clustering in cultured myotubes

Since we observed that SH3BP2 was localized along with αDB1 to the NMJ’s postsynaptic machinery, we examined whether SH3BP2 plays a role in the organization of AChRs. For this, we knocked down SH3BP2 expression in C2C12 differentiated myotubes using siRNAs validated for their ability to reduce SH3BP2 mRNA and protein levels (**Fig. S2, A and B**). To induce AChR clustering, myotubes were grown on laminin-111-coated surfaces or treated with soluble agrin^37^. AChR clusters formed in the cultured myotubes represent many aspects of AChR assemblies *in vivo*^38,39^. The control laminin-grown myotubes transfected with non-targeting siRNA formed a large number of small plaque-like AChR assemblies and many topologically complex, pretzel-shaped AChR clusters with F-actin enriched podosomes^40^ located at the sites of AChR perforations (**Fig. 3, A**). On the other hand, the agrin-treated control myotubes displayed many immature small AChR clusters (**Fig. 3, B**). Depletion of SH3BP2 in the laminin-111 treated myotubes led to a significant (p <0.001) reduction in both the number of complex clusters and the total number of AChR clusters (**Fig. 3, A, C, and D**). SH3BP2 inhibition in myotubes treated with agrin showed more than 50% reduction in the total number of AChR clusters compared to control (**Fig. 3, B, C, and D**). A similar effect was apparent for the two siRNAs (**Fig. 3, C, D**), arguing against the off-target effects of the siRNAs. We hypothesized that impaired AChR clustering upon SH3BP2 depletion could be due to decreased AChR produced by myotubes, compromised exocytosis, or cells’ inability to cluster AChR into larger assemblies. To test these three possibilities, we performed AChR precipitation experiments in which we labelled surface receptors by incubating live cells with biotin-conjugated bungarotoxin or added bungarotoxin to the cell homogenate, allowing for labelling total AChR. The amounts of surface and total AChR subunits α and δ precipitated with streptavidin magnetic beads were comparable between the control and SH3BP2-depleted myotubes (**Fig. 3, E**). These results suggest that the lack of SH3BP2 does not affect the expression of AChR or its transport to the cell surface. Instead, myotubes depleted from SH3BP2 have compromised the ability to cluster receptor molecules at the plasma membrane.

**Fig. 3.**
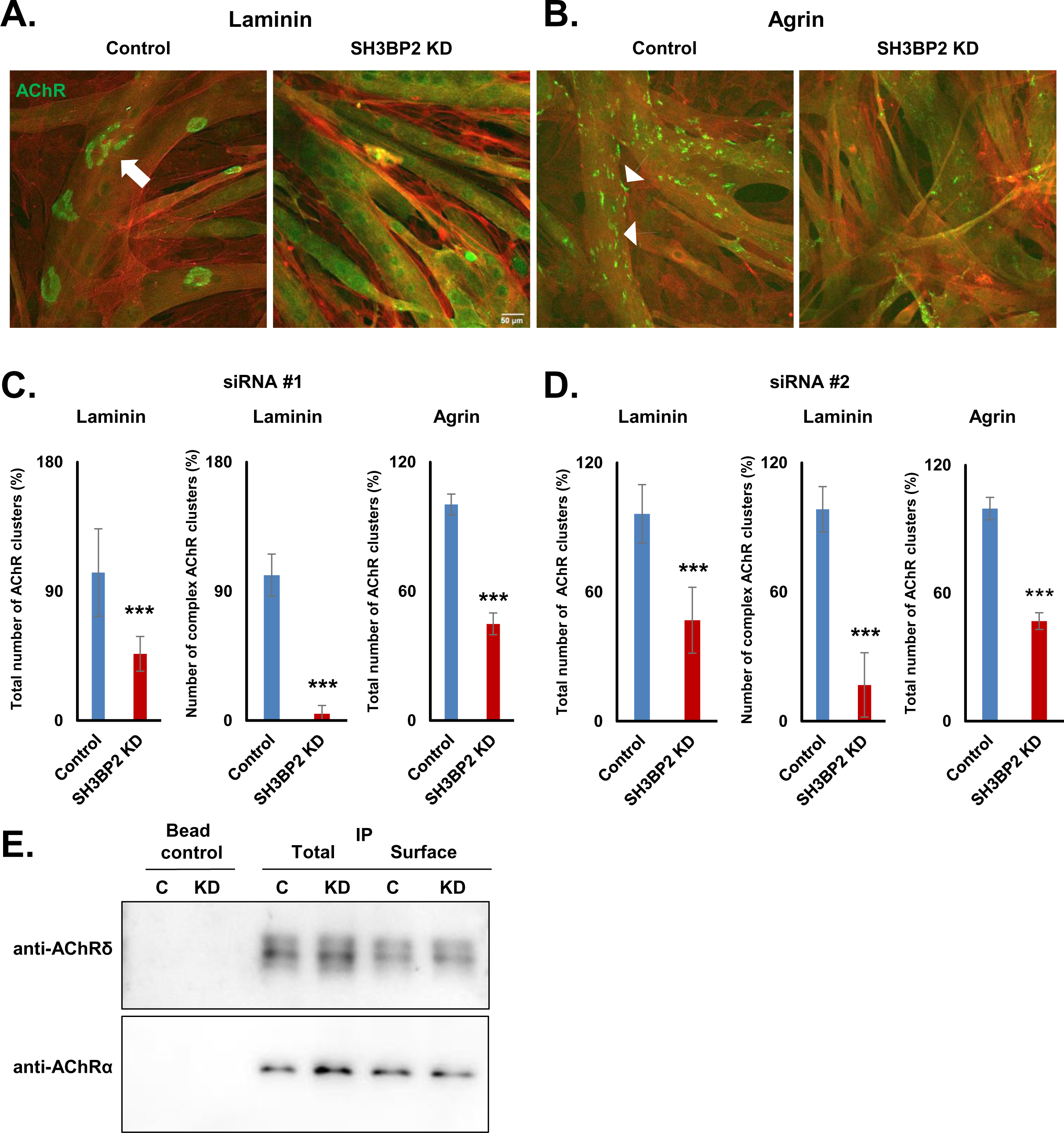
Depletion of SH3BP2 in myotubes affects AChR clustering. (A and B) Effect of SH3BP2 knock-down on AChR clustering in C2C12 myotubes grown on laminin-111 (A) or treated with soluble agrin (B). The arrow indicates an AChR cluster with complex morphology, and arrowheads point to small clusters induced by agrin. F-actin stained with phalloidin is shown in red as counterstaining. Control – non-targeting siRNA. Scale bar 50 µm. (C and D) The quantification of the total number of AChR clusters and the number of complex clusters (for laminin-111) in myotubes transfected with scramble siRNA (control) and SH3BP2-targeting siRNA #1 (C) and #2 (D). Error bars represent SEM; ***p < 0.001, two-tailed t-test. (E) Depletion of SH3BP2 in C2C12 myotubes did not affect the total AChR levels or surface AChR levels as detected by pull-down of AChR using biotin-bungarotoxin conjugates and neutravidin agarose. Beads control – precipitations from cell extract without bungarotoxin. C – control; KD – SH3BP2 knockdown.

### SH3BP2 interacts with DGC complexes and AChR subunits α and γ

Our results demonstrated that SH3BP2 localizes to the NMJ, interacts with αDB1, and regulates AChR clustering. To gain insight into underlying mechanisms, we isolated SH3BP2-interacting proteins from differentiated myotubes and performed the identification of proteins using mass spectrometry. For this, we coated magnetic beads with purified SH3BP2 and incubated them with a homogenate of laminin-grown C2C12 myotubes. After washing, beads were suspended in the sample buffer, and isolated proteins were resolved using SDS-PAGE electrophoresis (**Fig. 4, A**) and identified using mass spectrometry. Among the identified proteins were several postsynaptic proteins such as α and γ subunits of AChR, Low-density lipoprotein 4 (Lrp4), and several components of the dystroglycan complex, including utrophin, dystrophin, and α and β syntrophins (**Fig. 4, B and Supplementary Table 1**). To validate some of these interactions, we performed a set of co-immunoprecipitation experiments. First, we wanted to know if SH3BP2 interacts with DGC components besides αDB1. As shown in **Fig. 4, C**, syntrophin-1-GFP could co-precipitate SH3BP2-FLAG when the two proteins were expressed in Hek293 cells. Next, we decided to extend our analysis for β-dystroglycan, which provides a scaffold for the assembly of the entire DGC. Our results show that β-dystroglycan-GFP could also co-precipitate SH3BP2-FLAG from Hek293 cell lysate (**Fig. 4, D**). Therefore, SH3BP2 appears to interact with several components of the DGC. This property of SH3BP2 was similar to its binding partner, another adapter protein, Grb2, which has been shown to interact with the β-dystroglycan, αDB1, and AChR^23,41^.

**Fig. 4.**
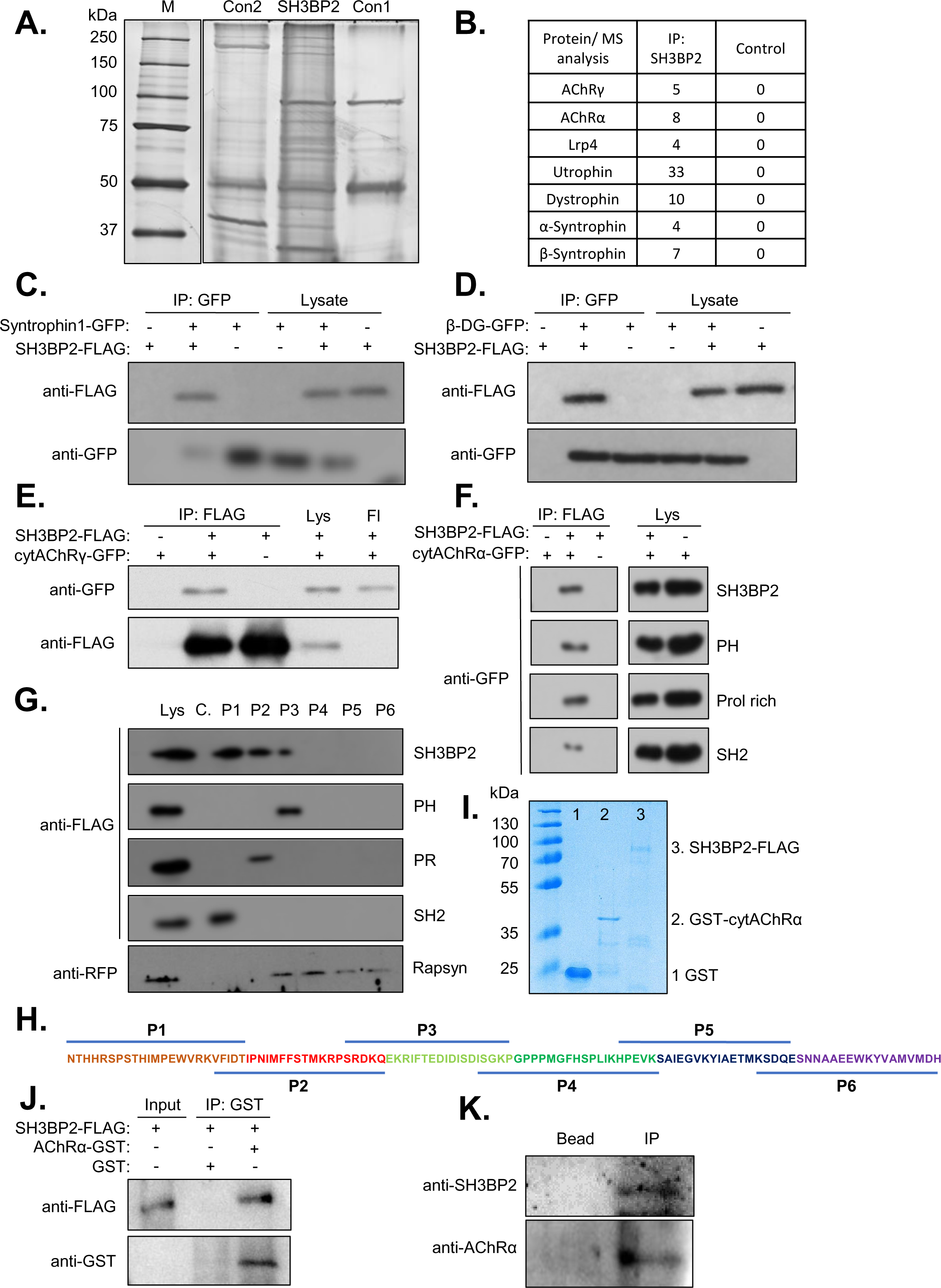
SH3BP2 interacts with syntrophin-1, β-dystroglycan, subunits α and γ of AChR. (A) Silver staining of the SDS-PAGE gel with proteins co-purified with SH3BP2 for mass spectrometry analysis. Con1 shows a control 1 – TAP-tag-GFP-SH3BP2 purified from Hek293 cells without exposure to C2C12 lysate; Con 2 represents a control 2 – IgG Dynabeads incubated with myotube extract without precoating with SH3BP2 protein; M - molecular weight marker. (B) Selected proteins that were specifically co-purified with TAP-tag-GFP-SH3BP2 were identified by MS/MS analysis. IP: SH3BP2 and Control columns show the number of peptides identified for each protein indicated in the left column. The control column shows the cumulative amount of peptides identified in both control samples. (C) Syntrophin-1 coprecipitates SH3BP2-FLAG from Hek293 cells extract. (D) β-dystroglycan cytoplasmic domain fused to GFP coprecipitates SH3BP2-FLAG from Hek293 cells homogenate. (E) SH3BP2-FLAG coprecipitates the cytoplasmic domain of AChR γ-subunit fused to GFP from Hek293 cells lysate. Lys – lysate of transfected Hek293 cells; Fl – unbound fraction. (F) SH3BP2-FLAG coprecipitates cytAChRα-GFP from Hek293 cell extract. Similarly, each domain: the PH domain (PH), proline-rich (PR), and SH2 domain (SH2) fused to FLAG coprecipitated cytAChRα from Hek293 cells lysate (Lys). (G) Coprecipitation of SH3BP2-FLAG, its individual domains fused to FLAG or Rapsyn-Cherry (positive control) from Hek293 cell extract with synthetic peptides covering the cytoplasmic domain of AChRα depicted in (H). Biotinylated peptides were attached to streptavidin Dynabeads and used to test their ability to precipitate indicated proteins. (I) Safe blue staining of purified GST (1), GST-tagged cytAChRα (2), and SH3BP2-FLAG (3). (J) Coprecipitation experiment showing direct interaction of purified SH3BP2-FLAG protein with the purified cytoplasmic domain of AChRα fused to GST (GST-cytAChRα). Precipitation was performed with anti-GST antibody, and precipitation of GST in the presence of SH3BP2-FLAG served as a negative control. (K) Endogenous AChR coprecipitates endogenous SH3BP2 from C2C12 myotubes lysate. Surface AChR was labeled in live C2C12 myotubes with biotin-bungarotoxin and precipitated from cell homogenates with neutravidin agarose. Beads – precipitates without bungarotoxin.

Our mass spectrometry analysis suggested interactions between SH3BP2 and AChR subunits α and γ. Since SH3BP2 is a cytoplasmic protein, its interaction should be mediated by the cytoplasmic domains of the receptor subunits. Each AChR subunit has four transmembrane domains (TMD), both terminals exposed to the outside of the cells, and two cytoplasmic domains. However, the cytoplasmic loops between the TMD I and II are believed to be too short to recruit any interacting proteins (seven and eight amino acids in mouse AChR α and γ subunits, respectively). Therefore, to test the interaction of SH3BP2 with AChR, we expressed in Hek293 cells the sequences encoding the cytoplasmic loops between TMD III and IV of AChRα (111 amino acids) and γ (146 amino acids) fused to GFP (named cytAChRα-GFP and cytAChRγ-GFP). Both proteins were co-precipitated by SH3BP2-FLAG from cell homogenates (**Fig. 4, E, and F**). We decided to focus our further analysis on the AChRα sequence since it is the obligatory subunit present in two copies in the AChR pentamer. In contrast, the γ subunit is present in pentamers during embryonic development and is exchanged for the adult ε subunit in the postnatal life^42^. To study which domain of SH3BP2 binds to AChRα, we performed co-immunoprecipitation experiments between the cytAChRα-GFP and PH domain, Proline-rich domain, and SH2 domain of SH3BP2 fused to FLAG. Interestingly, all three domains of the SH3BP2 protein showed interaction with cytAChRα (**Fig. 4, F**). This result is in agreement with our previous observation that all three domains of SH3BP2 can be recruited to the NMJ postsynaptic machinery independently (**Fig. 2**). To map the binding sites on the AChRα sequence, we examined the interaction between SH3BP2 and its domains with biotin-conjugated synthetic peptides that cover the entire sequence of the AChRα cytoplasmic loop in a partially overlapping way (**Fig. 4, G and H**). The full-length SH3BP2-FLAG protein was precipitated from Hek293 extract by magnetic streptavidin beads coated with peptides P1, P2, and P3 but not P4, P5, and P6, suggesting that the interaction with SH3BP2 occurs at the N’-terminal part of the cytAChRα (**Fig. 4, G and H**). Interestingly, each SH3BP2 domain was recruited to a different peptide, with the SH2 domain being precipitated by P1, the Proline-rich by P2, and the PH domain by the P3 peptide (**Fig. 4, G**). This suggests that the SH3BP2 protein has three binding sites on AChRα and that each site interacts with different domains of the SH3BP2 protein. As a control, we examined the ability of the peptides to precipitate the mCherry-tagged version of rapsyn from the Hek293 cell homogenate. Rapsyn is a scaffold protein that directly interacts with AChRα^43^. As expected, the rapsyn fusion protein was also precipitated by the peptides. However, its interaction was specific for peptides P3 to P6 covering the C-terminal fragment of the cytAChRα (**Fig. 4, G, and H**).

To examine if the interaction between AChRα and SH3BP2 is direct, we purified GST or GST-cytAChRα from bacteria (**Fig. 4, I**) and SH3BP2-FLAG from Hek293 cells. We performed precipitation from mixed protein solution using anti-GST coated Dynabeads. As shown in **Fig. 4, J**, GST-cytAChRα protein, but not GST alone, coprecipitated SH3BP2-FLAG, suggesting a direct interaction between these two proteins. To confirm that SH3BP2 interacts with AChR in myotube cells, we incubated live myotubes with biotin-conjugated bungarotoxin for 5 min to label the surface AChR and isolated them using NeutrAvidin beads that were added to the cell lysate. Western blot analysis of isolated proteins showed that AChR co-precipitates endogenous SH3BP2 (**Fig. 4, K**).

Collectively, our experiments revealed that SH3BP2 protein binds multiple components of the DGC, and at least interaction with αDB1 is direct. Additionally, they showed that SH3BP2 binds to at least two subunits of the AChR and that the interaction with AChRα is direct and polyvalent.

### SH3BP2 forms liquid-liquid phase separation

Phase separation is induced by proteins exhibiting polyvalent interactions, including those that oligomerize^44,45^. A previous pioneering study from Lin Mei’s laboratory has shown that rapsyn clustering triggers protein droplet formation recruiting AChR and that this mechanism contributes to receptor clustering at the NMJ^16^. Polyvalent interactions have often been reported for intrinsically disordered domain-containing proteins^16,46,47^. Since SH3BP2 contains an intrinsically disordered region (**Fig. 1, B**), we hypothesized that it could be responsible for self-aggregation triggering phase separation. To test this hypothesis, we performed co-immunoprecipitation for SH3BP2-MYC and EGFP-SH3BP2 expressed in Hek293 cells. This experiment demonstrated that SH3BP2-MYC can isolate EGFP-SH3BP2 (**Fig. 5, A**), suggesting that SH3BP2 can self-interact. For further studies, we purified the GST-EGFP-tagged SH3BP2 protein from transfected Hek293 cells extract (**Fig. 5, B**) and tested its ability to form liquid droplets *in vitro* at different protein and salt concentrations. At 500 mM NaCl, the GST-EGFP-SH3BP2 protein in 5 µM concentration had an even distribution in the confocal microscope field of view, as a high salt concentration disrupts protein interactions (**Fig. 5, C**). When the salt concentration was reduced to 125 mM, fluorescent protein aggregates became apparent (**Fig. 5, C**). The formation of fluorescent puncta depended on the protein concentration; it started at 1 µM and was the most prominent at 5 µM GST-EGFP-SH3BP2 (**Fig. 5, C**). Protein concentration also influenced the size, number of fluorescent puncta, and the kinetics of fluorescent puncta formation. At 5 µM, protein puncta were the most prominent, with up to 2 µm in diameter, and formed immediately after salt concentration decreased (**Fig. 5, C and D**). At lower protein concentrations, the puncta were smaller, and their formation required more time (**Fig. 5, C, and D**). Since SH3BP2 contains the ID domain expected to trigger protein aggregation and liquid-liquid phase separation, we purified from Hek293 cells the SH3BP2 mutant protein missing the ID domain (**Fig. 1, B and 5, B**). As expected, this protein failed to form protein aggregates (**Fig. 5, C**). We also investigated protein aggregation in centrifugation experiments (14,000 x g, 15 min) that allowed for the precipitation of the aggregates. Similarly to our microscopy experiments, little protein was detected in the pellet at 500 mM NaCl (**Fig. 5, E and F**). Significant protein amounts were detected in the pellets when the salt concentration was decreased to 125 mM, suggesting that the protein formed higher-weight assemblies (**Fig. 5, E, and F**).

**Fig. 5.**
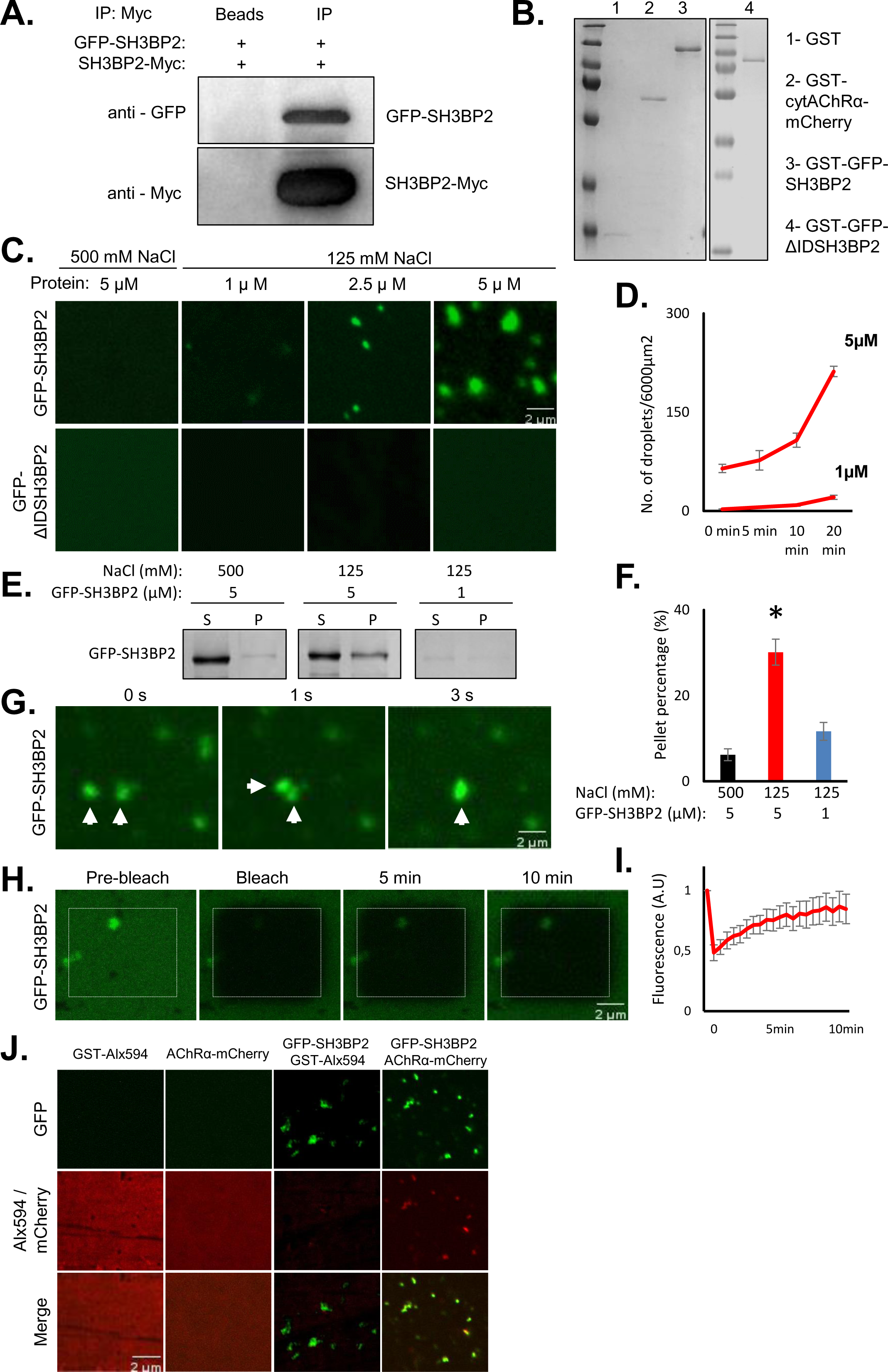
SH3BP2 forms protein condensates and recruits AChRα. (A) SH3BP2-MYC coprecipitates GFP-SH3BP2 from Hek293 homogenates. Beads – control precipitation without anti-MYC antibody. (B) Safe blue staining of purified GST, GST-cytAChRα-mCherry, GST-GFP-SH3BP2, and GST-GFP-ΔIDSH3BP2 showing purity of proteins. (C) Representative images showing that GST-GFP-SH3BP2 (GFP-SH3BP2), but not GST-GFP-ΔIDSH3BP2 (GFP-ΔIDSH3BP2) without intrinsic disorder (ID) domain phase separate into condensed droplets. Protein droplet formation was dependent on the NaCl and protein concentrations. (D) Quantification of droplet number at different time points (0 min, 5 min, 10 min, and 20 min) at 1 μM and 5 μM protein concentrations and 125 mM NaCl. (E) Condensation assay based on centrifugation shows increased amounts of GST-GFP-SH3BP2 (GFP-SH3BP2) in the pellets in a salt and protein concentration-dependent manner. S - supernatant, P - pellet. (F) Quantification of GST-GFP-SH3B2 (GFP-SH3BP2) in pellets from an experiment in E. Results are mean ± SEM; one-way ANOVA; ***p < 0.001, *p < 0.05; n=3. (G) Fusion of two GST-GFP-SH3BP2 (GFP-SH3BP2) droplets. (H) A representative image of the FRAP experiment showing the condensed droplets of GST-GFP-SH3BP2 (GFP-SH3BP2) exchanged with the surrounding aqueous phase. (I) Quantification of fluorescence recovery after photobleaching of SH3BP2 droplets. (J) GST-GFP-SH3BP2 (GFP-SH3BP2) protein droplets recruit GST-cytAChRα-mCherry (AChRα-mCherry) but not GST-Alexa Fluor 594 (right columns). The left columns show negative controls - GST-Alexa Fluor 594 and GST-cytAChRα-mCherry (AChRα-mCherry) failed to form protein droplets at 5 μM concentration without SH3BP2. Scale bars 2 μm.

To test the dynamics of SH3BP2 protein aggregates, we performed a time-lapse and fluorescence recovery after photobleaching (FRAP). Detailed analysis revealed that SH3BP2 protein aggregates are dynamic, and fluorescent puncta can undergo fusion within seconds (**Fig. 5, G**). When the fluorescence of individual puncta was photobleached, it was gradually recovered over time (**Fig. 5, H and I**). These results suggest that SH3BP2 aggregates are dynamic with a high rate of molecule exchange between condensates and surrounding liquid, characteristic of protein droplets.

Since SH3BP2 can form protein droplets, we asked whether they can recruit AChR. For this purpose, we purified from Hek293 cell the cytoplasmic domain of the AChRα subunit fused to the GST and mCherry (GST-cytAChRα-mCherry) (**Fig. 5, B**). For the control experiment, we purified GST (**Fig. 5, B**), which was subsequently labeled with Alexa Fluor 594. Neither GST-cytAChRα-mCherry nor GST-Allexa594 formed protein droplets (**Fig. 5, J**). However, when incubated with phase separating SH3BP2, the GST-cytAChRα-mCherry was recruited to protein droplets with mCherry signal overlapping with GFP fluorescence (**Fig. 5, J**). The interaction with SH3BP2 was specific for AChR since GST-Allexa594 was not targeted to SH3BP2 protein droplets (**Fig. 5, J**). Our results demonstrated that SH3BP2 can self-aggregate, forming dynamic protein droplets that can recruit AChRα.

### SH3BP2 promotes AChR clustering in cultured heterologous Hek293 cells

Since the SH3BP2 was binding directly to the cytoplasmic domain of AChRα and recruited it to the protein droplets, we determine whether SH3BP2 can cluster the AChR in heterologous cells that do not express most of the postsynaptic machinery components. To test this hypothesis, we expressed in Hek293 cells AChR subunits α, β, δ, and ε that are needed to form AChR pentamer. AChR at the cell surface was visualized by incubation with fluorescently labeled bungarotoxin. As reported previously, AChR had diffused distribution at the plasma membrane of transfected cells (**Fig. 6, A**). As a positive control, we co-transfected cells with Rapsyn-mCherry. As expected, rapsyn was targeted to the cell periphery and efficiently clustered AChR to the discrete patches (**Fig. 6, A, B, and C**). Previous reports suggest that the myristylation signal of rapsyn is important for AChR clustering activity^48^. SH3BP2 does not have the myristoylation signal, and Hek293 cells transfected with EGFP-SH3BP2 showed the GFP signal accumulating inside the cells **(Fig. 6, A and B**). Therefore, we fused a membrane targeting tag (from the Lyn protein) to the EGFP-SH3BP2 sequence to promote its membrane localization **(Fig. 6, A**). We found that the expression of the membrane form of EGFP-SH3BP2 significantly increased AChR clustering in Hek293 cells (**Fig. 6 B and C**). Next, we examined if the ID motif of the SH3BP2 protein, which mediates protein droplet formation, is required for AChR clustering. For this, we expressed in Hek293 cells a membrane-targeted GFP-ΔIDSH3BP2 protein lacking the ID sequence. The mutant construct was efficiently expressed and targeted to the plasma membrane, but its ability to cluster AChR was substantially compromised (**Fig. 6, A, B, C**). These results confirm that SH3BP2 can promote AChR clustering in heterologous cells and suggest the critical role of the ID domain in this process.

**Fig. 6.**
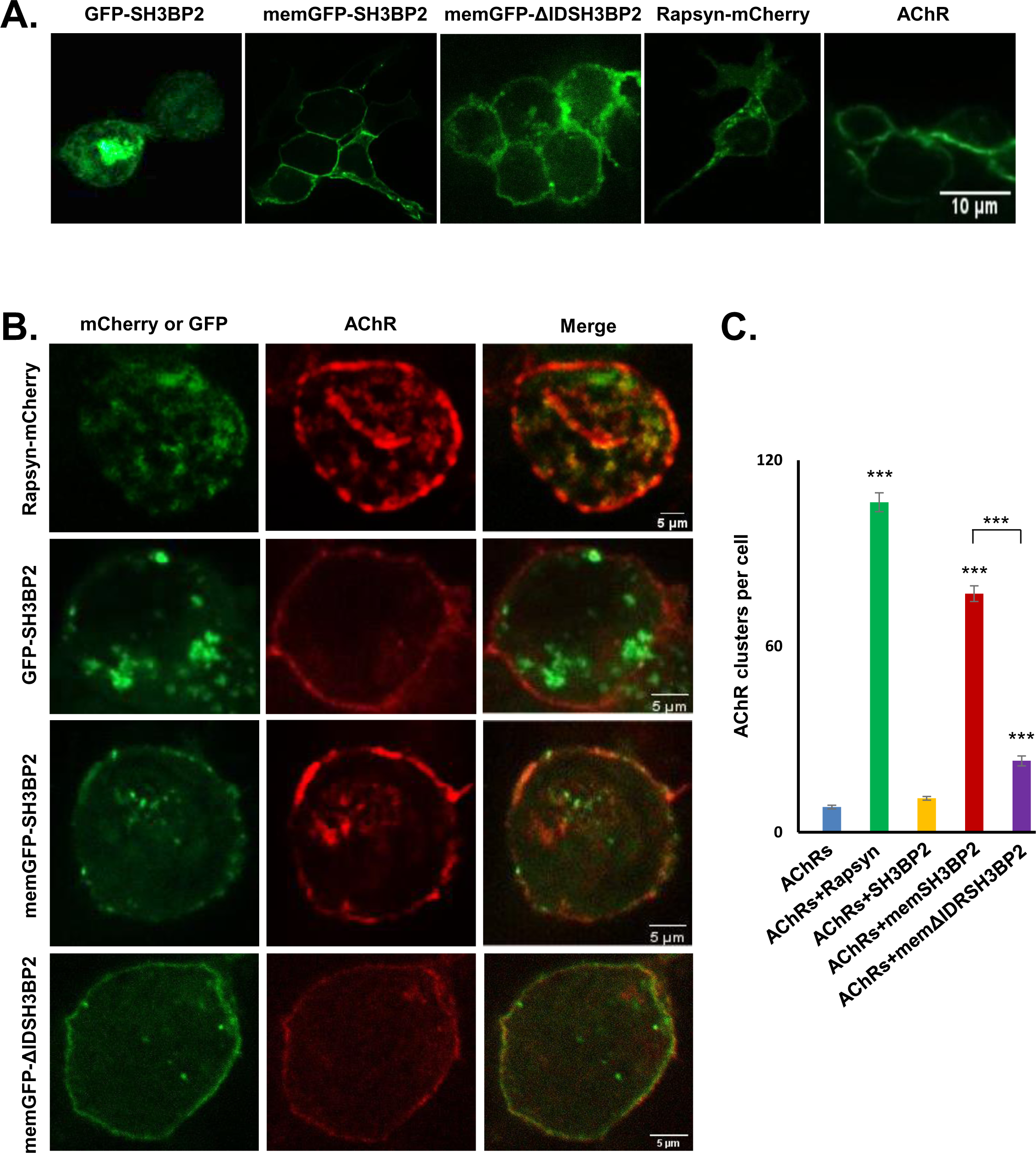
SH3BP2 clusters AChR in Hek293 cells. (A) Representative images of Hek293 cells expressing GFP-SH3BP2, membrane-targeted SH3BP2-GFP (memGFP-SH3BP2), memGFP-SH3BP2 without the intrinsic domain (memGFP-ΔIDSH3BP2), Rapsyn-mCherry, and AChR pentamers (subunits α, β, δ, and ε). Scale bar 10 µm. (B) AChR clustering assay in Hek293 cells. Rapsyn and MemGFP-SH3BP2 were able to cluster AChR on the cell surface. Without a membrane tag, GFP-SH3BP2 localized in the cytoplasm and failed to aggregate AChR. MemGFP-ΔIDSH3BP2 showed substantially decreased AChR clustering ability. Scale bar 5 µm. AChR was visualized by labeling live cells with bungarotoxin. (C) Quantitative analysis of AChR clusters per cell. N=20 cells from three independent experiments; data were shown as mean ± SEM. ***p < 0.001, one-way ANOVA.

### SH3BP2 plays an important role in the NMJ organization

Next, we asked whether SH3BP2 acts as a regulator of the NMJ postsynaptic machinery in vivo. We used conditional SH3BP2 knockout (SH3BP2 fl/fl) mice and employed two strategies for ablating SH3BP2 expression. One was based on crossing SH3BP2 fl/fl mice with Myf5-Cre knock-in mice (**Fig. 7, A**) expressing Cre recombinase under the control of *Myf5* regulatory elements^49^, leading to a muscle-specific SH3BP2 deletion (SH3BP2 mKO). In the second strategy, we delivered Cre recombinase to the Tibialis anterior muscle of SH3BP2 fl/fl mice by AAV-based infection.

**Fig. 7.**
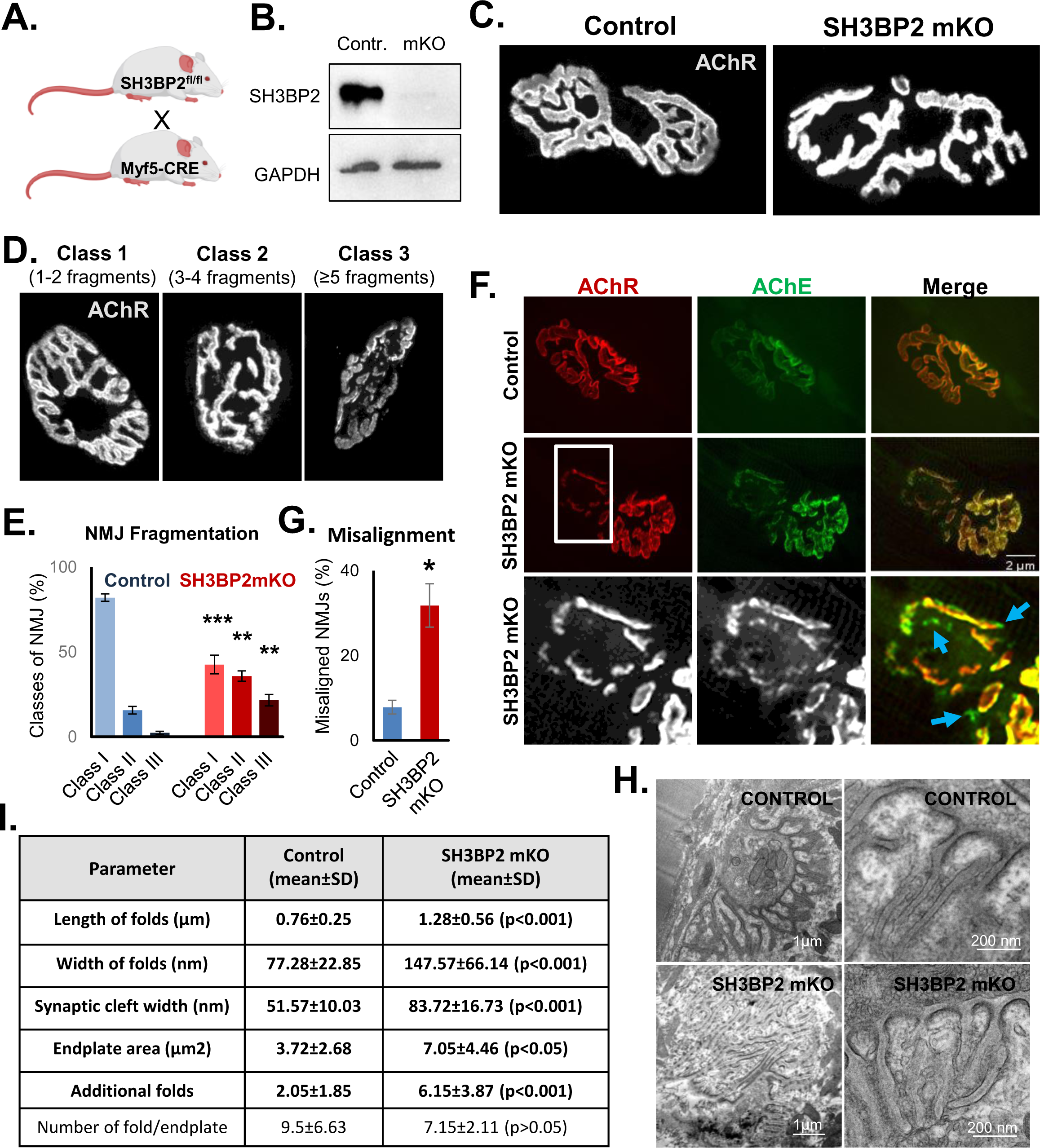
Abnormal NMJ organization in SH3BP2 mKO mice. (A) Strategy for generation of muscle-specific SH3BP2 knockout mice (SH3BP2 mKO). (B) Western blot analysis showing SH3BP2 protein amounts substantially decreased in SH3BP2 mKO Tibialis anterior muscle homogenate. (C) Increased fragmentation of the NMJ postsynaptic machinery in the Tibialis anterior of SH3BP2 mKO mice. (D) Classification of NMJs according to their degree of fragmentation (Class I = 1-2 fragments, Class II = 3-4 fragments, and Class III – more than 4 fragments). (E) Quantification of NMJ fragmentation. Results are mean ± SEM; n=5 mice per group; more than 132 NMJ per group; ***p < 0.001, *p < 0.05; one-way ANOVA. (F) Misalignment of AChR and acetylcholine esterase (AChE) in the soleus of SH3BP2 mKO mice. The lower panel shows magnification of the area in the middle panel; blue arrows show areas containing AChE without AChR; scale bar 2 µm. (G) Quantification of the number of NMJs with misaligned AChR and AChE. Results are mean ± SEM; n = 4 mice per group; more than 100 NMJs were analyzed per genotype; *p < 0.05; unpaired t-test. (H) Representative electron microscopy images of NMJs in Tibialis anterior muscles showing increased length and width of junctional folds in SH3BP2 mKO. Mutant mice also exhibit mislocalization of the electron-dense structures from the crests into the depts of exuberant folds (lower left image). Scale bars were 1 µm (left column) and 200 µm (right column). (I) Quantification of the various NMJ parameters from electron microscopy analysis. The length and width of synaptic folds, as well as the width of the synaptic cleft, are larger in SH3BP2 mKO mice (p < 0.001). Similarly, the area of synaptic gutters (endplate area) and the number of membrane invaginations that were not associated with synaptic gutters (additional folds) was increased in the absence of SH3BP2 (p < 0.001). On the other hand, the number of folds per endplate was not significantly affected (p > 0.05). Results are mean ± SD; n = 4 mice per group; unpaired Student’s t-test.

Homozygous SH3BP2 fl/fl mice bearing the Myf5-Cre transgene were viable and born in accordance with the Mendelian ratio. Western blot analysis of Tibialis muscle homogenate revealed a substantial reduction of SH3BP2 expression (**Fig. 7, B**). The residual expression likely arises from non-muscle cells or incomplete excision of floxed SH3BP2 sequences.

Microscopic analysis of teased Tibialis anterior fibers from SH3BP2 mKO mice revealed that neurofilament architecture was similar in control (SH3BP2 fl/fl) and SH3BP2 mKO mice (**Fig. S3, A**). However, NMJs appeared more fragmented in mutant mice than in control animals (**Fig. 7, C**). To quantify this effect, we acquired images of more than 100 NMJs from five animals per genotype and performed unbiased automated synaptic morphology analysis using an NMJ morph, an algorithm that is used by ImageJ^50^. For fragmentation analysis, we categorized NMJs into three classes: class I of synapses with one to two fragments, class II with three to four fragments, and class III with five or more fragments (**Fig. 7, D**). Our analysis revealed that SH3BP2 mKO mice have significantly reduced numbers of class I NMJs and increased numbers of class II and III junctions (**Fig. 7, E**). At the same time, synapses in the Tibialis muscles had normal size, the area occupied by AChR or occupied by the perforations in the junctions (**Fig. S3, B, C, and D**). Similarly, NMJs in the diaphragm and slow-twitching soleus muscles were more fragmented in SH3BP2 mKO, but other junctional morphology parameters were normal (**Fig. S3, E-L**).

In a parallel experimental strategy, we first infected STOP-Tomato Cre reporter mice with a virus expressing Cre and GFP to test the efficiency of AAV infection of Tibialis muscle and Cre activity. Virus infection led to robust GFP and tdTomato expression in muscle fibers (**Fig. S4, A**). We then used the same AAV to infect the Tibialis muscle in SH3BP2 fl/fl mice and WT animals that served as a control. To verify the excision of SH3BP2 floxed sequences, we performed muscle genotyping and Western blot analysis. Expected PCR product around 500 bp corresponding to the deleted floxed sequence was apparent for muscle genomic DNA from the Tibialis muscle of SH3BP2 fl/fl mice infected with AAV-CRE but not from the uninfected contralateral leg or the AAV-infected leg of WT mice (**Fig. S4, B**). Western blot analysis confirmed the reduction of SH3BP2 expression after viral infection in SH3BP2 fl/fl mice and normal expression levels in the uninfected leg or AAV-infected muscle from WT mice (**Fig. S4, C**). For NMJ morphological analysis, we performed viral infection at two time points in development. The first group of animals was treated at postnatal day 0 (P0), and tissues were collected two months later (**Fig. S4, E, and G**). In the second group, the virus was administered at P60, and muscles were collected at P90 (**Fig. S4, F, and H**). Automated synapse analysis with the NMJ Morph software revealed that in both age groups, SH3BP2 fl/fl mice infected with AAV-CRE have substantially fragmented NMJs compared to WT mice injected with AAV-CRE (**Fig. S4, D, G, and H**). Thus, in both experimental strategies, SH3BP2 deletion led to NMJ morphological abnormalities, and experiments with virus infection additionally demonstrated that SH3BP2 plays a function in the maintenance of synaptic organization independent from development since at P60 NMJ postnatal developmental remodeling is completed^51,52^.

Fragmentation of the NMJ could be associated with the gradual removal of the AChR from the synapse, leading to decreased continuity of synaptic architecture^53,54,55^. We, therefore, performed an analysis of the localization of the acetylcholine esterase (AChE), which is released by the muscle to the synaptic cleft where it is associated with the ECM and, therefore, remains much longer at the junction after other components are removed^26^. Teased fibers from the Tibialis muscle stained with fasciculin II to visualize AChE and bungarotoxin to visualize AChR in WT mice had a good alignment of the two synaptic components (**Fig 7, F and G**). In contrast, muscles from SH3BP2 mKO mice have discrete synaptic regions that were misaligned, containing AChE without AChR (**Fig 7, F and G**). SH3BP2 mKO mice showed almost four times more NMJs with visible misalignment of the synaptic components in the Tibialis muscles than control mice.

Our results encouraged us to study the ultrastructural organization of the NMJs in SH3BP2 mKO using electron microscopy. As shown in **Fig. 7, H and I**, mutant mice frequently had exuberant postsynaptic folds of abnormal length and thickness. Also, the postsynaptic density localization was not limited to the crests of synaptic folds but was frequently located in patches along the exuberant folds (**Fig. 7, H, lower left panel**). SH3BP2 mKO mice also formed more folds outside synaptic gutters (additional folds) (**Fig. 7, I**). The width of the synaptic cleft was also significantly increased (**Fig. 7, I**). Therefore, the lack of SH3BP2 expression leads to substantial changes in NMJ general morphology, molecular organization, and ultrastructure of the junctions.

### Reduced muscle strength in the absence of SH3BP2

To study if the synaptic defects in the absence of SH3BP2 are associated with impaired muscle strength, we performed grip strength analysis and running ability on voluntary running wheels and a treadmill. As shown in **Fig. 8, A**, SH3BP2 mKO mice had significantly decreased grip strength measured for all paws compared to SH3BP2 fl/fl (control) mice. Similarly, SH3BP2 fl/fl mice with AAV-Cre delivered bilaterally to Tibialis muscles have reduced grip strength on hind limbs compared to WT mice infected with the AAV-Cre (**Fig. S4, I**). SH3BP2 mKO mice and SH3BP2 fl/fl infected with AAV-Cre also run much less on the voluntary running wheels (**Fig. 8, B and S4, J**). In the following experiments, we tested animals’ fatigue on a treadmill operating at an accelerating speed. The experiment lasted until animals became exhausted, defined as refusing to run for 15 consecutive seconds despite an aversive stimulus in the form of the air puff. SH3BP2 mKO mice and SH3BP2 fl/fl treated with AAV-Cre become fatigued faster after running for shorter distances and after reaching lower maximal speed than control animals (**Fig. 8, C, D, and E and S4, K, L, and M**).

**Fig. 8.**
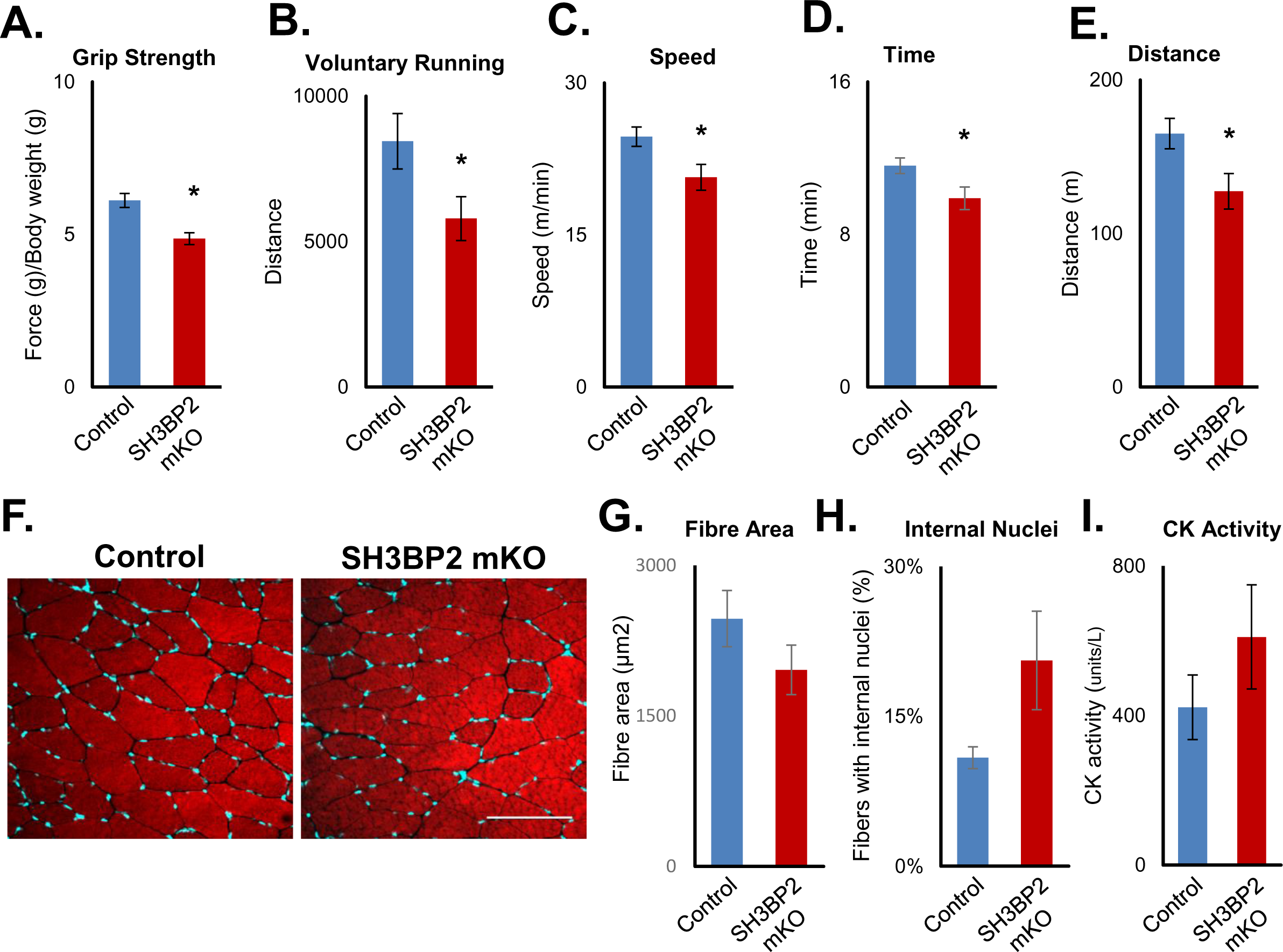
Reduced muscle strength but normal muscle fiber morphology in SH3BP2 mKO mice. (A) Decreased grip strength of SH3BP2 mKO mice. (B) Reduced voluntary running distance by SH3BP2 mKO mice in running wheel experiment. (C-E) SH3BP2 mKO mice have reduced fitness in the treadmill experiments. Mutant mice ended the test reaching lower maximal speed (C), shorter running time (D), and distance (E). *p < 0.05; n = 10 mice per group; unpaired t-test. (F) Representative images of Tibialis anterior muscle cross sections stained with phalloidin for F-actin (red) and DAPI (cyan) to visualize nuclei. Quantification of (G) muscle fiber diameter and (H) fibers with internal nuclei in Tibialis anterior cross-sections. Results are mean ± SEM; n = 3 mice per group; *p < 0.05; Chi-square test. Results are mean ± SEM; n = 3 mice per group. (I) Creatine kinase (CK) activity in the sera from control and SH3BP2 mKO mice. Results are mean ± SEM; n = 4 mice per group; p > 0,05; two-tailed Student’s t-test.

Reduced muscle strength and animal fitness could be the cause of muscle atrophy; therefore, we performed histological analysis of Tibialis muscle cross-sections. General fiber morphology appeared normal in SH3BP2 mKO, and quantification of muscle fiber area revealed no significant differences compared to control (**Fig. 8, F and G**). Muscles undergoing degeneration and regeneration cycles have nuclei that migrate from the periphery of the fibers to their central regions. In SH3BP2 mKO muscle cross-sections, we did not observe any typically centralized nuclei, and the fraction of nuclei within the fibers, which we classified as internal, was not significantly different from control animals (**Fig. 8, F, and H**). Damaged muscle fibers release creatine kinase, increasing this protein’s levels in blood serum. However, SH3BP2 mKO mice have levels of creatine kinase in serum comparable to controls (**Fig. 8, I**). Collectively, our data suggest that SH3BP2 deletion in skeletal muscles is not associated with muscle degeneration or injury and argue for the neuromuscular transmission defects as a primary cause of muscle weakness.

### Impaired neuromuscular transmission in SH3BP2 mKO mice

To study neuromuscular transmission, we performed experiments in which we electrically stimulated the Tibialis anterior muscle (**Fig. 9, A**) or the sciatic nerve (**Fig. 9, E**) and measured the force generated by the hind paw of anesthetized mice. We found that the force generated by SH3BP2 mKO mice in response to single muscle stimulation at 5 mA for 0.2 ms was slightly decreased compared to SH3BP2 fl/fl (control) animals (**Fig. 9, B**). When the Tibialis muscle was repetitively stimulated at 50, 100, and 125 Hz, the generated force in tetanic contraction was similar in control and mutant mice, and only at 50 Hz the differences reached statistical significance (**Fig. 9, C and D**). However, in a single twitch experiment, sciatic nerve stimulation revealed a much more substantial force decrement in SH3BP2 mKO mice (**Fig. 9, F**). Also, repetitive nerve stimulation of the sciatic nerve showed that the tetanic force generated by SH3BP3 mKO mice was much weaker than that caused by control animals (**Fig. 9, G, and H**). The observed differences were statistically significant for all stimulation frequencies in this case. Collectively, our results suggest that SH3BP2 mKO mice may have some defects in the muscle contraction mechanisms and compromised neurotransmission at the NMJ.

**Fig. 9.**
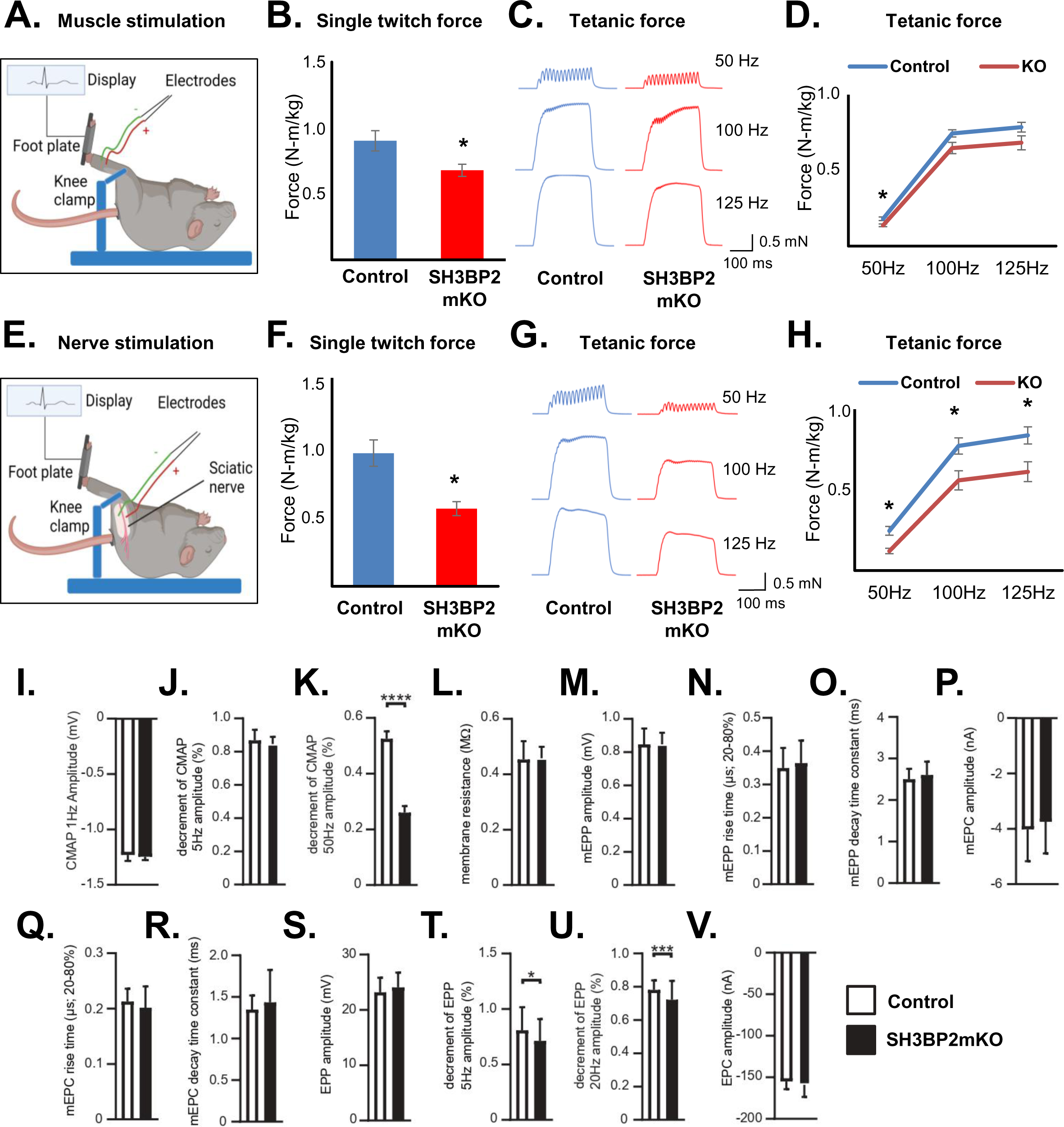
Impaired neuromuscular transmission in SH3BP2 mKO mice. (A) Schematic diagram of experiment with the single twitch and tetanic force measurement in response to direct muscle stimulation. (B) The single twitch force generated by control and SH3BP2 mKO mice in response to muscle stimulation at 5 mA for 0.2 ms. (C) Representative tetanic curves in response to direct muscle stimulation at 50, 100, and 125 Hz frequencies. (D) Quantification of tetanic force in response to direct muscle stimulation. All results are mean ± SEM; n = 5 mice per group; *p < 0.05, unpaired t-test. (E) Schematic diagram of experiment with the twitch and tetanic force measurements in response to nerve stimulation. (F) A significant reduction in single twitch force was observed in SH3BP2 mKO mice in response to sciatic nerve stimulation. (G) Representative tetanic curves in response to repetitive nerve stimulation at 50, 100, and 125 Hz frequencies. (H) In response to sciatic nerve stimulation, a significant reduction in generated tetanic force was observed in SH3BP2 mKO mice. All results are mean ± SEM; n = 5 mice per group; *p < 0.05, unpaired t-test. (I-K) CMAP analysis of diaphragm muscles revealed significant decrement at 50 Hz stimulation in SH3BP2 mKO mice. (L) Muscle fiber membrane resistance was normal without SH3BP2. mEPP amplitude (M), rise time (N), and decay time constant (O) were also normal in SH3BPO2 mKO mice. Similarly, mEPC amplitude (P), rise time (Q), and decay time constant (R) were also normal in mutant mice. EPP amplitude was unaltered in a single stimulation experiment (S), but repetitive stimulation at 5 Hz (T) and 20 Hz (U) revealed amplitude decrement in SH3BP2 mKO mice. Results are mean ± SEM; n=5 mice per group; ****p < 0.0001, ***p < 0.001, *p < 0.05; unpaired t test.

To monitor the mechanical characteristics of neurotransmission defects in SH3BP2 mKO mice, we performed electrophysiological recordings on diaphragm muscles ex vivo. The compound muscle action potential (CMAP) in response to nerve stimulation at 1 and 5 Hz stimulation was comparable between WT and mutant diaphragms (**Fig. 9, I, J**). However, a substantial decrease of the decrement in CMAP was observed at 50 Hz stimulation in mutant mice (**Fig. 9, K**). Next, we performed single fiber recordings. We first checked the resistance of the muscle fiber membrane, which was normal in mutant mice (**Fig. 9, L**), suggesting that muscle fiber sarcolemma are intact. To characterize neuromuscular transmission, we measured miniature endplate potentials (mEPPs), which are local depolarizations of the sarcolemma around endplates in response to spontaneous ACh release^56,57^. mEPP amplitudes, rise time, and decay time constant in SH3BP2 mKO mice were comparable with those in control mice (**Fig. 9 M, N, and O**), suggesting functional AChR channel characteristics in SH3BP2 mKO mice. Then we analyzed miniature endplate currents (mEPC), but their amplitudes, rise time, and decay time constant were also normal in SH3BP2 mKO mice (**Fig. 9, P, Q, and R**). Endplate potential (EPP) and endplate current (EPC) were also normal in mutant mice (**Fig. 9, S, V**). However, repetitive stimulation at 5 Hz and 20 Hz revealed a significant decrease of decrement of EPP in SH3BP2 mKO mice, suggesting increased fatigability (**Fig. 9, T, U**). These defects might relate to structural NMJ impairments as reflected by an increased NMJ fragmentation^58^ (**Fig. 7, Fig. S3, and Fig. S4**).

## Discussion

### Summary of results

Our previous work identified a scaffold protein SH3BP2 as a potential interactor of αDB1 recruited to the phosphorylated tyrosine Y713^23^. SH3BP2 has been shown to regulate signal transduction in bone cells and be involved in the pathology of Cherubism^35,59^. Still, its localization and function in skeletal muscles have not been investigated. This study demonstrates that SH3BP2 binds directly to phosphorylated αDB1 and that the SH2 domain mediates this interaction. Immunohistochemical analysis and muscle electroporation experiments confirmed that SH3BP2 localizes to the NMJ postsynaptic domain and, similarly to αDB1, co-localizes with AChR^23^. Our BiFC experiments suggest that the interaction of SH3BP2 with αDB1 occurs at the NMJ, where αDB1 has been shown to concentrate^23^. Unexpectedly, the SH2 domain and αDB1 appeared dispensable for synaptic localization of SH3BP2. Moreover, each of the three SH3BP2 domains targeted GFP to the NMJ. These results suggested that all three SH3BP2 domains interact with the postsynaptic components. Our proteomics-based identification of SH3BP2-interacting proteins from muscle cell extract identified four DGC components. We provide evidence that SH3BP2 interacts with syntrophin-1, β-dystroglycan, and αDB1. Therefore, SH3BP2 exhibits polyvalent interactions with the DGC. Additionally, our proteomics experiment and subsequent co-immunoprecipitations identified that SH3BP2 binds AChR subunits α and γ and that at least interaction with AChRα is direct. Interestingly, our domain mapping experiments revealed that all three SH3BP2 domains can interact with AChRα at three different sites. This observation is consistent with our imaging data, showing that all three SH3BP2 domains are recruited to the NMJ. Thus, SH3BP2 exhibits polyvalent binding also to the AChR pentamers, which contain two copies of the α subunit. Proteins that exhibit polyvalent interactions can form a network of interactions inducing liquid-liquid domain separation^16^. This phenomenon is often mediated by intrinsically disordered protein regions^47^. We demonstrated that SH3BP2 can interact with itself, triggering phase separation, and that this process requires the ID region of SH3BP2. Protein condensates form a unique microenvironment, allowing for interactions with additional proteins. As expected, phase-separating SH3BP2 was able to recruit the cytoplasmic domain of AChRα. This observation suggests that SH3BP2 may cluster the postsynaptic AChR receptors. We demonstrated that SH3BP2 targeted to the cell periphery can cluster AChR in Hek293 cells and that this process depends on the ID region. Consistently, C2C12 myotubes depleted of SH3BP2 failed to cluster AChR, although their plasma membrane expression was normal. Mice with muscle-specific SH3BP2 deletion have increased NMJ fragmentation and misalignment of AChR with AChE, suggesting the removal of receptors from the synaptic regions. Also, ultrastructural analysis revealed multiple alterations in the NMJ organization, e.g., increased number, length, and width of synaptic folds and synaptic cleft. Similar results, i.e., increased NMJ fragmentation, were observed when SH3BP2 deletion was achieved by infecting muscle with AAV-CRE. Subsequent experiments with twitch force analysis after electrical stimulation showed that SH3BP2 mKO mice have defects in muscle ability to generate force in response to direct stimulation and impaired neuromuscular transmission. Electrophysiological recordings confirmed this observation, revealing defects in CMAP and EPP at higher stimulation rates. On the other hand, our behavioral analysis demonstrated that SH3BP2 mKO mice have reduced grip strength and ability to run and increased fatigue. At the same time, general muscle histology has not revealed signs of muscular atrophy, and blood levels of the creatine kinase were normal in mutant mice. Similarly, the deletion of SH3BP2 with AAV-Cre caused impaired muscle strength and increased fatigability. All these results point to important SH3BP2 functions in the neuromuscular system.

### Interaction of SH3BP2 and DGC

Our study demonstrates that SH3BP2 interacts with several DGC components, which serve as a scaffold for stabilizing the postsynaptic components, but the underlying molecular mechanisms are not fully understood^8^. Similarly, αDB1 and its phosphorylation have been shown to play an important role in stabilizing AChR, but again, the underlying mechanisms remained unclear^23,24^. The DGC and αDB1 provide a platform for assembling the cortical molecular machinery involved in signal transduction and cytoskeleton organization^8,60^. In this study, we have shown that SH3BP2 is peripherally recruited to the DGC. Lack of functions in several integral DGC components leads to muscular dystrophy, e.g., dystroglycan, dystrophin, and sarcoglycan^6,61–64^. Without those components, the entire DGC falls apart. On the other hand, the lack of function for other DGC proteins affects the NMJ without showing the symptoms of muscular dystrophy. For example, knockout of syntrophin-1 or utrophin does not cause muscular dystrophy^65,66^. Similarly, lack of SH3BP2 does not cause symptoms of muscular dystrophy. On the other hand, αDB1 knock out mice showed unstable and patchy distribution of AChR with 50% of dystrophic fibres^24,25^. αDB1 recruits many proteins regulating the postsynaptic organization, and SH3BP2 appears to have a role in AChR clustering but not in maintaining general fiber integrity. However, scaffold proteins often have redundant functions, and we cannot exclude that some of the SH3BP2 molecular functions are compensated by other scaffold proteins, e.g., Grb2 that interacts with SH3BP2, αDB1, dystroglycan, and AChR^23,67,68^ or SH2B1 that has been identified as αDB1-binding partners, but its role at the NMJ awaits investigation^23^. The lack of functions in several DGC components affects synaptic folds formation and organization. For instance, deletion of utrophin or αDB1 or mutations in dystrophin caused reduced synaptic folds^62,65,69^. On the other hand, SH3BP2 mKO mice have increased length and number of synaptic folds. The lack of SH3BP2 also leads to the mislocalization of electron-dense postsynaptic machinery from the crests of folds to patches distributed along the exuberant folds. Such phenotype was observed in mice lacking αDB1 or with reduced rapsyn expression and was interpreted as compensatory mechanisms for mutants with compromised AChR clustering^70,10^. This is in agreement with our observations that SH3BP2 is involved in the clustering of AChR similar to rapsyn, although to a lesser extent.

Rapsyn binds to the groove between AChRα and AChRβ and also with β-dystroglycan^5,48,71^. This interaction suggests that rapsyn can connect AChR to DGC complexes. Similarly, we demonstrated that SH3BP2 interacted with the cytoplasmic loop of AChRα and AChRγ. SH3BP2 also interacts with β-dystroglycan, suggesting it may connect AChR to DGC complexes to stabilize AChR. Rapsyn is the key molecule that clusters AChR, and the lack of rapsyn in mice leads to death at birth due to the complete absence of the postsynaptic machinery. In contrast, mice with SH3BP2 deletion are viable and exhibit relatively mild postsynaptic defects. This strongly argues that rapsyn has much more critical functions in the organization of the AChR than the SH3BP2. Rapsyn is targeted to the plasma membrane by myristylation, and seven TPR motifs promote rapsyn oligomerization that provides a scaffold for AChR clustering^14,16,72^. Rapsyn clustering activity has also been shown to depend on the E3 ligase activity, and a mutation in the cysteine residue impairs its E3 ligase activity and AChRs clustering. This suggests that rapsyn can also act as a signaling molecule^17^. Additional functions of rapsyn involve interactions with actin and intermediate cytoskeleton and with the cortical microtubules^18–20,73,74^. Therefore, rapsyn has many functions at the NMJ besides aggregating AChR into clusters. SH3BP2, on the other hand, has only three protein domains apart from its intrinsically disordered sequences.

Recent studies demonstrated that rapsyn oligomerization triggers liquid-liquid phase separation and that this phenomenon regulates AChR aggregation^16^. Phase separation is a new concept that provides fundamental insight into the biophysical principles regulating the assembly of the postsynaptic machinery in the central and peripheral nervous systems^75,76^ SH3BP2 is a second example of a protein in the muscle postsynaptic machinery that triggers phase separation. Our findings strengthen the proposed mechanisms that the NMJ postsynaptic machinery formation involves phase separation. This opens a perspective for further investigation and enhances our understanding of mechanisms regulating synaptogenesis in the nervous system.

## MATERIALS AND METHODS

### Ethics statement

Procedures using animals were conducted in compliance with the ethics committee guidelines for animal experimentation and regulations at the Łukasiewicz–PORT, the Nencki Institute of Experimental Biology, Poland (permissions 709/2015, 723/2015, 369/2017, 065/2020/P1, 066/2020/P1, 085/2021/DZ/NZP/P1, 055/2023, 058/2023/NZP), and the Friedrich-Alexander University Erlangen-Nürnberg, Germany (permissions I/39/EE006, TS-07/11).

### Animals

All the mice used in this study had a C57/BL6 genetic background. Myf5-Cre mice were a kind gift from Markus Ruegg (University of Basel, Switzerland). SH3BP2^loxP^ mice (Sh3bp2<tm1c(KOMP)Wtsi>) with the loxP sites flanking the exons 6 and 7 of the *SH3BP2* gene were from Jackson laboratory^77^ and were crossed with Myf5-Cre mice to generate muscle-specific knockout of SH3BP2 (SH3BP2 mKO). The generation of STOP-Tom reporter mice was described before^78^. This study used both male and female mice. The animals were kept in the animal house on a 12-hour light/dark cycle with free access to food and water. Before experiments, animals were habituated to the experimentator. Mice were euthanized by cervical dislocation in accordance with the national and institute’s ethical guidelines. Depending on the experiments, Tibialis anterior, soleus, and diaphragm muscles were collected for cryosectioning or Western blot experiments. For electrophysiology experiments, muscles were placed into a physiological solution immediately after dissection. For the NMJ analysis, transcardial perfusion was performed with phosphate-buffered saline (PBS, pH 7.4) followed by 4% paraformaldehyde (PFA, pH 7.4) to fix tissues. This procedure was performed on mice anesthetized with ketamine/xylazine or immediately after cervical dislocation.

### Antibodies and staining

All primary antibodies used in this study are shown in **Supplementary Table 2**. For immunofluorescence studies, the following secondary antibodies were used: goat anti-rabbit IgG (H+L) coupled to Alexa Fluor 488 (A-11034, Thermo Fisher Scientific), Alexa Fluor 546 (A-11035, Thermo Fisher Scientific) or Alexa Fluor 568 (A-11011, Thermo Fisher Scientific); goat anti-mouse IgG (H+L) conjugated with Alexa Fluor 488 (A28175, Thermo Fisher Scientific) or Alexa Fluor 568 (A-11004, Thermo Fisher Scientific). The secondary antibodies for Western blotting were HRP-coupled anti-IgG polyclonal antibodies (Cell Signaling). To visualize the AChR in the myofibers, myotubes, or Hek293 cells, we used fluorescently labeled α-bungarotoxin (B35451 or B13422; Thermo Fisher Scientific). Alexa Fluor 546 Phalloidin (A22283, Thermo Fisher Scientific) was used to visualize F-actin, and fasciculin II (F-225; Alomone Labs, labeled with FluoReporter FITC kit from Thermo Fisher Scientific) was used for AChE labeling.

### Plasmids and siRNA

The EGFP–αDB1 (mouse) plasmid and CMV-Rapsyn-mCherry plasmid were described previously^24,73^. The plasmids encoding the full-length α1, β1, δ, and ε nAChR subunits were a kind gift from Dr. Lin Mei (Case Western Reserve University; USA). SH3BP2 (mouse) and its various domains (PH, PR, and SH2 domains) were sub-cloned into a modified pCDNA3.1 backbone containing a FLAG or GFP tag. SH3BP2 was also cloned into pBiFC-VC155^80^, a gift from Chang-Deng Hu (Addgene #22011), and αDB1 was cloned into pBiFC-VN173^80^, a gift from Chang-Deng Hu (Addgene #22010). The full-length syntrophin-1 (mouse), a cytoplasmic loop of AChRα (mouse; amino acid 317-428), a cytoplasmic loop of AChRγ (mouse; amino acid 330-476), the cytoplasmic domain of β-dystroglycan (mouse; amino acid 773-893) were cloned into EGFP-N2 plasmid. A membrane localization signal from the Lyn protein (amino acid sequence: MGCIKSKGKDS) was added to the GFP-tagged SH3BP2 and subcloned into the pCDNA3.1 backbone. The SH3BP2 without the intrinsic disorder domain (amino acid 164-449) with the membrane localization sequence and the GFP tag was subcloned into the pCDNA3.1. For protein purification from bacteria, we cloned the cytoplasmic domain of AChRα (mouse; amino acid 317-428) with or without mCherry into pGEX-4T1 (Cytiva 28-9545-49; Merck). GFP-SH3BP2 with or without the intrinsic disorder domain (amino acid 164-449) into pEBG-GST^81^, a gift from David Baltimore (Addgene #22227). Knockdown experiments were performed with the following siRNAs: siRNA1 (SI02739478, Qiagen), siRNA2 (SI02692193, Qiagen), siRNA3 (SI00204218, Qiagen) targeting SH3BP2 and non-targeting control siRNA (12935300, Invitrogen).

### Protein complex purification and mass spectrometry analysis

For identification of SH3BP2-interacting proteins, Hek293 cells overexpressing TAP-GFP-SH3BP2 were lysed in lysis buffer (50 mM Tris-HCl [pH 8.0], 150 mM NaCl, 1% Igepal CA-630 [NP-40], 0.5% sodium dodecyl sulfate [SDS], 10% glycerol, 1 mM dithiothreitol [DTT], 1 mM NaF) supplemented with Mini Protease Inhibitors cocktail (11873580001, Roche) and incubated with Dynabeads M-270 Epoxy (14311D, Invitrogen) coated with Rabbit IgG (I5006-10MG, SIGMA) overnight with rotation at 4°C. The next day, the beads were washed four times in wash buffer (50 mM Tris-HCl [pH 8.0], 500 mM NaCl, 1% NP-40, 0.5% SDS, 1 mM DTT, 1 mM NaF, and Mini Protease Inhibitors cocktail). The beads enriched with TAP-GFP-SH3BP2 were equilibrated with C2C12 cells lysis buffer (50 mM Tris-HCl [pH 8.0], 150 mM NaCl, 0.1% NP-40, 10% glycerol, 1 mM DTT, 1 mM NaF, and Mini Protease Inhibitors cocktail) and incubated with C2C12 myotubes lysates overnight at 4°C with rotation. Next day, the beads were washed three times with C2C12 cells wash buffer (50 mM Tris-HCl [pH 8.0], 150 mM NaCl, 0.1% NP-40, 1 mM DTT, 1 mM NaF, and Mini Protease Inhibitors cocktail), equilibrated with the TEV buffer (50 mM Tris-HCl [pH 8.0], 150 mM NaCl, 0.5% NP-40, 0.5 mM EDTA, 1 mM DTT, and 1 mM NaF) and proteins were eluted by incubating the beads in TEV buffer containing TEV protease (T4455, Sigma Aldrich) overnight in 4°C. The eluted proteins were precipitated with Trichloroacetic acid, and the pellet was washed with ice-cold acetone. A part of the samples was analyzed by SDS-PAGE electrophoretic separation on polyacrylamide gels followed by silver staining, and the rest of the samples were sent to the MS Bioworks facility (USA) for protein identification.

### Immunoprecipitation

Synthetic biotinylated proteins were attached to Streptavidin coupled Dynabeads M-280 (11205D, Life Technologies) in binding buffer (50 mM Tris-HCl, 150 mM NaCl, 50 mM NaH_2_PO_4_, 10mM imidazole, 0.1% NP40, 10% glycerol, 10mM β-mercaptoethanol, EDTA-free mini protease inhibitor cocktail (Roche), 1 mM phenylmethylsulphonyl fluoride, pervanadate, pH 8.0). After three washes with the same buffer, beads were incubated with purified proteins in the binding buffer or with an extract from Hek293 cells lysed with the binding buffer. After incubation, beads were washed three times with the binding buffer, and proteins were eluted in 2x Laemmli sample buffer with 50 mM DTT by boiling at 95°C for 10 min. For coimmunoprecipitation experiments, Protein G Dynabeads (G 10004D, Invitrogen) were coated with antibodies indicated on the figures according to the manufacturer’s protocol. Beads were incubated overnight at 4°C with extracts of Hek293 cells expressing indicated proteins. After three washes with the lysis buffer, proteins were eluted in 2x Laemmli sample buffer with 50 mM DTT by boiling at 95°C for 10 min. The precipitation of the surface and total AChR from the myotubes were performed as described previously^60^. For the precipitation of surface AChR, the myotubes were incubated with BTX-biotin (B1196, Invitrogen) for 5 minutes at 37°C and washed three times with PBS, followed by lysis of myotubes in lysis buffer (100 mM Hepes pH 7.4, 150 mM NaCl, 1 mM PMSF, 0.5% NP-40, 1% CHAPS). For precipitation of total AChR, myotubes were lysed in lysis buffer and incubated with BTX-biotin for 30 minutes at 4°C. NeutrAvidin beads (29200, Thermo Fisher Scientific) were incubated with lysates overnight at 4°C. The beads were washed three times in lysis buffer, and proteins were eluted in 2x Laemmli sample buffer by boiling for 10 min.

### Western blot

For Western blot, the lysates or protein precipitates were mixed with the sample buffer, boiled for 10 min, resolved by SDS-PAGE electrophoresis, and transferred to nitrocellulose membranes (66485, Pall Corporation) using Trans-Blot Turbo (1704270, Bio-Rad) following manufacturer’s recommendation. Membranes were incubated with blocking buffer containing 5% milk in TBST (20 mM Tris-HCl, 150 mM NaCl [pH 7.6], and 0.1% Tween20) for 1h, and incubated with their respective antibody in the blocking buffer overnight at 4°C. The membranes were washed with TBST and incubated with appropriate secondary antibodies conjugated to horseradish peroxidase. The membranes were washed with TBST, developed with Clarity Chemiluminescent Substrate (1705060, Bio-Rad), imaged with photographic film (CL-XPosure, Thermo), and digitized using a scanner (Epson), or were digitally acquired on the Biorad ChemiDoc XRS Imaging System.

### Cell culture and transfection

C2C12 myoblasts were cultured and differentiated as described previously^82^. Briefly, C2C12 cells (91031101, Sigma) were cultured on 0.2% gelatin-coated plates in Dulbecco’s modified Eagle’s medium (670087, DMEM, Gibco™) containing 20% fetal bovine serum supplemented with penicillin, streptomycin, and fungizone. Then, cells were differentiated on Petri dishes or 8-well Permanox chamber slides (177445, Sigma-Aldrich) precoated with 10 μg/ml solution of laminin 111 (23017–015, Invitrogen) in differentiation medium (DMEM supplemented with 2% horse serum (26050088, Thermo Fisher Scientific). For agrin induced AChR clustering experiment, 8-well Permanox chamber slides were coated with 0.2% gelatin instead of laminin, and myotubes were treated with 10 nM recombinant rat Z’-spliced agrin (550-AG-100/CF, Biokom) for 12 h. For fluorescence AChR cluster analysis, differentiated cells were fixed for 7 min in 4% PFA at room temperature, washed in PBS, and blocked with 2% goat serum and 0.5% Triton X-100 for 30 min. The myotubes were stained using Alexa Fluor 555-coupled bugarotoxin (B35451, Thermo Fisher Scientific) and Alexa Fluor 633 Phalloidin (A22284, Thermo Fisher Scientific) for F-actin. Cells were imaged on the Zeiss Cell Observer spinning disk confocal microscope, and images were processed using Fiji. For Western blot analysis and protein precipitation experiments, differentiated myotubes were lysed with ice-cold lysis buffer (50 mM Tris-HCl, 150 mM NaCl, 50 mM NaH_2_PO_4_, 10 mM imidazole, 0.1% NP40, 10% glycerol, protease inhibitor cocktail, and 50 mM DTT, pH 8.0), and the lysis suspension was incubated on a rotary shaker for 1 h at 4°C and centrifuged at 13,000 x g at 4°C. The supernatant was stored at −20°C. For qRT-PCR, cells were treated with Trisure reagent (BIO-38032, Bioline), and RNA was isolated using the manufacturer’s recommendation. The knockdown of SH3BP2 in C2C12 myotubes was performed following the procedures described previously^23^. Briefly, cells were transfected with siRNA targeting SH3BP2 or non-targeting control siRNA using Lipofectamine RNAiMAX (13778075, Invitrogen) on day three after differentiation induction following the manufacturer’s recommendation. On day five after differentiation induction, cells were fixed with 4% PFA or treated with Trisure reagent (BIO-38032, Bioline) for qRT-PCR analysis or lysed with the Lysis buffer for Western blot analysis or coprecipitation experiments.

Hek293 cells (CRL-1573, ATCC) were cultured in DMEM (670087, Gibco™) containing 10% fetal bovine serum supplemented with penicillin, streptomycin, and Fungizone. Transient transfections were performed with Lipofectamine 2000 (11668019, Invitrogen), according to the manufacturer’s instructions. Hek293 cells were lysed using the same method as C2C12 cells.

### RNA extraction, reverse transcription, and PCR

The total RNA was extracted from myotubes or Tibialis muscles using Trisure Reagent (BIO-38032, Bioline) following the manufacturer’s recommendations. About 1 μg of RNA was used for cDNA preparation using the High-Capacity cDNAReverse Transcription Kit (4374966, Applied Biosystems). qRT-PCR was performed using Fast SYBR Green Master Mix (4309155, Thermo Fisher Scientific) on the Step One Plus instrument (Thermo Fisher Scientific). For the qRT-PCR, the following primers were used: 5′-GAAGACTATGAGAAGGTGCCG -3′ and 5′-AAGATTCATCCCACACGACC-3′ specific to SH3BP2, and 5′-GGCCTTCCGTGTTCCTAC-3′ and 5′-TGTCATCATACTTGGCAGGTT-3′ specific to GAPDH.

### Immunofluorescence microscopy

For immunostaining of muscle fibers, mice were perfused with 4% PFA, and isolated dissociated muscle fibers of the Tibialis anterior (TA), soleus, and diaphragm muscles were incubated with blocking buffer (2% BSA and 2% goat serum in PBS with 0.1% Triton X-100) for 30 min. The fibers were then incubated with primary antibodies in PBS with 0.1% Tween 20 (PBST) at 4°C overnight, followed by three washes in PBST. Next, tissues were incubated for 1 h at room temperature with fluorescent secondary antibodies and BTX–A555 (B35451, Life Technologies) for 30 min. Coverslips were mounted on slides with FluorSave reagent (Calbiochem) or Fluoromount mounting medium (F4680, Merck). For the Tibialis tissue cross-section, the images were acquired with a spinning disk confocal microscope (Zeiss). Images were processed with Fiji software.

### In vivo electroporation and AAV infection

For electroporation of the Tibialis anterior muscle mice were anesthetized by isoflurane inhalation, and 30 μg DNA plasmid (1 μg/μL in PBS) was injected into the muscle with a Hamilton syringe. The muscle was electroporated with 10 pulses of electrical current (150 V/cm) of 20 ms each at 1 s intervals using an ECM-830 electroporator (45-0052INT, BTX Harvard Apparatus). Lidocaine gel was applied on the skin after electroporation and intraperitoneal injection of Metacam 0.4 ml/kg was performed twice daily for three days and. Mice were sacrificed 7 days after the electroporation. AAV9-CRE-GFP particles expressing CRE and GFP (SL100842) and AAV9-GFP particles expressing GFP (SL100840) were purchased from SignaGen Laboratories. For injection at P0, pups were placed on ice for 5 min, and Tibialis anterior muscles were injected with 3 µL (1×10^12^ vg/mL) of AAVs. For injection at P60, mice were anesthetized with isoflurane inhalation, and Tibialis muscles were injected with 10 μL (1×10^12^ vg/mL) of AAVs. Lidocaine gel was applied on the skin in the place of injection.

### Synthetic Peptides and Recombinant Proteins

The synthetic peptides corresponding to αDB1 sequence (Y713: DEST–Biotin–Ahx– TQPEDGNYENESVRQ-NH2; Y713-P: DEST–Biotin–Ahx–TQPEDGNpYENESVRQ-NH2; Y730: DEST–Biotin– Ahx–RQLENELQLEEYLKQKLQDE-NH2; and Y730P: DEST– Biotin–Ahx–RQLENELQLEEpYLKQKLQDE-NH2; where NH2 denotes peptide amidation, ‘p’ indicates phosphorylation of the tyrosine residue, and Ahx indicates a linker) were purchased from Lifetein. The peptides covering the cytoplasmic domain of AChRα (P1: Biotin-NTHHRSPSTHIMPEWVRKVFIDT; P2: Biotin-VFIDTIPNIMFFSTMKRPSRDKQ; P3: Biotin-SRDKQEKRIFTEDIDISDISGKP; P4: Biotin-ISGKPGPPPMGFHSPLIKHPEVK; P5: Biotin-HPEVKSAIEGVKYIAETMKSDQE; P6: Biotin-KSDQESNNAAEEWKYVAMVMDH) were purchased from Lifetein, USA. The GST-cytAChRα and GST-mCherry-cytAChRα were purified from BL21 cells. Transformed BL21 E. coli were grown in 200 ml of LB medium at 18°C. The cells at an optical density between 0.4 and 0.6 were induced with 0.5 mM isopropyl 1-thio-β-galactopyranoside (IPTG) for 4 h at 18°C. The bacteria were collected and lysed in high salt buffer (500 mM NaCl, 25 mM Tris, pH 7.4, 5 mM DTT supplemented with protease inhibitors (11873580001, Sigma) and phosphatase inhibitors (4906845001, Sigma)) by sonication for 1 min. The extract was then centrifuged at 10,000 g, and the supernatant was collected. About 30 μL of equilibrated glutathione beads (17-0756-01, GE Healthcare) were incubated with the supernatant overnight at 4°C. After three washes with high salt buffer, proteins were eluted in elution buffer (10 mM glutathione, 500 mM NaCl, 25 mM Tris, pH 7.4). GST-mGFP-SH3BP2-FLAG and GST-mGFP-SH3BP2-FLAG without intrinsic disorder domain (GFP-ΔIDSH3BP2) were expressed and purified from Hek293 cell as described before^16^. Cells were resuspended in a high salt buffer (500 mM NaCl, 25 mM Tris, pH 7.4, 5 mM DTT supplemented with protease inhibitors (11873580001, Sigma) and phosphatase inhibitors (4906845001, Sigma)) 72 h after transfection, and the GST-tagged proteins were purified using Glutathione beads (as described above for purification from bacteria). GST used as a control was labeled with Alexa Fluor 594 Microscale Protein Labeling Kit (A30008, Invitrogen) following the manufacturer’s recommendation. SH3BP2-Flag was expressed in Hek293 cell and purified using anti-FLAG agarose beads (A2220-5ML, Sigma-Aldrich) and eluted with FLAG peptide (F4799-4MG, Sigma-Aldrich) following manufacturer’s recommendation.

### Phase separation

The phase separation of SH3BP2 was performed as described before^16^. All the proteins were stored in a high salt buffer (500 mM NaCl, 25 mM Tris, pH 7.4, 5 mM DTT). Phase separation experiment was performed at room temperature in a buffer containing 125 mM NaCl, 25 mM Tris, pH 7.4, and 5 mM DTT. For analysis of protein recruitment to SH3BP2 droplets, 2.5 μM GST-mGFP-SH3BP2-FLAG was incubated with 2.5 μM GST-mCherry-cytAChRα or GST-Alexa Fluor 594. Mixed proteins were then incubated at room temperature for 5 min before microscopy. To visualize the droplets, the solution was placed over a clean slide, a spacer with paper tape was applied around the liquid, and the slide was covered with a cover slide without touching the solution. Images were collected by Zeiss Cell Observer spinning disk confocal microscope using a 60x oil objective.

### Sedimentation assay

GST-mGFP-SH3BP2-FLAG protein was stored in the high salt buffer containing 500 mM NaCl, 25 mM Tris, pH 7.4, and 5 mM DTT. The solution was centrifuged at 14,000 x g for 1 min before the experiment. The protein was diluted to 1 µM and 5 µM into low salt buffer (125 mM NaCl, 25 mM Tris, pH 7.4, 5 mM DTT) and incubated for 60 min at room temperature. Then, the samples were centrifuged at 14,000 x g for 15 min at room temperature. The supernatant was collected in a separate tube, and the pellet was washed once with a low-salt buffer. The supernatant and the pellet were then resolved by SDS-PAGE and analyzed by Safe blue (NBS-SB1L, NBS Biologicals) staining using the manufacturer’s recommendation. The intensities were quantified by Fiji software.

### FRAP

FRAP was performed as described before^16^. The GST-mGFP-SH3BP2-FLAG was reconstituted in a low salt buffer as described above, and samples were imaged after 5 min of incubation at room temperature. Images with 0.1% laser power were acquired before and after bleaching with 5% laser power for 100 ms. FRAP assay was performed on a Leica SP8 confocal microscope, and the image was analyzed using Fiji to calculate FRAP within the droplets.

### AChR Clustering assay

AChR clustering assay was performed as described previously^84^. Briefly, Hek293 cells were plated on glass coverslip in a six-well culture dish and co-transfected with four plasmids (1 μg of mixed DNA) encoding AChR subunits α, β, δ, and ε, and one of the following constructs mCherry-tagged rapsyn (500 ng), mGFP-SH3BP2 (500 ng), mem-mGFP-SH3BP2 (500 ng), or mem-mGFP-ΔIDSH3BP2 (500 ng). Cells were fixed the next day with 4% PFA and stained with BTX in the blocking buffer without detergent to detect surface AChR. The number of AChR puncta per cell was analyzed. Three independent experiments were performed for each experimental point.

### Muscle cross-sections and NMJ Analysis

Mice were sacrificed by cervical dislocation, and Tibialis anterior muscles were collected, embedded in OCT (05-9801, Bio Optica), frozen in isopentane cooled with liquid nitrogen, and stored at −80°C. Muscle 10 µm-thick cryosections were generated using Leica CM 3050 S cryostat, collected on Ultra Plus microscope slides (10149870, Thermo Fisher Scientific), dried, and stored at −20°C. For NMJ analysis, Tibialis anterior, soleus, and diaphragm muscles were dissected from mice perfused with 4% PFA. The teased muscle fiber bundles were stained with bungarotoxin to visualize AChR, Fascicullin II to visualize AChE, and with indicated antibodies. Images of 25 to 30 NMJs per mouse were analyzed using the “NMJ-morph” ImageJ plugin^50^.

### Creatine Kinase Activity

The blood was collected from the male mice and allowed to clot by incubating at room temperature for 1 h followed by centrifugation (10,000 rpm for 10 min. at 4°C), and serum was collected. Creatine kinase activity was analyzed with the Creatine Kinase Activity Assay Kit (MAK116-1KT, Sigma Aldrich) using the manufacturer’s recommendation.

### Grip Strength Test

Grip strength tests were performed as described before^87^. Mice were habituated with the experimenter for five consecutive days, and the grip strength was measured using a Grip Strength Meter (47200, Ugo Basile) for all four paws (SH3BP2 mKO) or for hind limbs (SH3BP2 fl/fl infected with AAV-Cre). Mice were allowed to grasp the grid connected to the meter, followed by gentle pulling until the grid was released, and the maximal force was recorded. The whole procedure was repeated ten times for each animal with an interval of 1 min between repetitions. The mean force from ten repetitions was normalized to body weight.

### Voluntary Running Wheel and Treadmill Experiment

Mice were placed in cages with upright running wheels (1800/50, Ugo Basile) for 12 h during the dark phase with free access to water and food. The distance run by each mouse was recorded. The performance on a treadmill determining physical endurance was performed as described before^89^. Briefly, mice were familiarized with the treadmill (57630, Ugo Basile) for three consecutive days. In the experiment, mice were placed on a treadmill and allowed to run at a 10 m/min starting speed for 5 min. After that, the speed was gradually increased by 1 m/min every minute until mice got exhausted, but not longer than 50 min. Exhaustion was defined as a decline in running for 15 seconds despite an aversive stimulus in the form of an air puff provided when mice stopped. The maximal speed, running time, and running distance were recorded at the end of the experiment.

### Muscle twitch and tetanic force measurement

The twitch and tetanic force measurements were performed as described before^87^, using an Aurora Scientific 1300A device. Mice were anesthetized with ketamine and xylazine and placed on a 37°C heating pad. The left knees of the mice were stabilized by pressing the knee with a clamp, and the feet were fixed onto the footplate connected to the transducer. The angle of the footplate was set at 17°. For muscle stimulation, the Tibialis anterior muscles were stimulated by placing the needle electrode subcutaneously and stimulation at 5 mA for 0.2 ms. Ten repetitions of stimulation and force recording were performed at 30 s intervals.

For the nerve stimulation, the sciatic nerve was exposed surgically and stimulated by two needle electrodes positioned in close proximity. For the single twitch experiment, the nerve was stimulated at 5 mA for 0.2 ms, and the generated force was recorded in 10 repetitions at 30-second intervals. For the tetanic force determination, muscle or sciatic nerve were stimulated at frequencies of 50, 100, and 125 Hz for 300 ms in ten repetitions at 2-minute intervals. The recorded force was normalized to body weight.

### Electrophysiological recording

The electrophysiological recording of the diaphragm-phrenic nerve preparations was performed as described before^90^. The diaphragm-phrenic nerve preparations were kept in Liley’s solution and gassed with 95% O_2_ and 5% CO_2_ at room temperature. At the beginning of the experiment, the compound muscle action potential (CMAP) was recorded using a micropipette with a tip diameter of approximately 10 µm filled with bathing solution. The electrode was then positioned 100 µm above the surface of the muscle, and CMAP was recorded. Intracellular recordings were performed in a nerve-independent (mEPPs/mEPC) or nerve-dependent (EPP/EPC) manner with electrodes introduced into single fibers. Two intracellular electrodes (resistance 10–15 MΩ) were inserted within 50 µm of the NMJs under visual inspection. At most NMJs, 50–100 spontaneous quantal events were recorded during 1 min. Decrements of EPPs were calculated employing the mean of the first and the last five recordings. pCLAMP 10 was used for recordings and analysis.

### Electron Microscopy

For electron microscopy, mice were perfused in 4% paraformaldehyde with 1% glutaraldehyde. Tibialis anterior muscles were collected and stained with bungarotoxin to visualize AChR. Small muscle specimens enriched in NMJs were dissected under a fluorescent microscope. Collected muscle fragments were washed with PBS, post-fixed with 2.5% OsO_4_, stained *en-bloc* with 2% uranium acetate, dehydrated in graded concentrations of ethanol, and embedded in Agar low viscosity resin (AGR1078, Agar Scientific). Semi-thin sections were cut using a glass knife, collected on glass slides, stained with 1% toluidine blue, and inspected with a light microscope to identify NMJs. Thin sections were cut from selected locations using a diamond knife and placed on copper grids for analysis using a STEM detector in a Zeiss Auriga SEM/FIB microscope with an accelerating voltage of 20 kV.

### Statistical analyses

GraphPad Prism version 7.0 was used to perform statistical analysis. Data were analyzed using a Student t-test to compare two groups. Unless otherwise indicated, data were given as mean ± SEM. For a multiple factorial analysis of variance, one-way or two-way ANOVA was applied. The significance was **** p < 0.0001, *** p < 0.001, ** p <0.01, or * p < 0.05, no stars for p ≥ 0.05.

### Data availability

All the data were provided as the supplementary dataset in the manuscript, and the reagents used in the manuscript will be made available upon considerable request.

## Acknowledgments

We are very grateful to Markus Ruegg from the University of Basel for providing us with Myf5-Cre mice. We would also like to thank Przemysław Kaczor and Margareta Jabłońska for their help with confocal microscopy and Paloma Alvarez-Suarez for her collaborative efforts. We want to thank Michał Malewicz and Mateusz Kucharczyk from Łukasiewicz-PORT for their careful reading of the manuscript and valuable comments. We would like to thank BioRender.com for the preparation of the diagram.

## Author Contributions

Conceptualization, K.B., B.S.P., and T.J.P.; methodology, K.B., B.S.P., Y.B., A.S., T.D.C., M.G., J.K., G.C., S.H., and T.J.P.; validation, B.S.P., S.H., and T.J.P.; formal analysis, B.S.P., M.G., K.B., A.S., T.D.C., S.H., and T.J.P.; investigation and data curation, B.S.P., K.B., M.G., Y.B., A.S., T.D.C., G.C., S.H., and T.J.P.; resources, T.J.P.; writing—original draft preparation, B.S.P.; writing—review and editing, B.S.P., S.H., and T.J.P.; supervision, T.J.P.; project administration, B.S.P. and T.J.P.. All authors have read and agreed to the published version of the manuscript.

---

**Supplementary Table 1.**
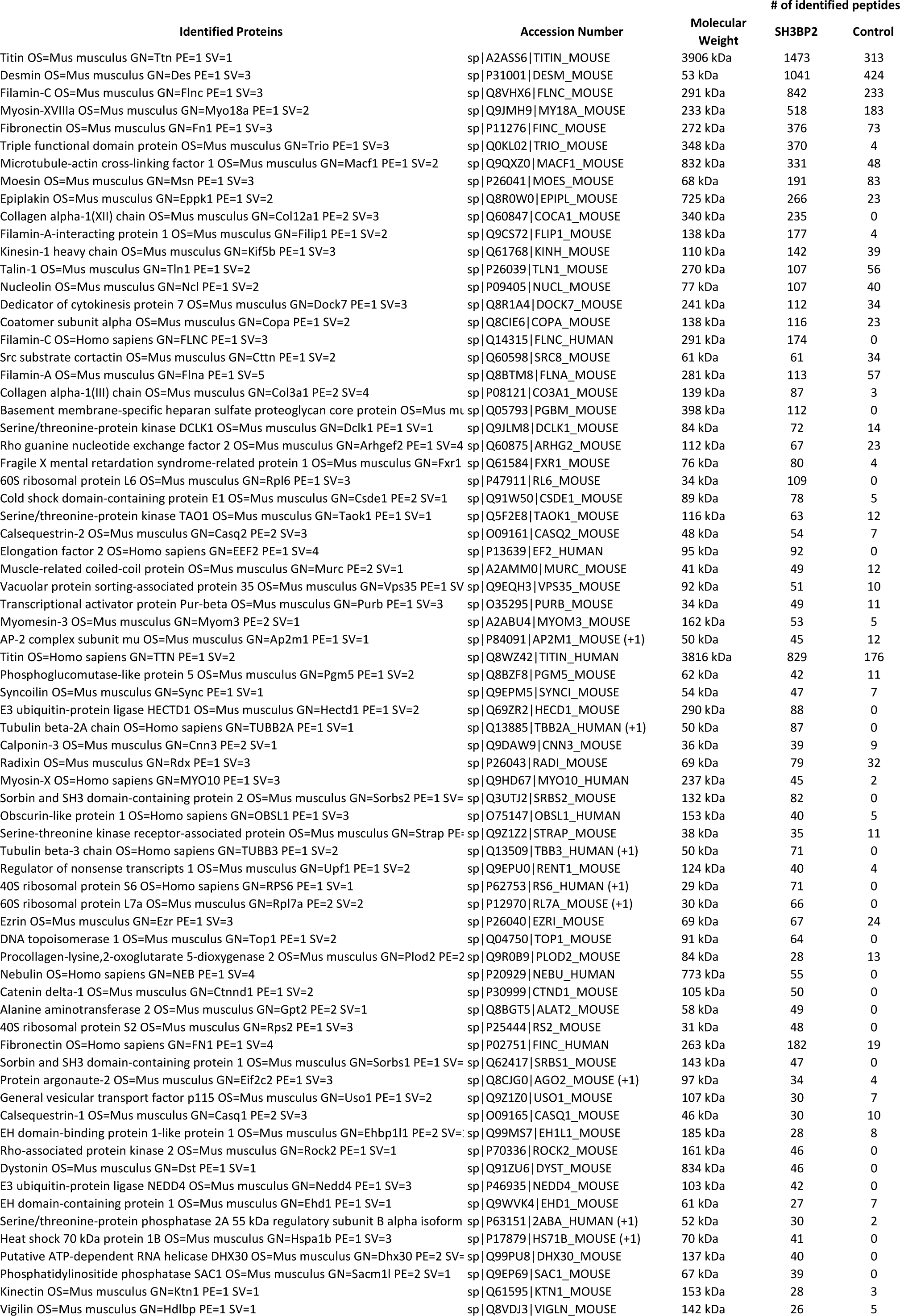
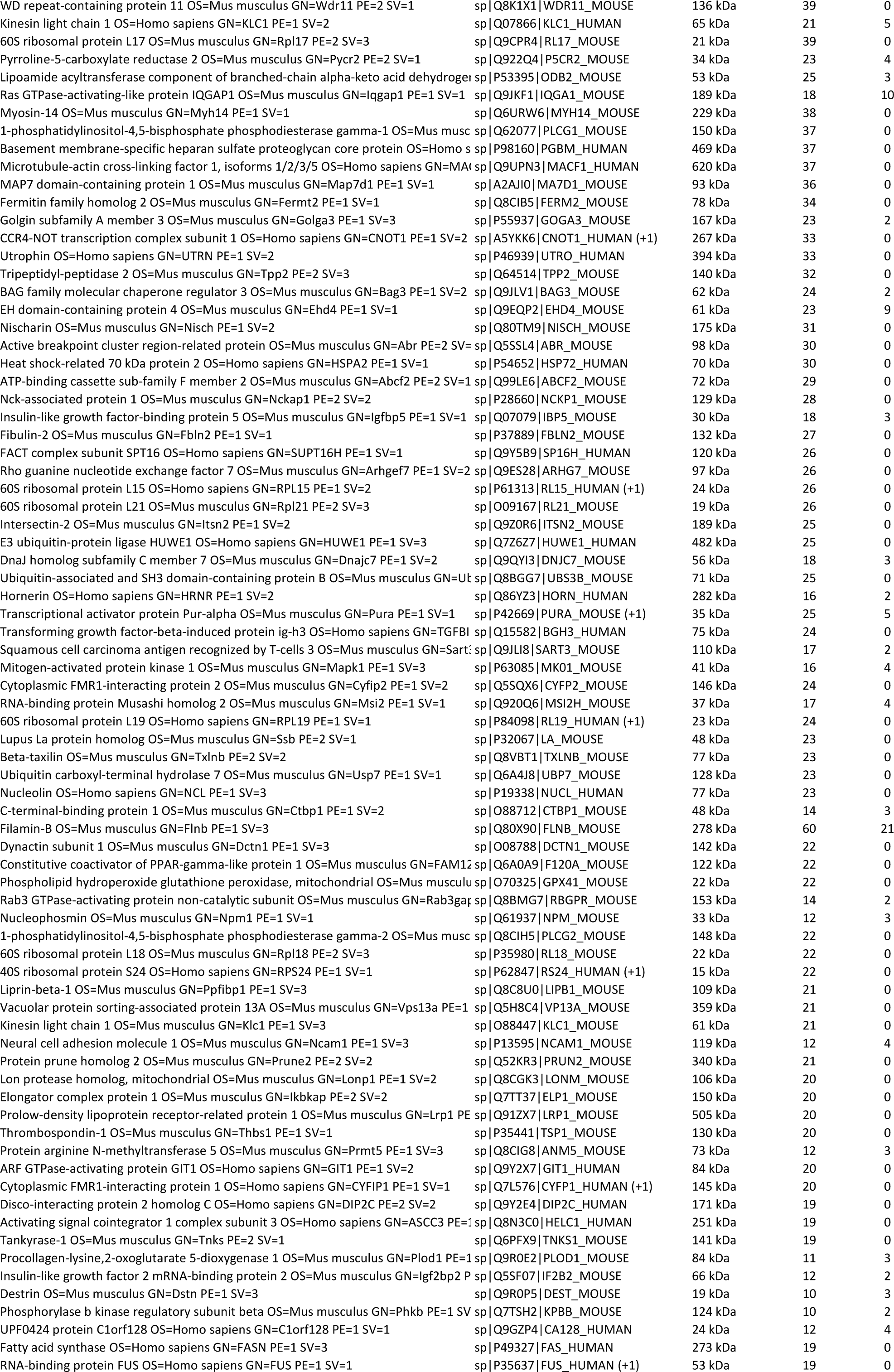
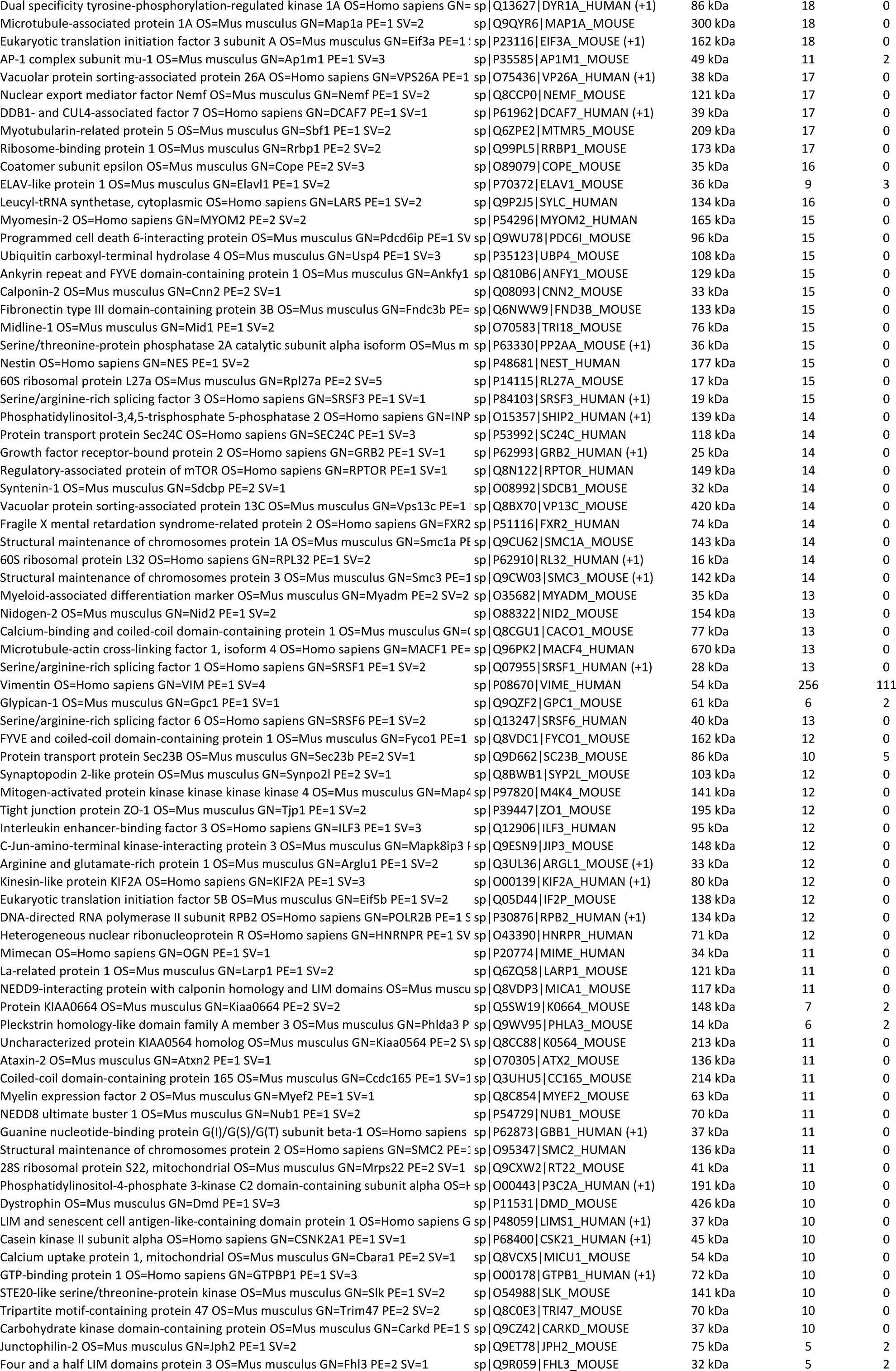
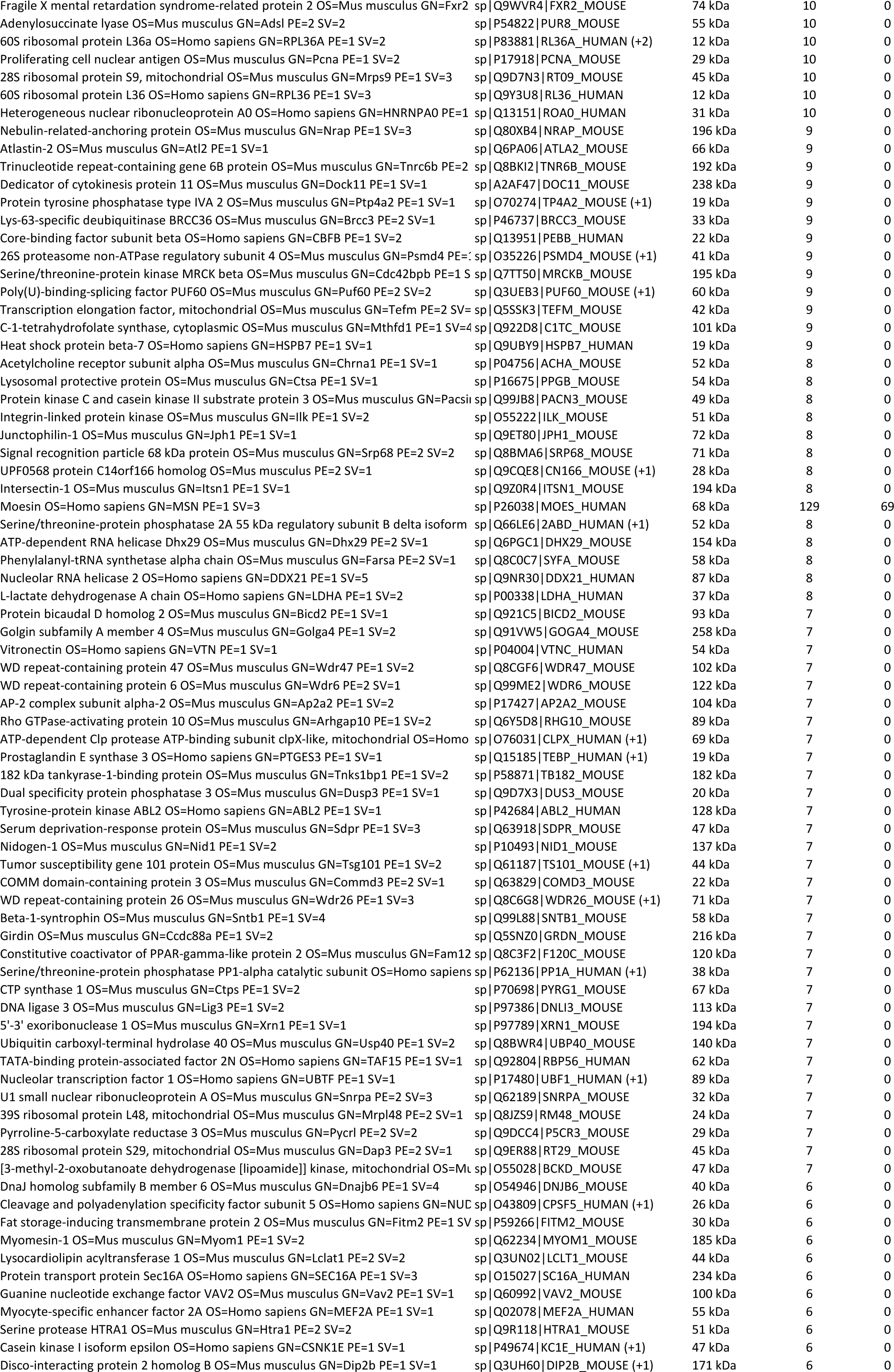
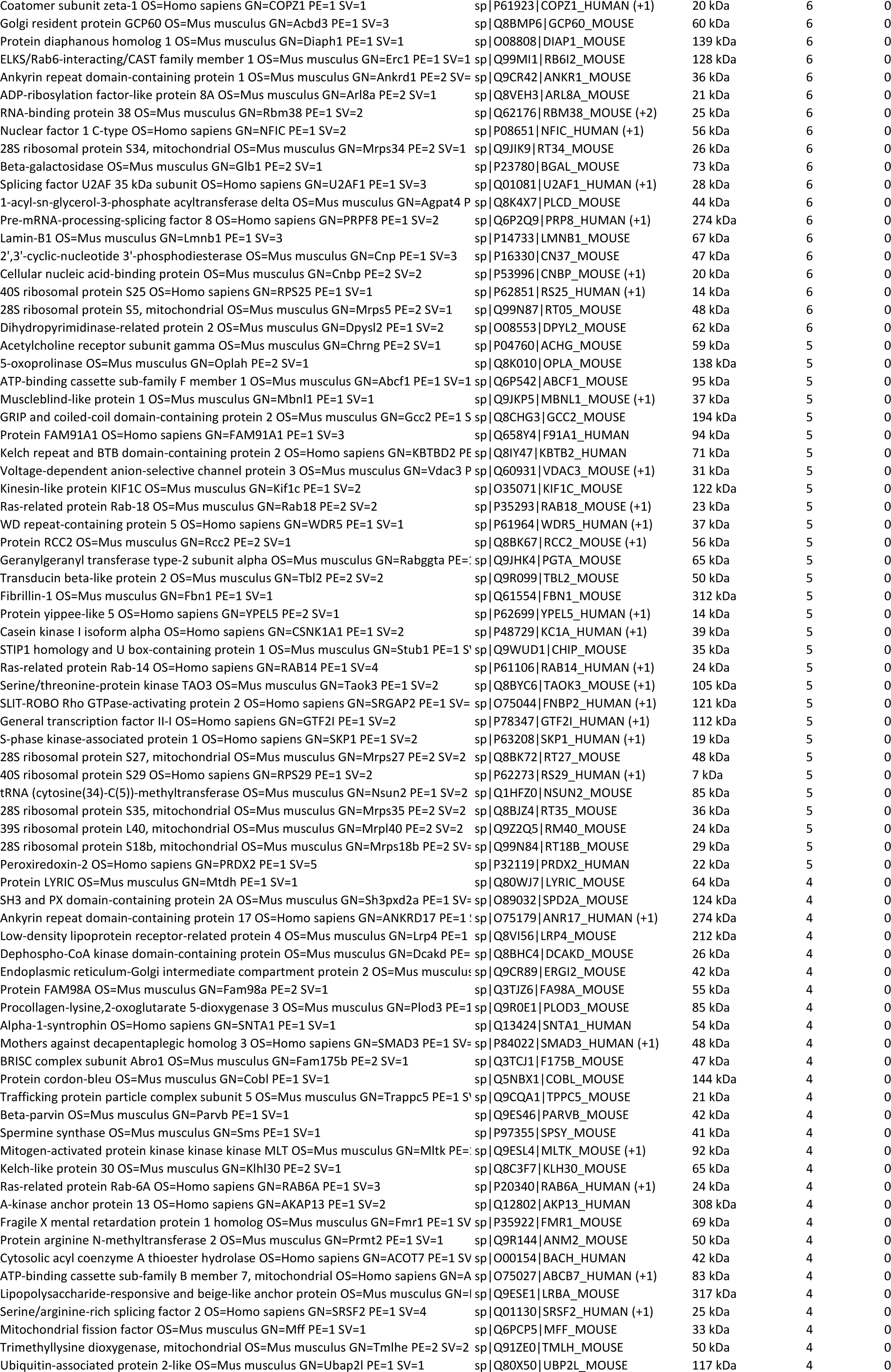
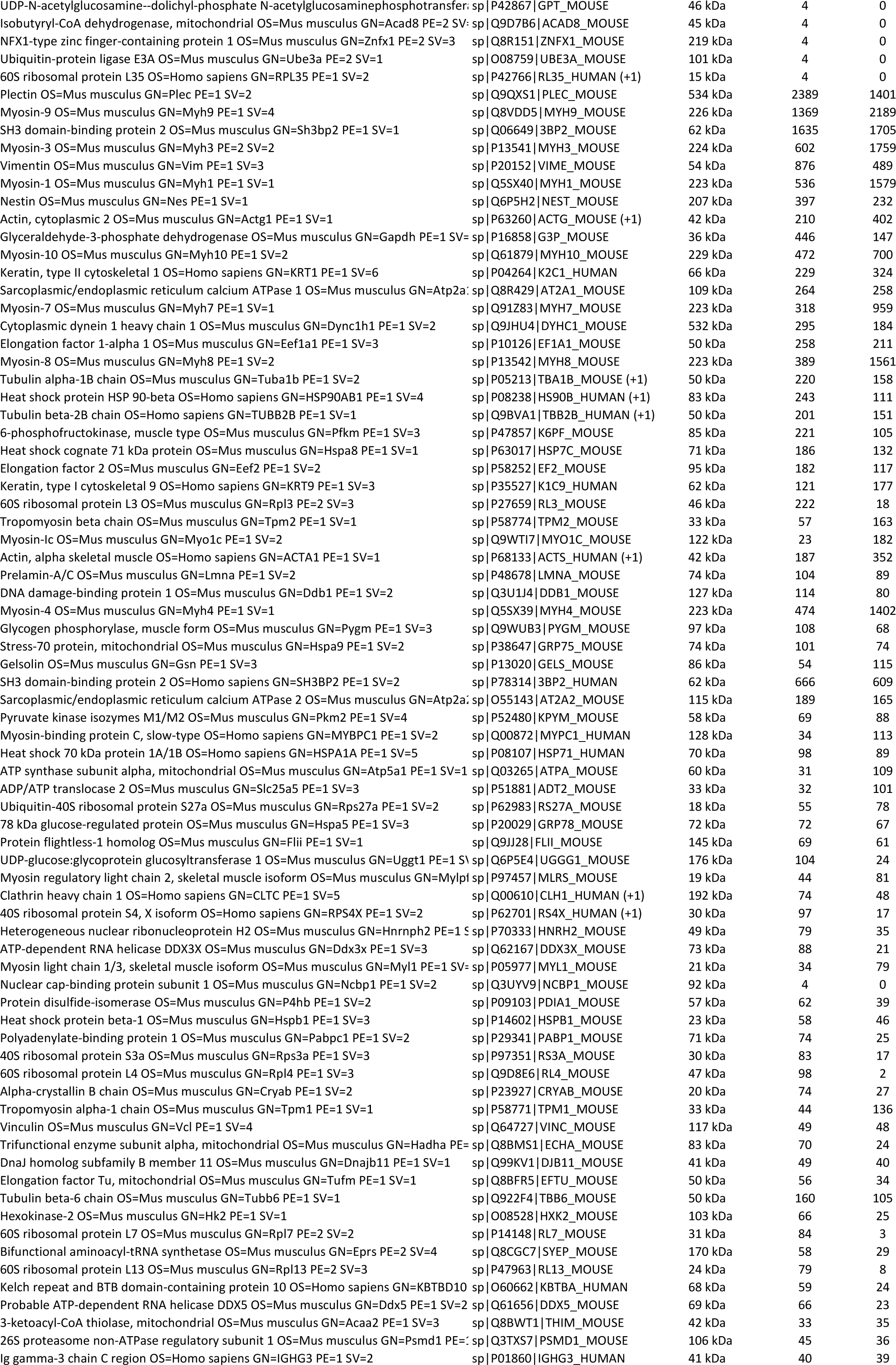
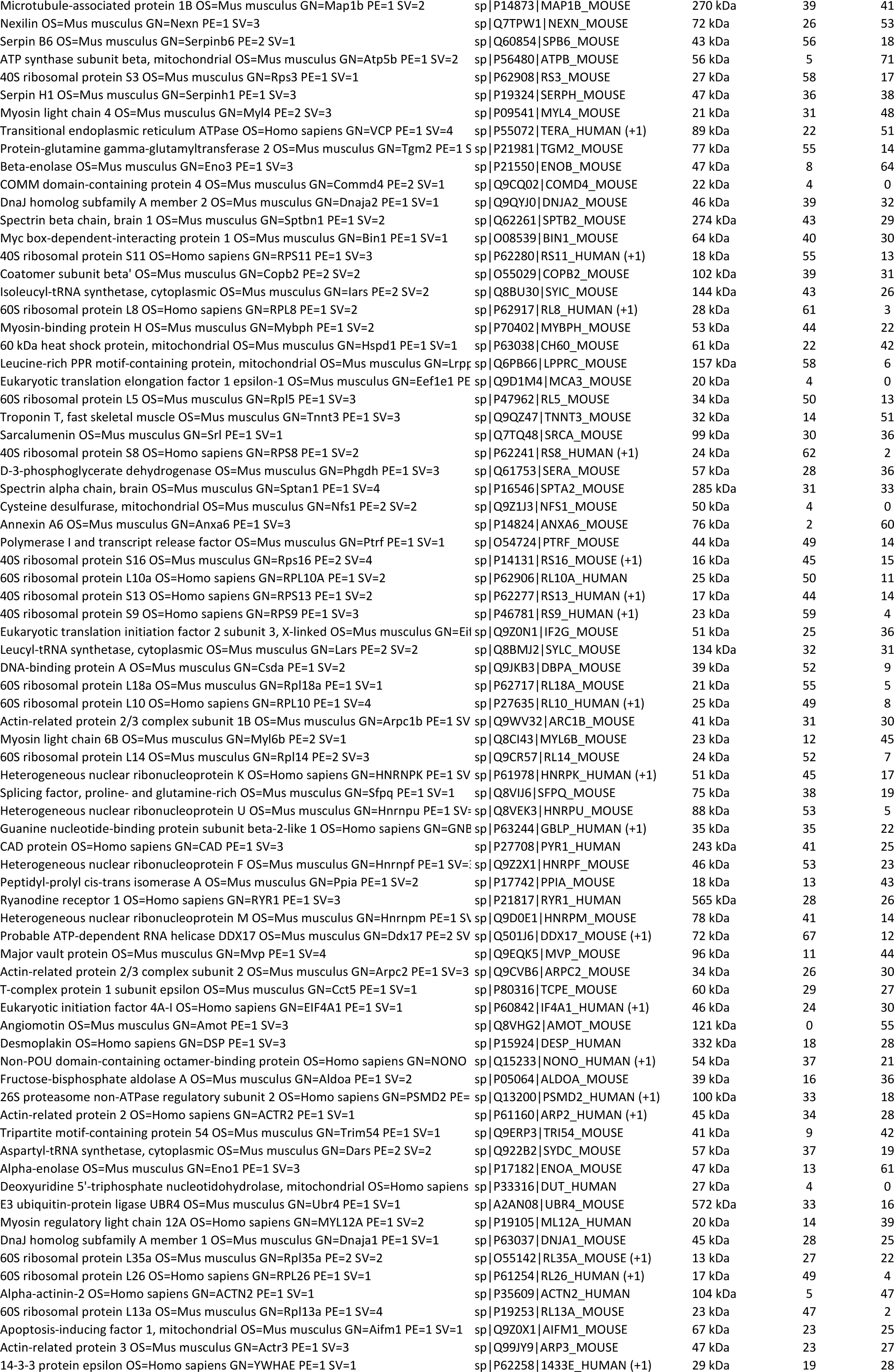
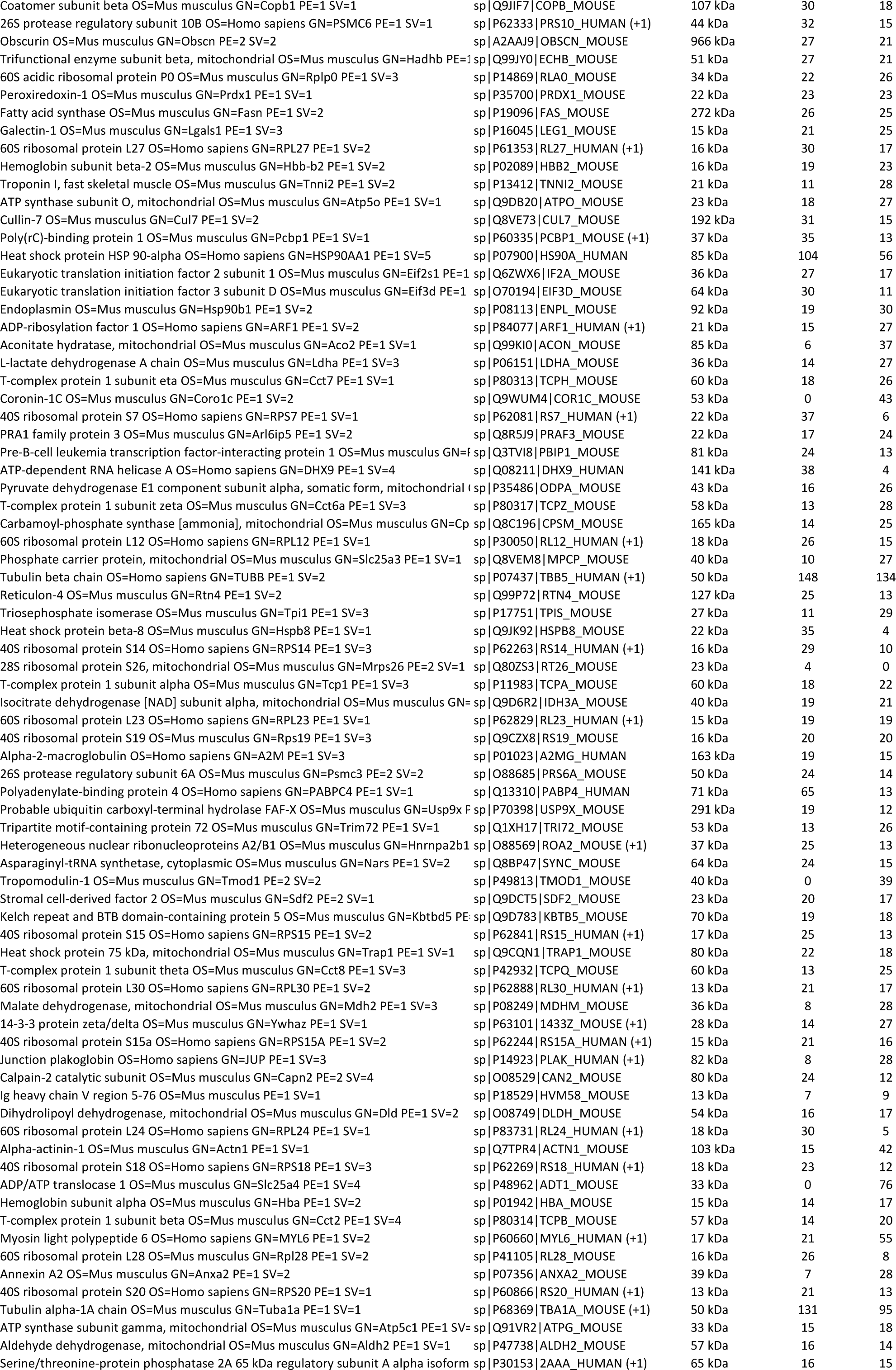
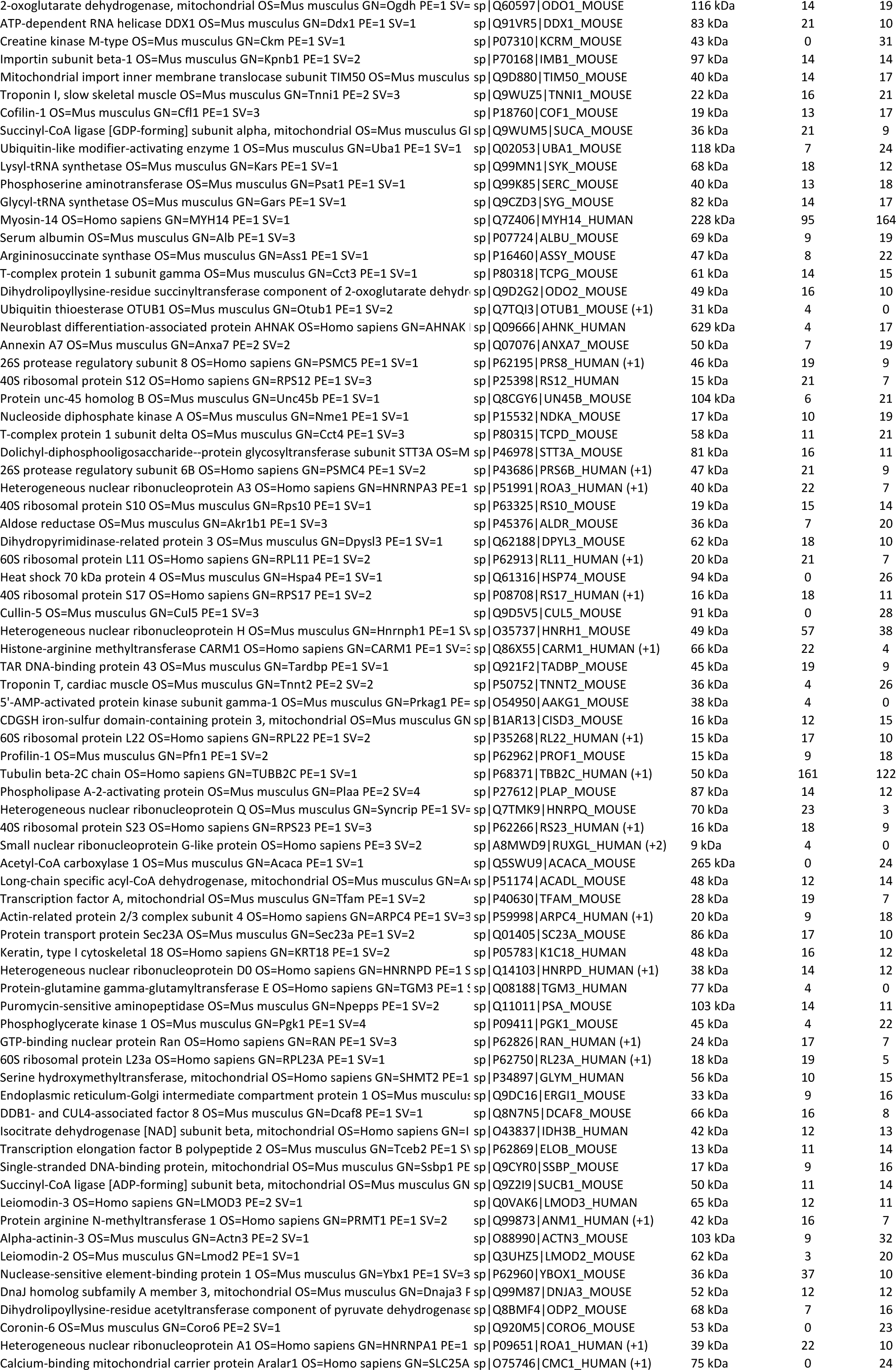
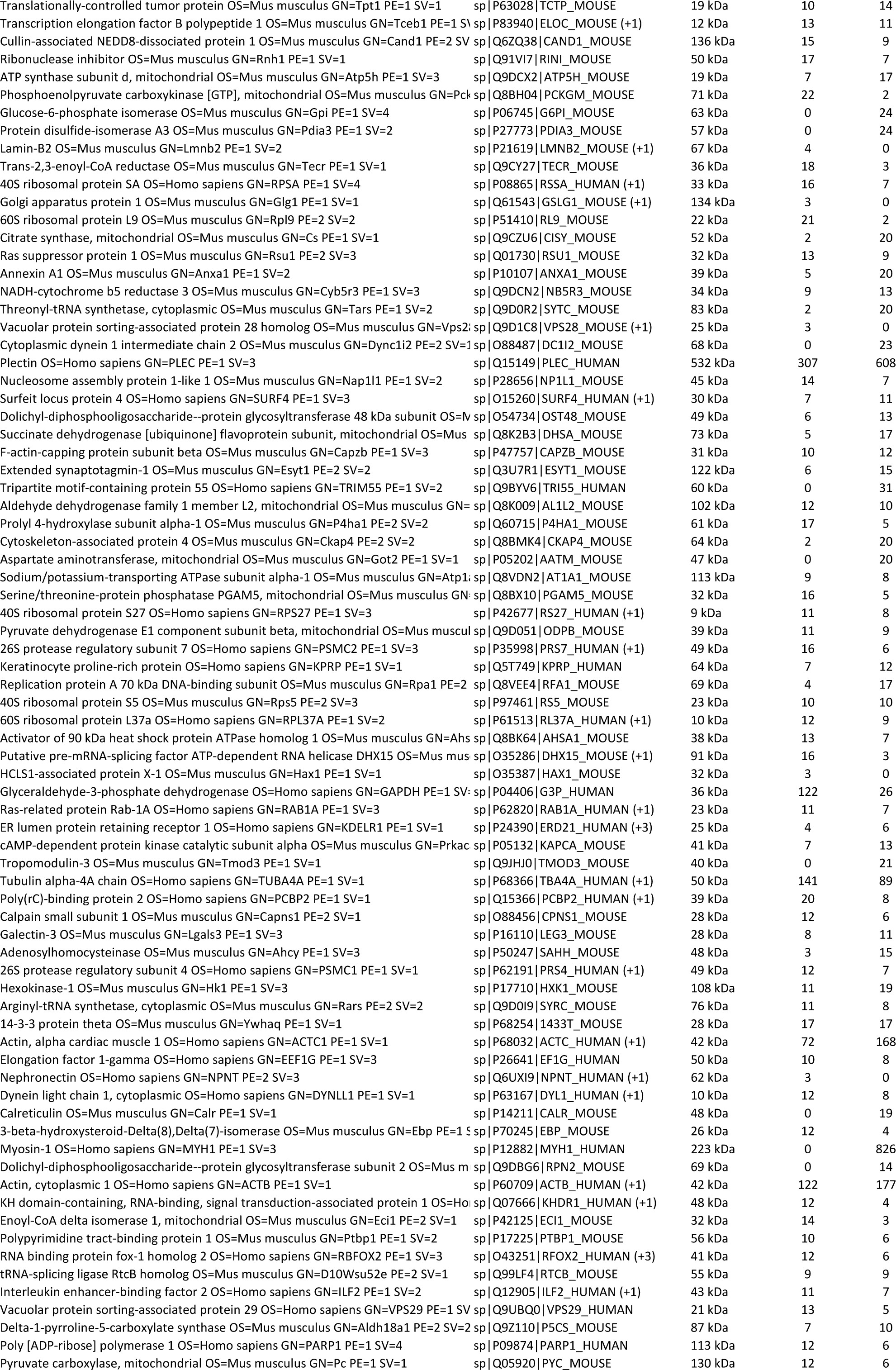
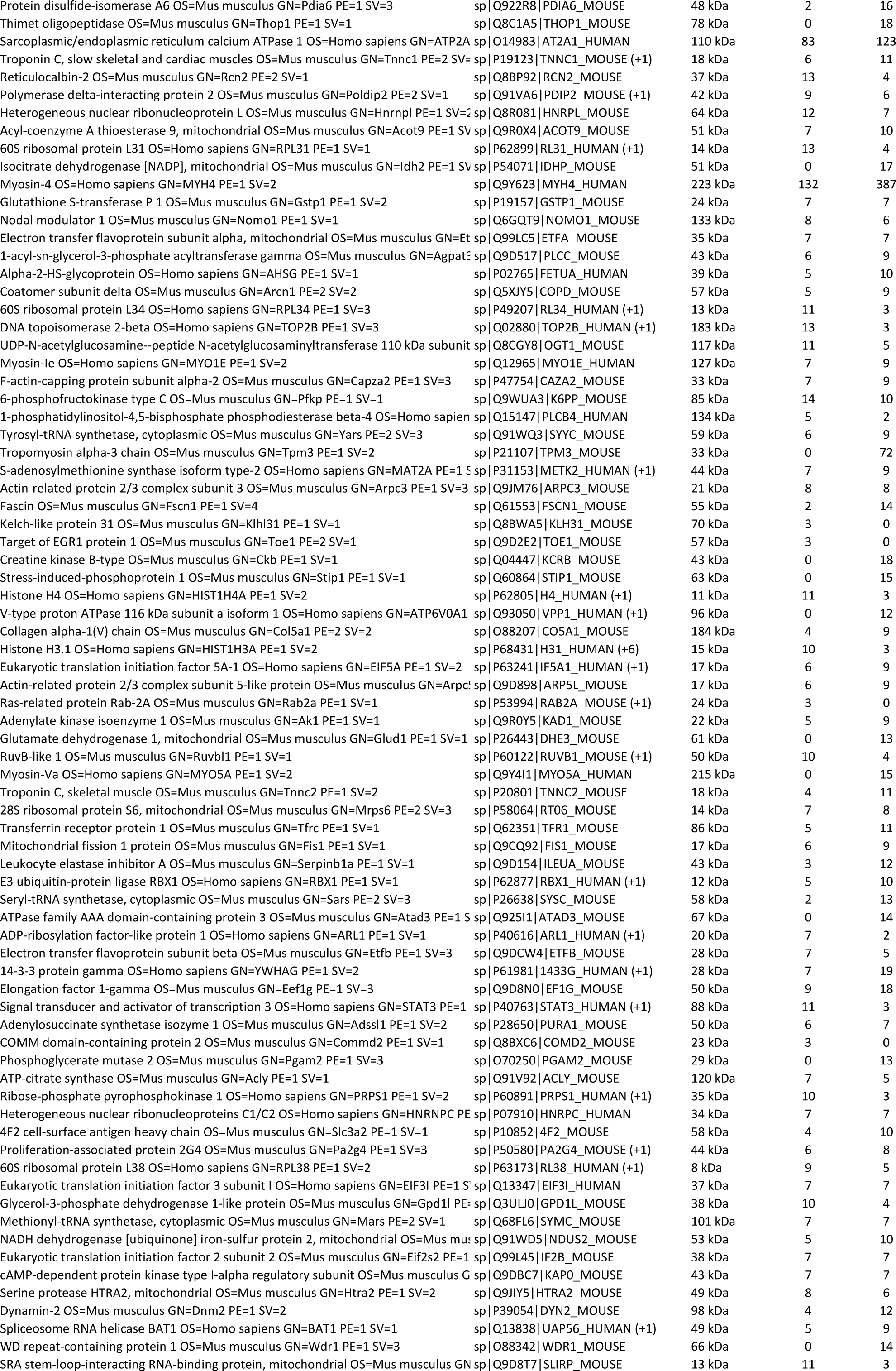
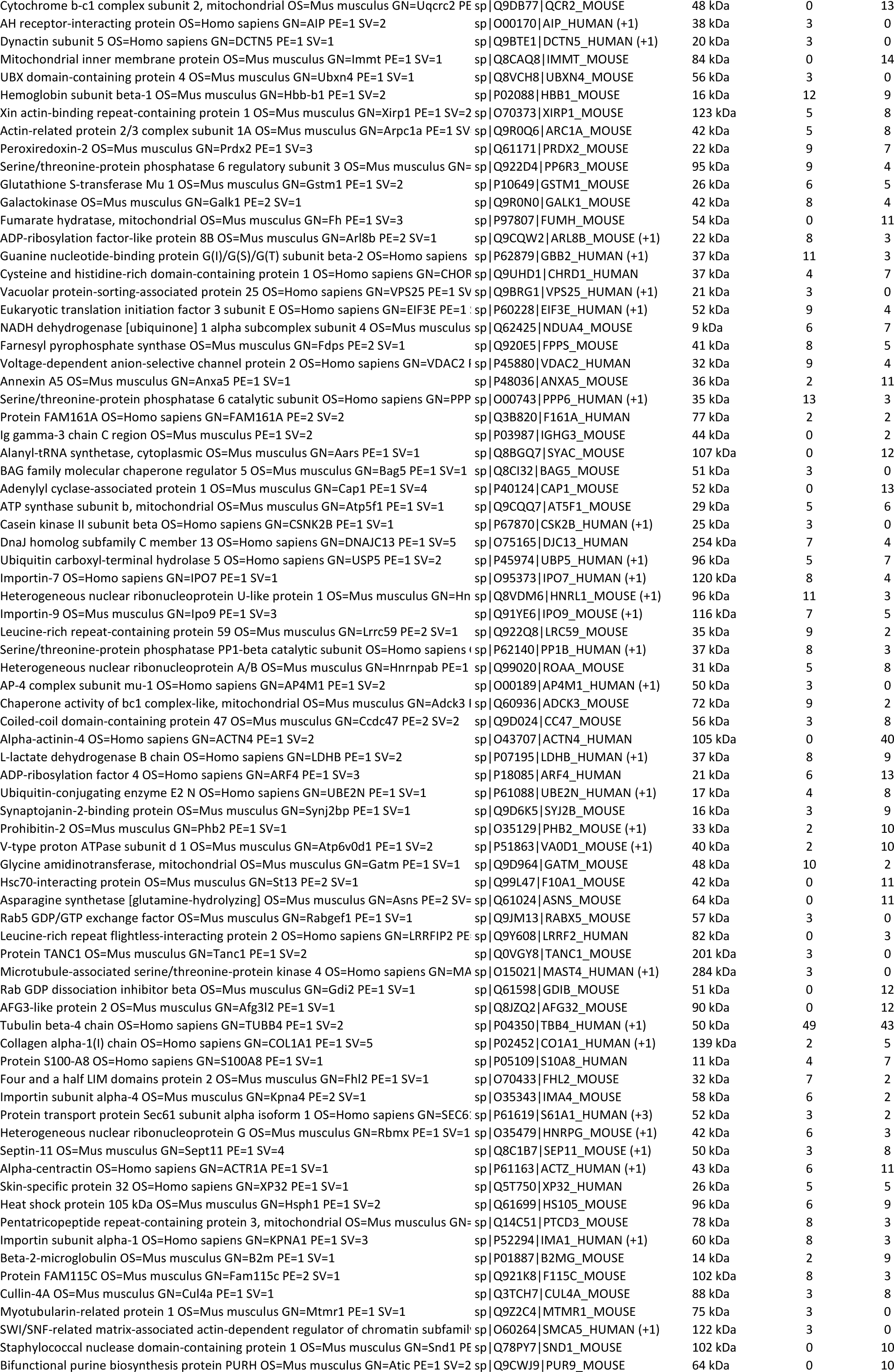
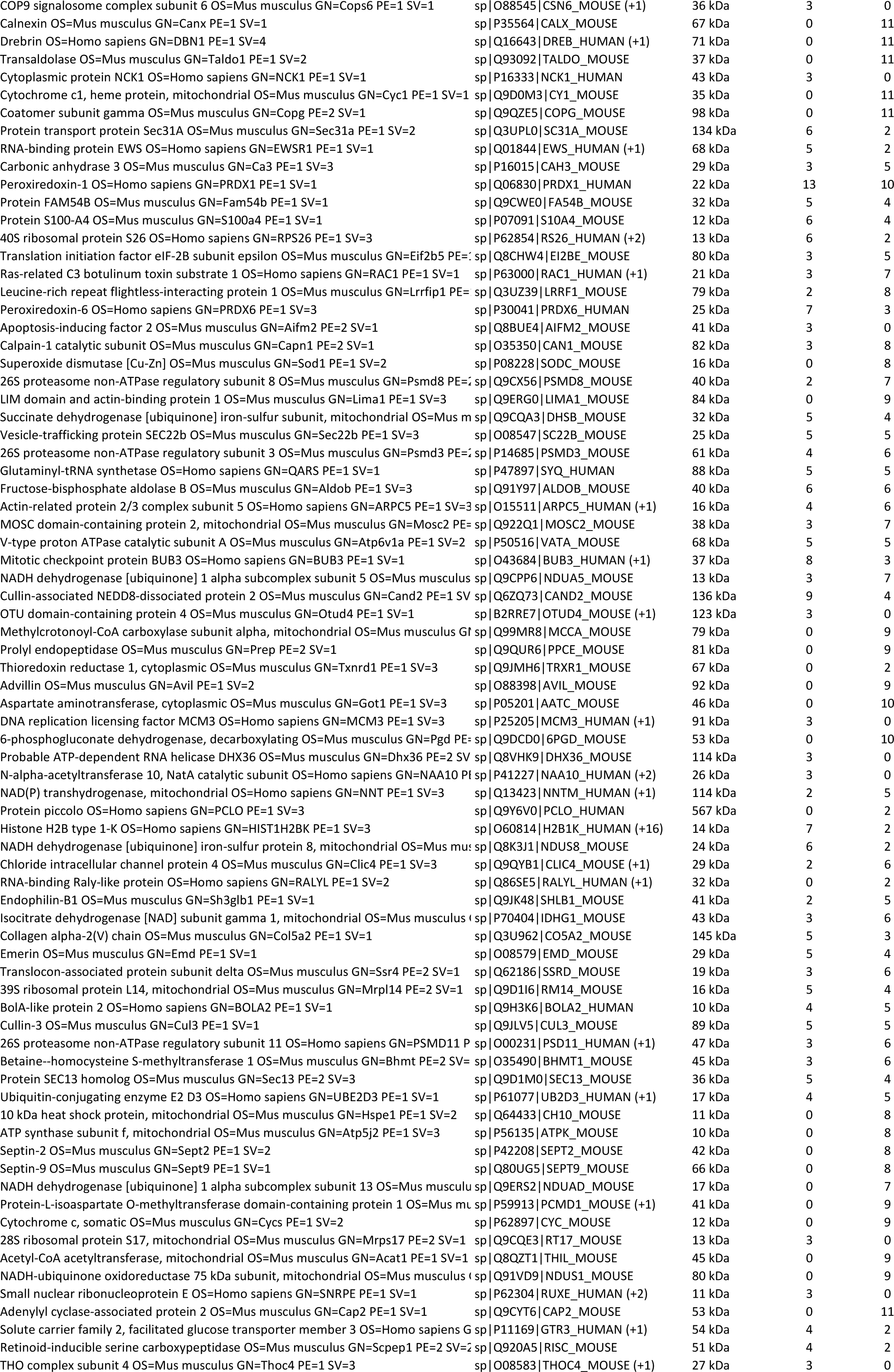
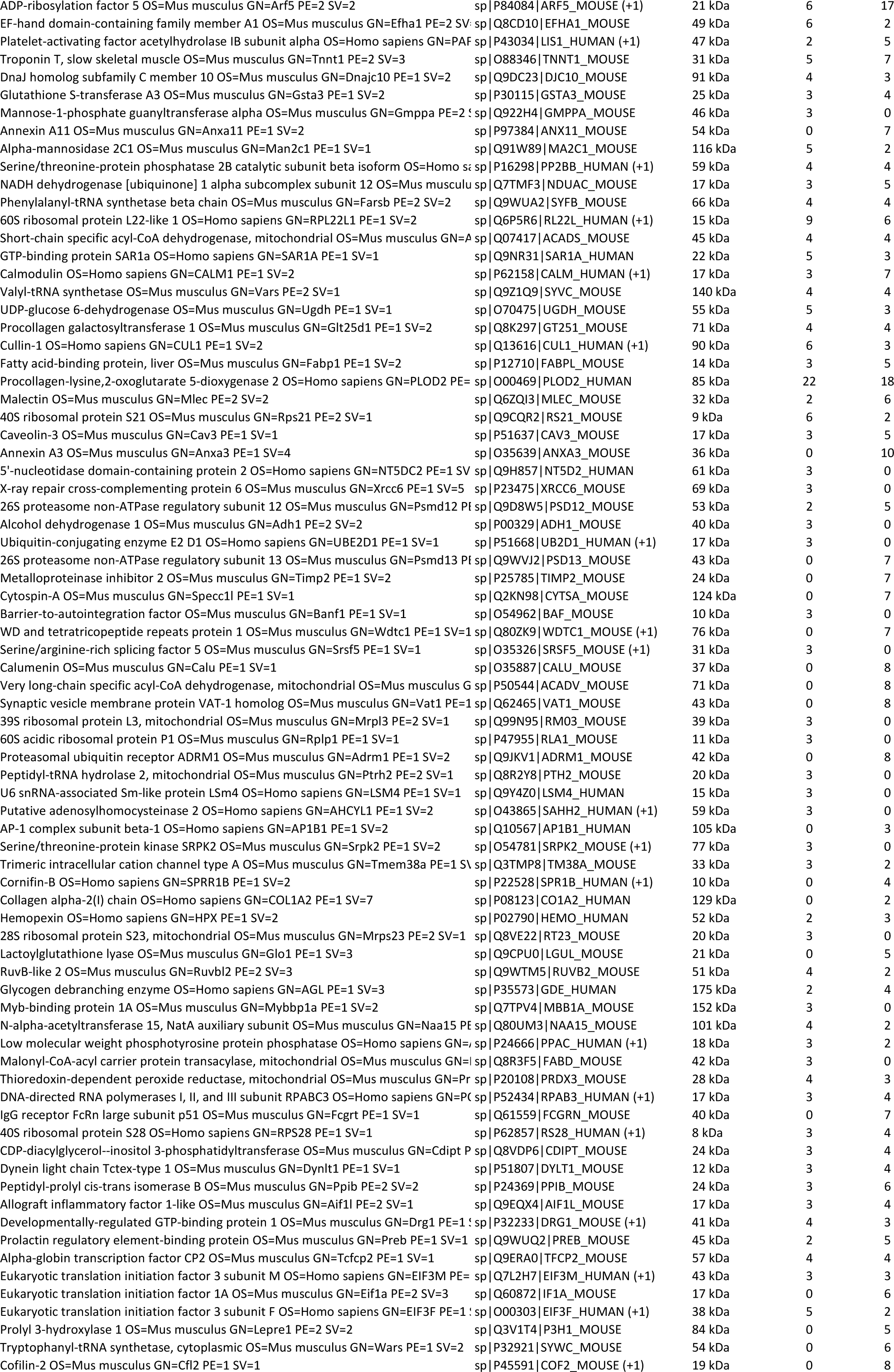
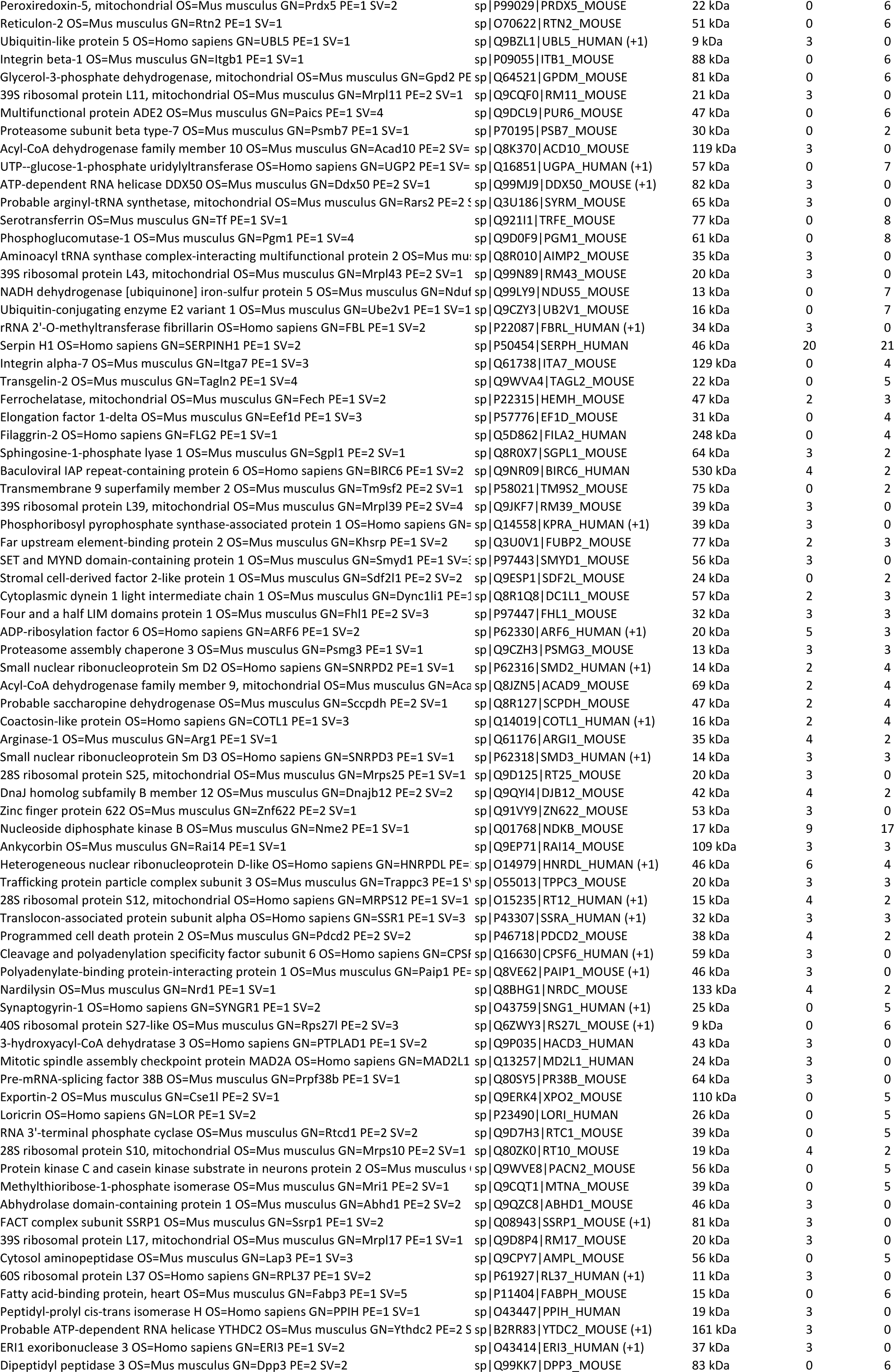
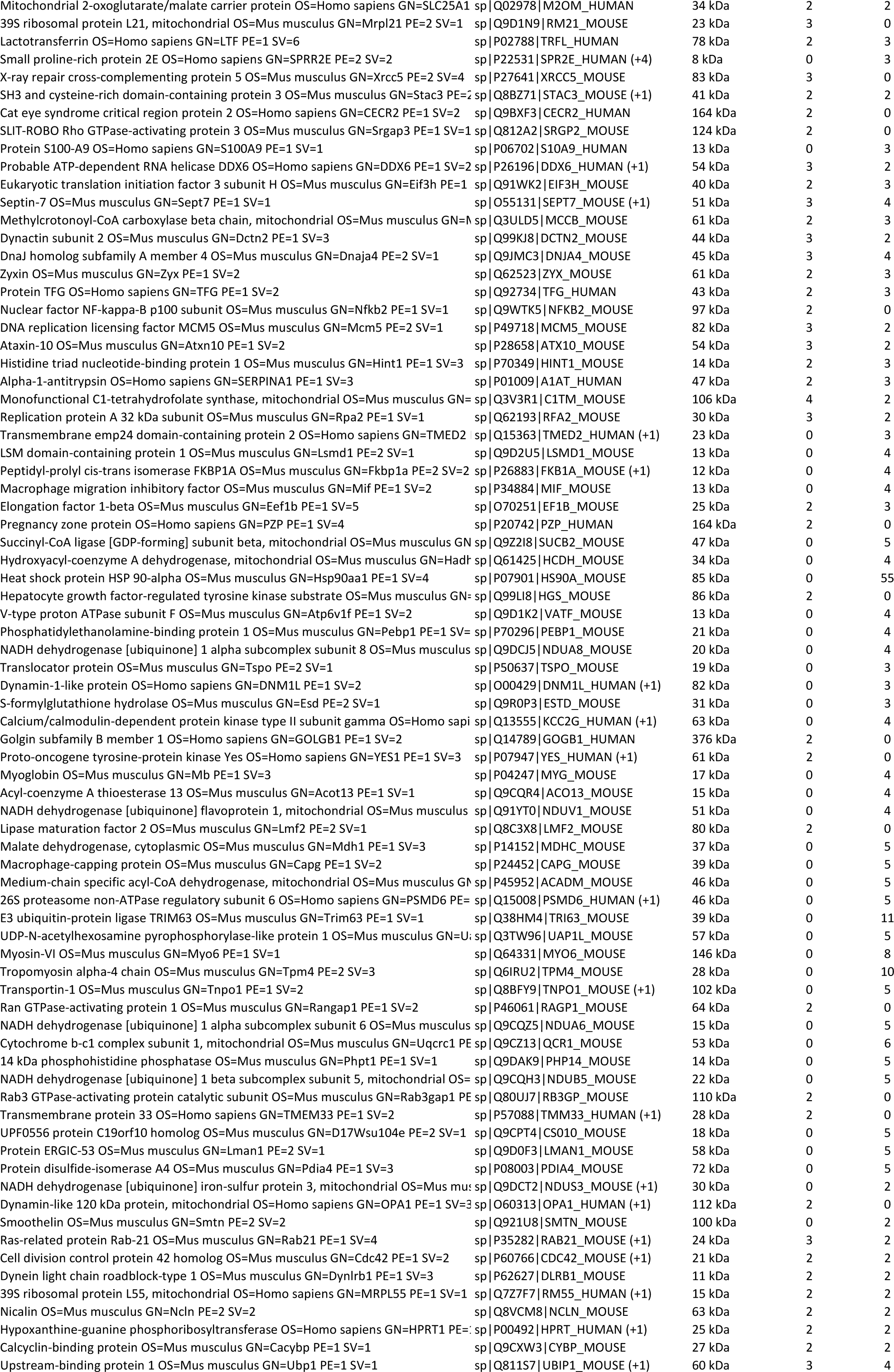
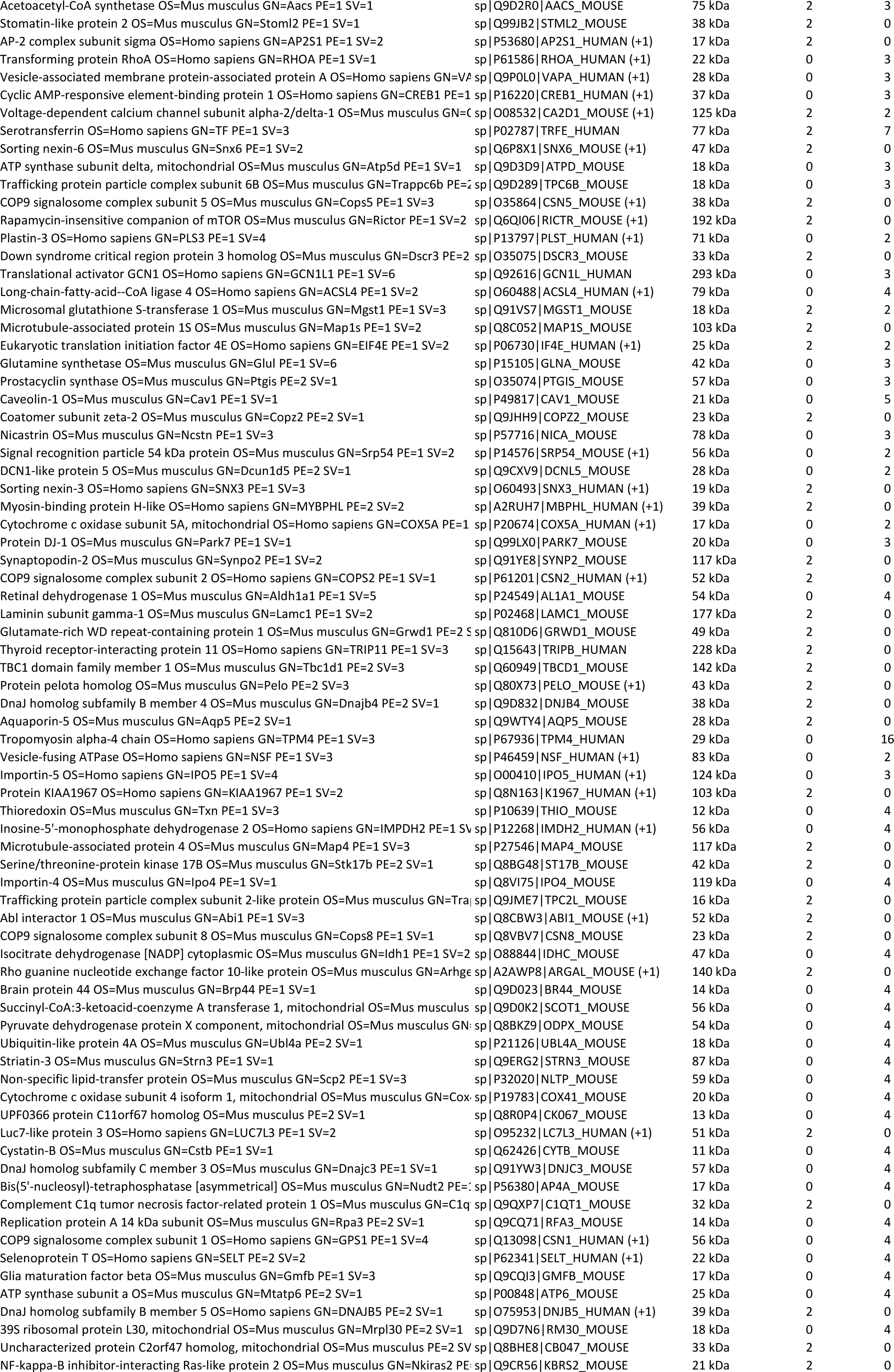
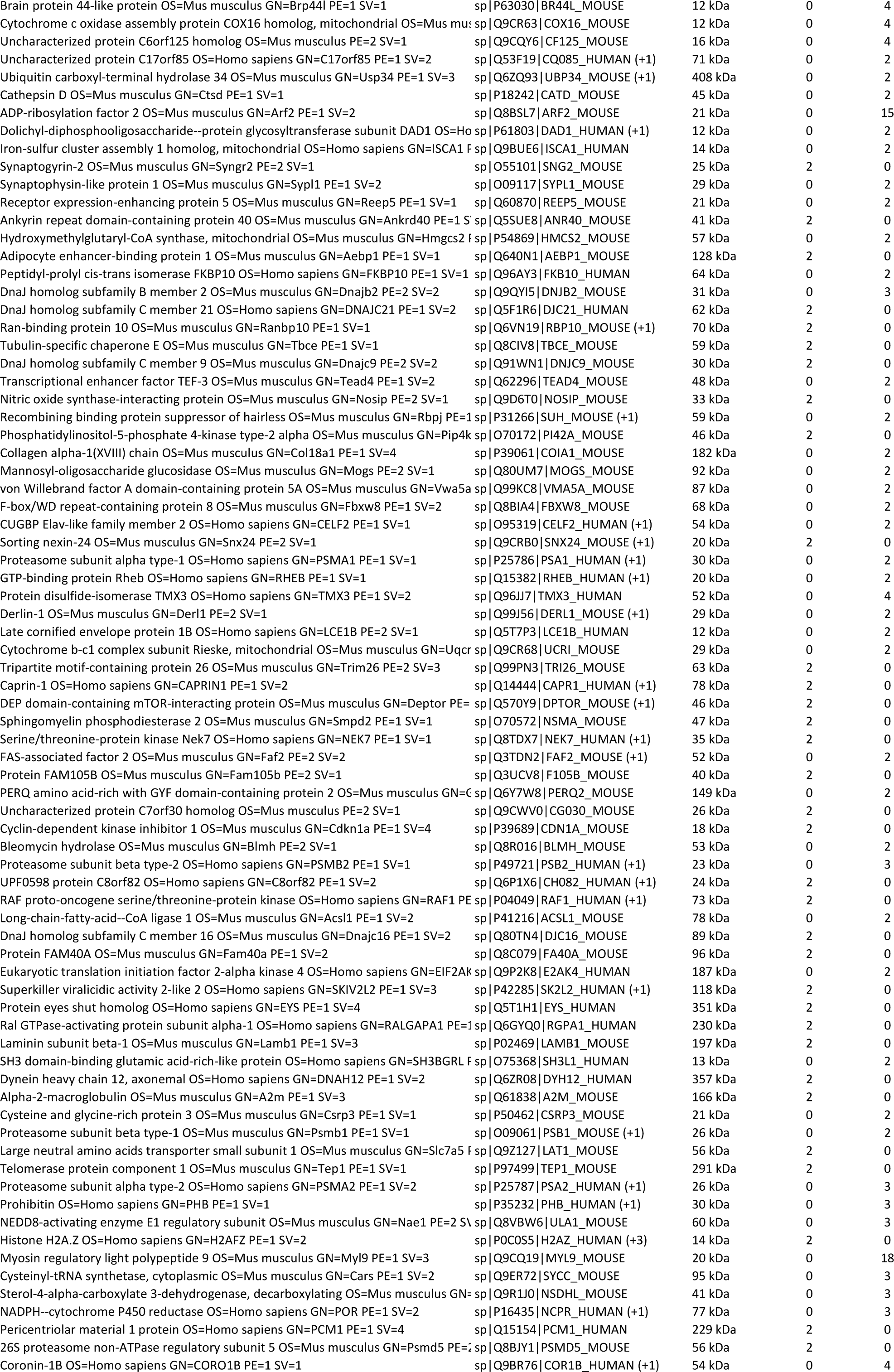
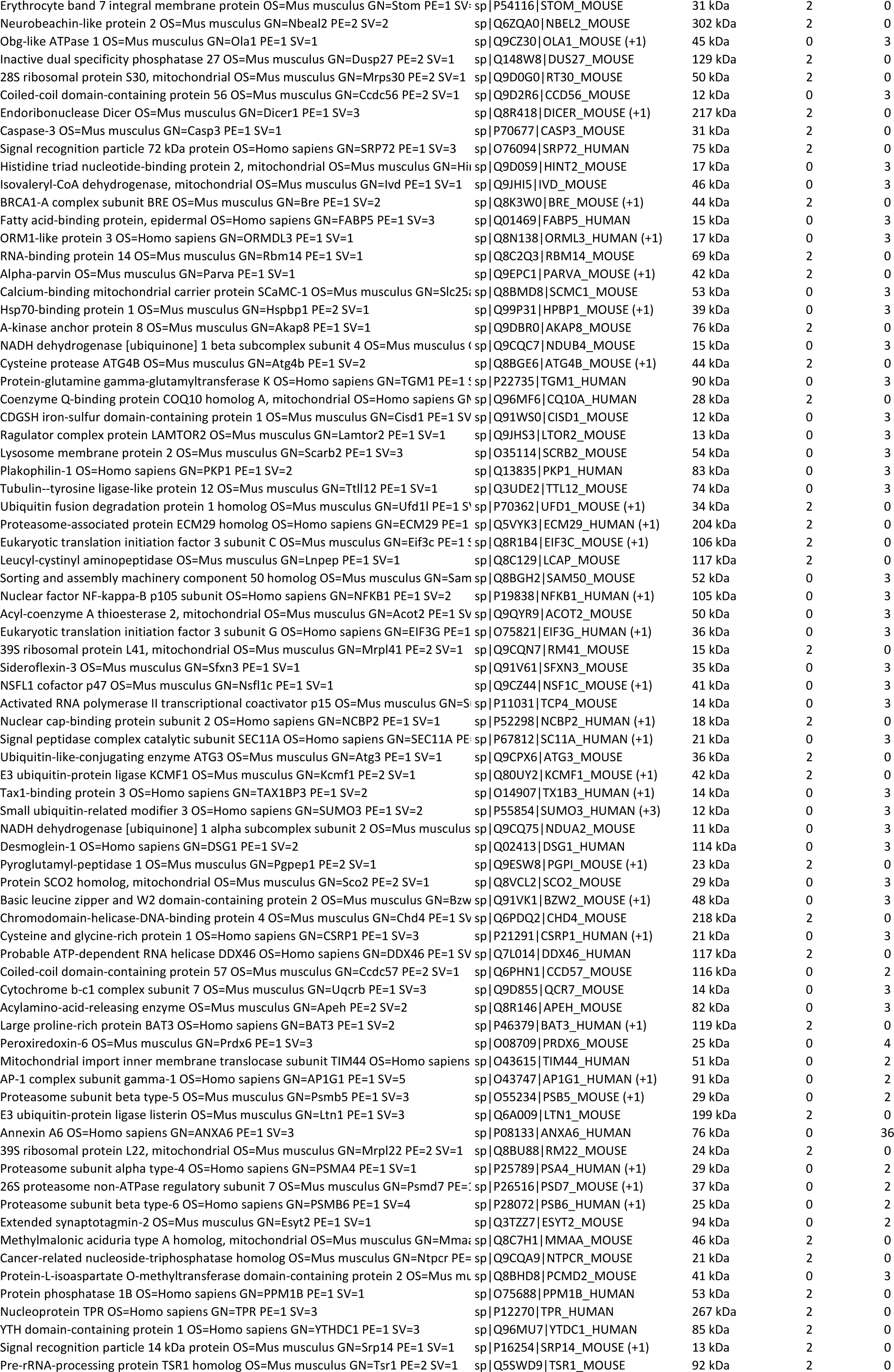
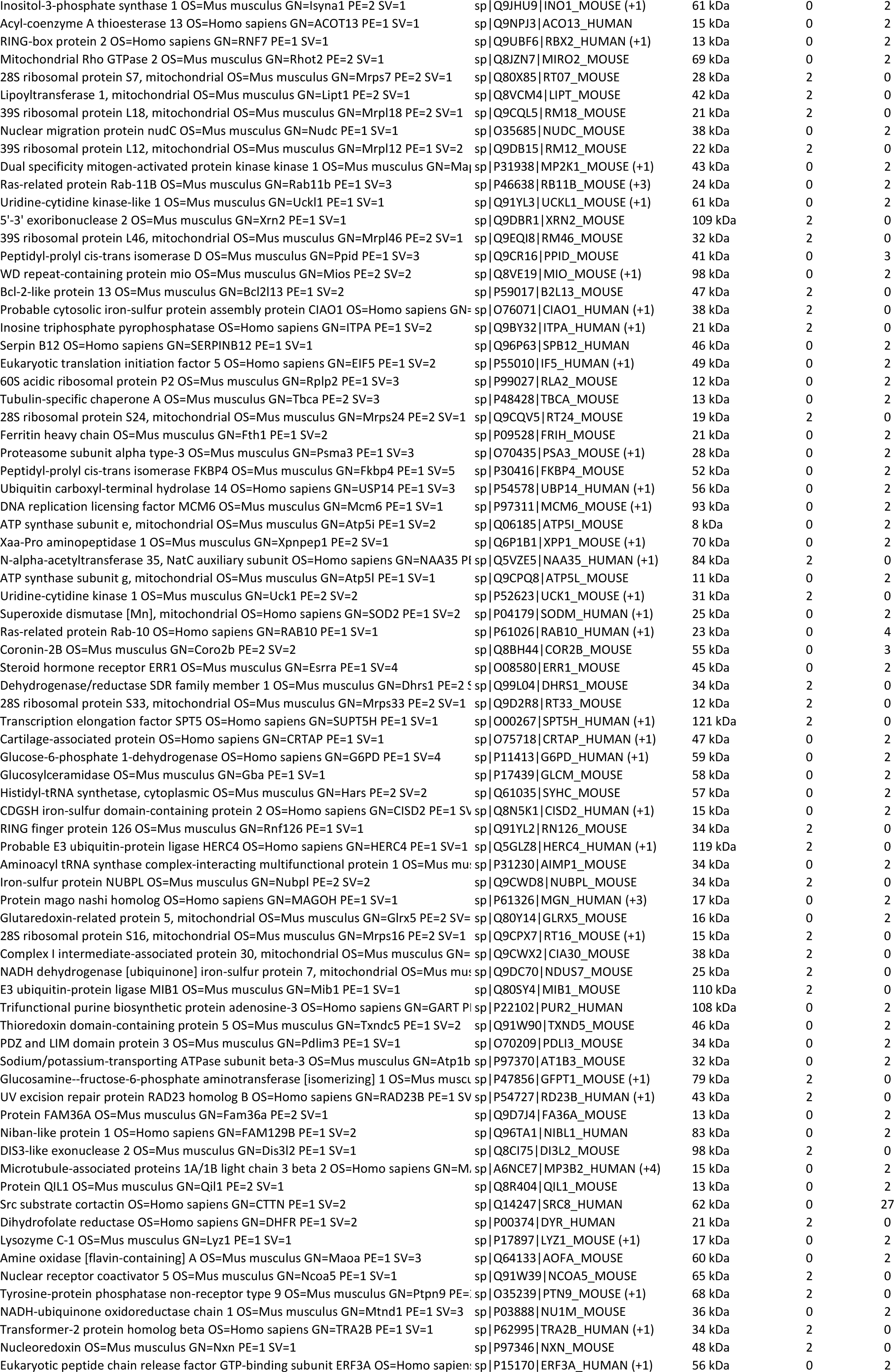
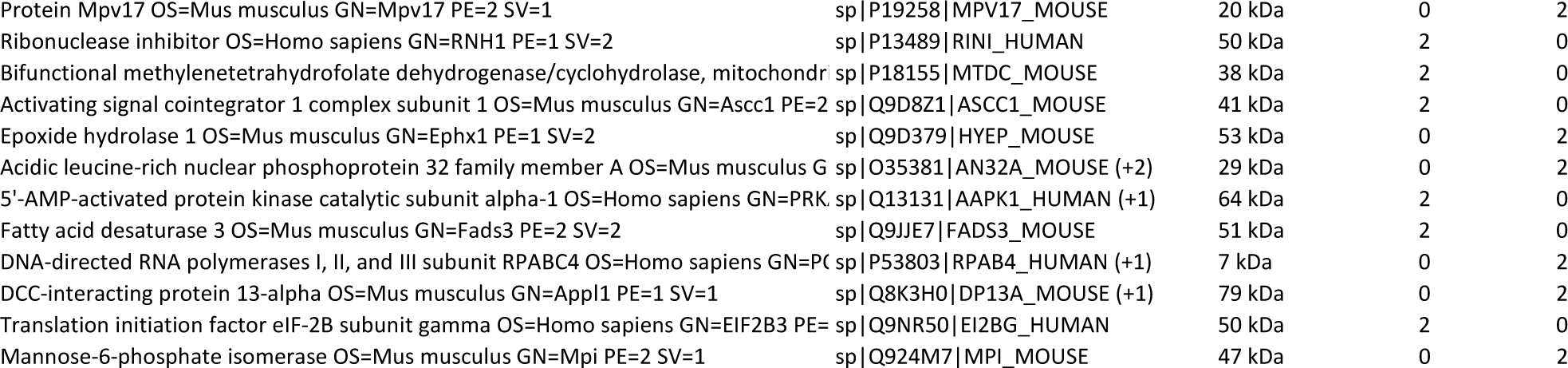
List of proteins interacting with SH3BP2 identified by mass spectrometry analysis.

**Supplementary Table 2.** List of primary antibodies used in the study.

| Antibodies | Source | Identified | Dilution for IF | Dilution for WB | Link |
| --- | --- | --- | --- | --- | --- |
| anti-SH3BP2 | Abnova | H00006452-M01 | 1:200 | 1:3000 | https://www.abnova.com/en-global/product/detail/H00006452-M01 |
| anti-Synaptophysin 1 | Sysy | 101004 | 1:50 | N/A | https://www.sysy.com/product/101004 |
| anti-α-Actinin | Santa Cruz Biotechnology | sc-17829 | 1:100 | N/A | https://www.scbt.com/p/alpha-actinin-antibody-h-2 |
| anti-Ryr | Abcam | ab2868 | 1:100 | N/A | https://www.abcam.com/products/primary-antibodies/ryanodine-receptor-antibody-34c-ab2868.html |
| anti-Titin | Abcam | ab193218 | 1:100 | N/A | https://www.abcam.com/en-pl/products/primary-antibodies/titin-antibody-ab193218 |
| anti-FLAG | Sigma | F3165 | N/A | 1:10000 | https://www.sigmaaldrich.com/PL/pl/product/sigma/f3165?gclid=EAIaIQobChMIwf_i4sbXg_gMVe4RoCR00AwkXEAAYASAAEgIrvfD_BwE |
| anti-GFP | Invitrogen | A11120 | N/A | 1:3000 | https://www.thermofisher.com/antibody/product/GFP-Antibody-clone-3E6-Monoclonal/A-11120 |
| anti-GFP | GeneTex | GTX113617 | N/A | 1:3000 | https://www.genetex.com/Product/Detail/GFP-antibody/GTX113617 |
| anti- α-Tubulin | Abcam | ab18251 | N/A | 1:20000 | https://www.abcam.com/en-pl/products/primary-antibodies/alpha-tubulin-antibody-microtubule-marker-ab18251 |
| anti- c-MYC | Thermofisher scientific | MA1-980 clone 9E10 | N/A | 1:3000 | https://www.thermofisher.com/antibody/product/c-Myc-Antibody-clone-9E10-Monoclonal/MA1-980 |
| anti-GST | Sigma | G7781 | N/A | 1:2000 | https://www.sigmaaldrich.com/PL/pl/product/sigma/g7781 |
| anti-RFP | Rockland | 600-401-379 | N/A | 1:1000 | https://www.rockland.com/categories/primary-antibodies/rfp-antibody-pre-adsorbed-600-401-379/ |
| anti-GAPDH | Sigma | MAB374 | N/A | 1:10000 | https://www.sigmaaldrich.com/PL/pl/product/mm/mab374 |
| anti-AChRδ | Thermofisher scientific | MA3-043 | N/A | 1:3000 | https://www.fishersci.com/shop/products/chrna1-monoclonal-antibody-88b-invitrogen/MA3043 |
| antiAChRα1 | Santa Cruz Biotechnology | sc-65829 | N/A | 1:1000 | https://datasheets.scbt.com/sc-65829.pdf |
| anti Neurofilament | Millipore | AB1991 | 1:300 | N/A | https://www.merckmillipore.com/PL/pl/product/Anti-Neurofilament-H-200-kDa-Antibody-lysine-serine-proline-repeat,MM_NF-AB1991 |

**Supplementary Fig. 1.**
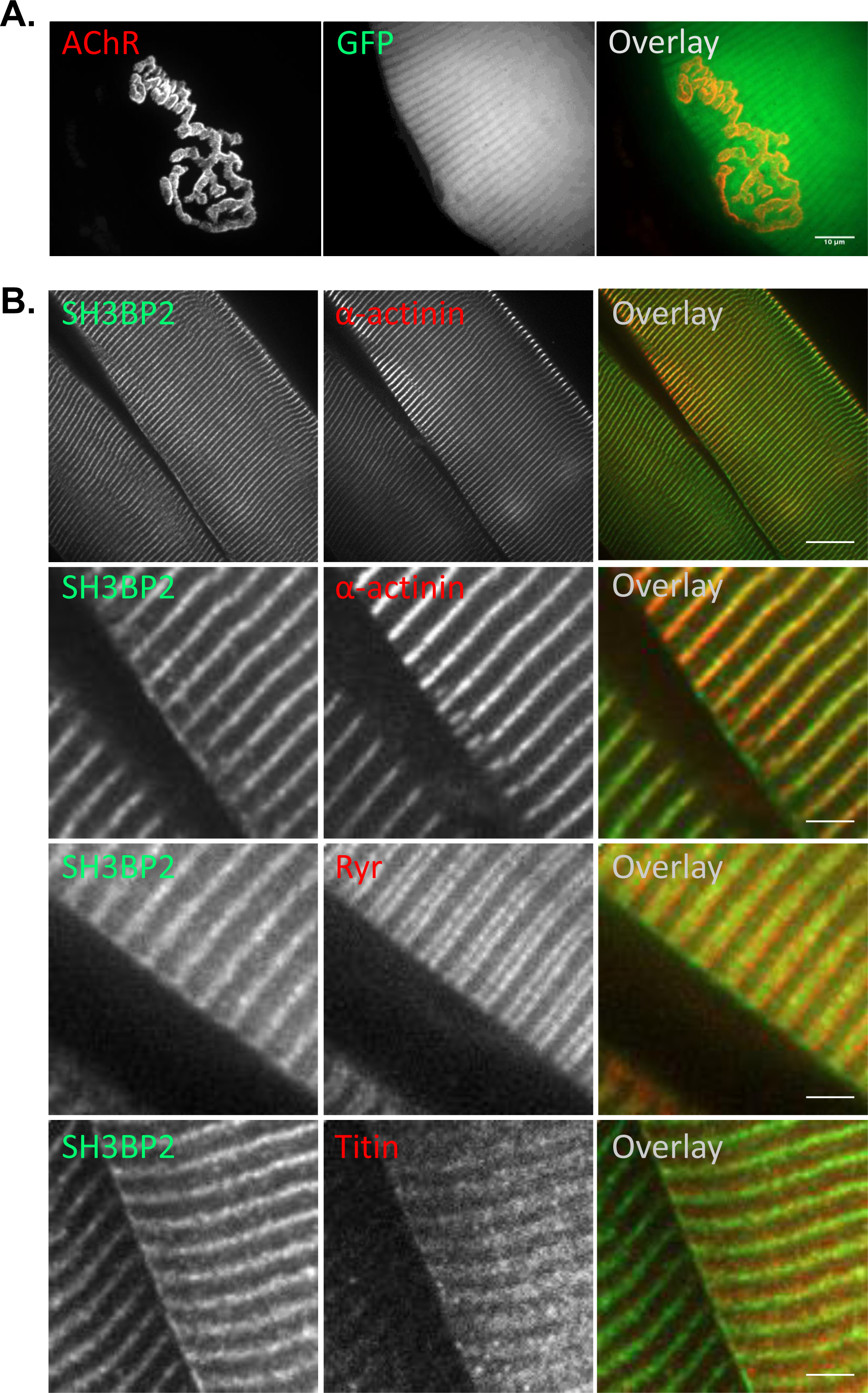
Localization of SH3BP2 with the contractile machinery. (A) A representative image showing the Tibialis anterior electroporated with a plasmid expressing GFP - a negative control for SH3BP2 localization experiments. (B) Representative images showing the colocalization of SH3BP2 immunofluorescence with α-actinin, Ryanodine receptors, and Titin immunoreactivity in the Tibialis anterior muscle. Scale bars 10 μm.

**Supplementary Fig. 2.**
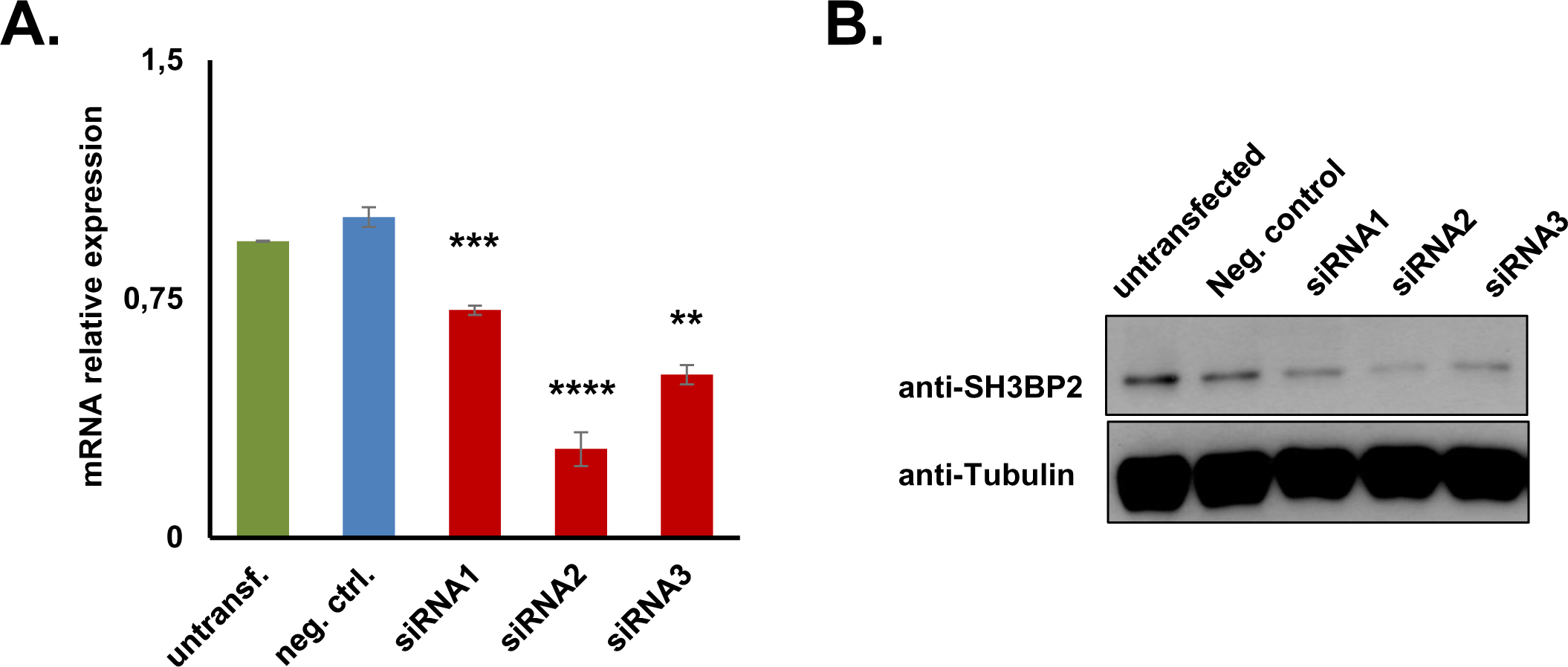
Validation of SH3BP2 KD in C2C12 myotubes. (A) Validation of SH3BP2 knock-down at the mRNA level using qRT-PCR and at the protein level using Western blot (B). C2C12 myotubes were transfected with either siRNAs targeting SH3BP2 or a scrambled, non-targeting siRNA (neg. ctrl.).

**Supplementary Fig. 3.**
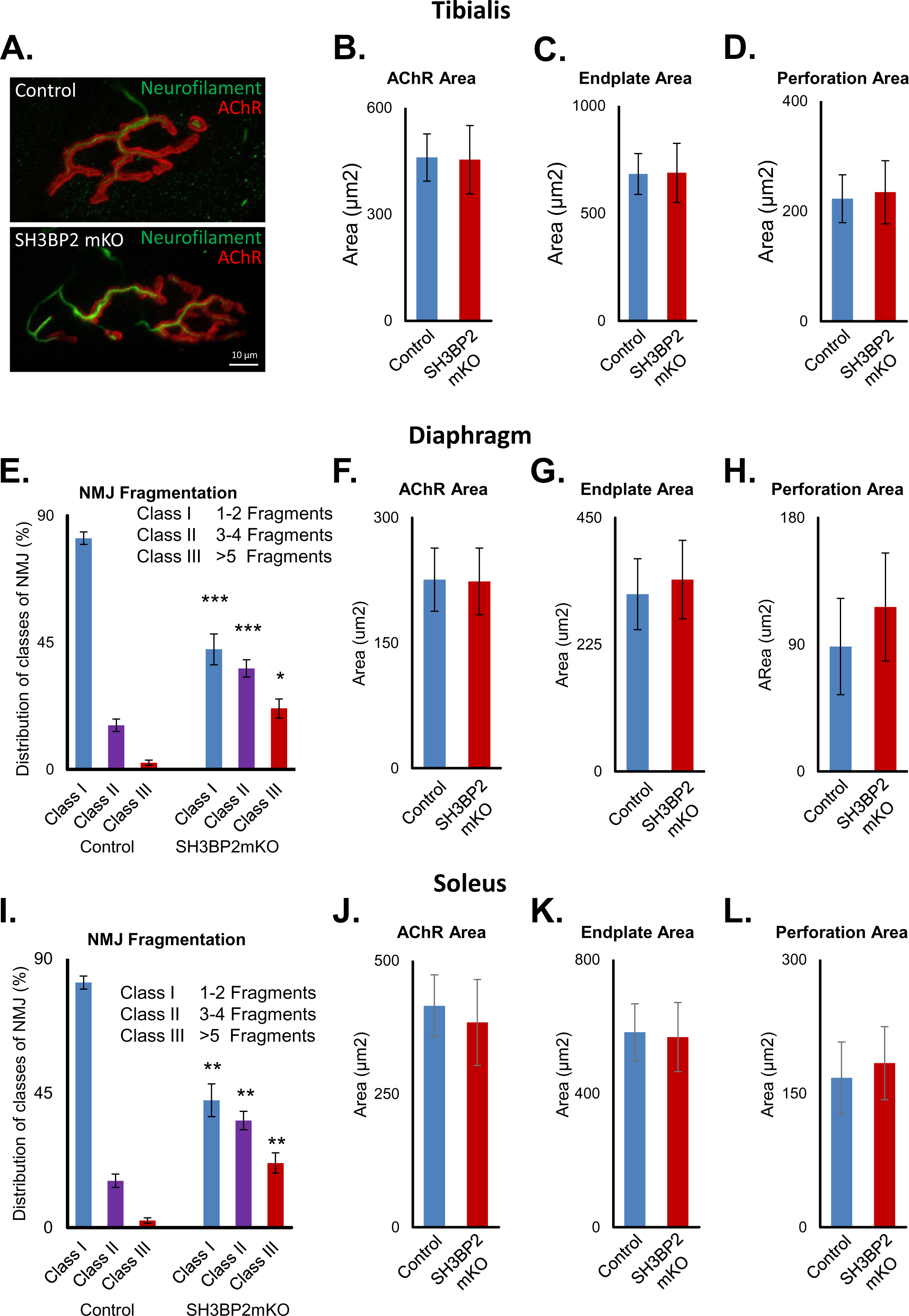
Increased NMJ fragmentation in the diaphragm and soleus muscles of SH3BP2 mKO mice. (A-D) Tibialis muscle analysis. (A) NMJs in SH3BP2 mKO mice are more fragmented, but the nerve morphology appears normal as evaluated based on the neurofilament immunoreactivity. SH3BP2 mKO mice have no significant alterations in the AChR area (B), endplate area (C), or NMJ perforation area (D) in the Tibialis anterior muscles. Results are mean ± SEM; n = 5 mice per group; more than 132 NMJ per group; p > 0,05; two-tailed Student’s t-test. (E-H) Diaphragm muscle analysis. (E) Increase fragmentation of NMJs in the diaphragm of SH3BP2 mKO mice. Results are mean ± SEM; n = 4 mice per group; more than 105 NMJ per group; ***p < 0.001, *p < 0.05; one-way ANOVA. SH3BP2 mKO mice have no significant alterations in the AChR area (F), endplate area (G), or NMJ perforation area (H) in diaphragm muscles. Results are mean ± SEM; n = 4 mice per group; more than 84 NMJ per group; p > 0,05; two-tailed Student’s t-test. (I-L) Soleus muscle analysis. (I) Increase fragmentation of NMJs in soleus muscles of SH3BP2 mKO mice. Results are mean ± SEM; n=4 mice per group; more than 94 NMJ per group; ***p < 0.001, *p < 0.05; one-way ANOVA. SH3BP2 mKO mice show no significant alterations in the AChR area (J), endplate area (K), or NMJ perforation area (L) in soleus muscles. Results are mean ± SEM; n = 4 mice per group; more than 95 NMJ per group; p > 0,05; two-tailed Student’s t-test.

**Supplementary Fig. 4.**
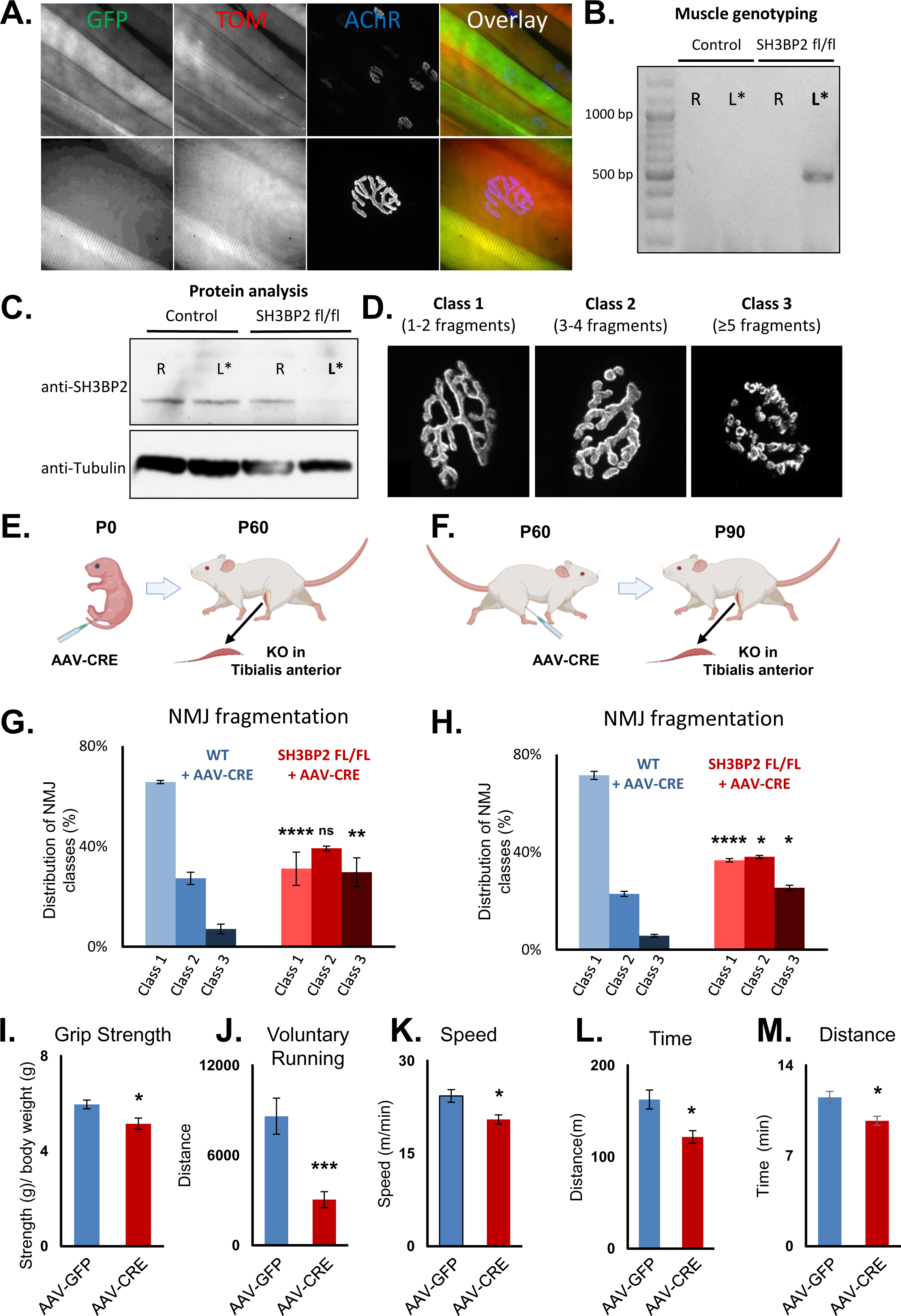
AAV-mediated deletion of SH3BP2 in Tibialis anterior leads to NMJ fragmentation and reduced muscle strength. (A) Representative images showing the tdTomato (TOM) expression in STOP-TOM Cre reporter mice after AAV-Cre-GFP (AAV-Cre) infection, a proof of principle experiment. The bottom panel shows NMJ at higher magnification. (B) Muscle genotyping shows an expected PCR band of approx. 500 bp confirming the deletion of SH3BP2 exon 6 and 7 after AAV-Cre infection (SH3BP2 fl/fl, left side, indicated with a star). The controls include the contralateral leg of the same animal without infection and infected and uninfected Tibialis from WT animals. (C) Western blot analysis showing reduced SH3BP2 protein expression in SH3BP2 fl/fl mice after AAV-Cre infection (indicated with a star). (D) Classification of NMJs according to their fragmentation (Class I – 1-2 fragments, Class II – 3-5 fragments, Class III – more than 5 fragments). (E and F) The experimental design for results shown in G and H, respectively. Tibialis anterior NMJ fragmentation quantification for animals infected at P0 (G) and P60 (H). The control group constitutes Tibialis muscles from WT animals after AAV-Cre infection. Results are mean ± SEM; n = 3 mice per group, except P0 AAV infection in SH3BP2 fl/fl group, (n = 2); more than 90 NMJs were analyzed; *p < 0.05; **p < 0.01; ****p <0.0001; two-tailed Student’s t-test. (I) Decreased grip strength of SH3BP2 ^fl/fl^ mice injected bilaterally with AAV-Cre virus into the Tibialis anterior muscles at P90. The control mice were bilaterally injected with the AAV-GFP virus. (J) Reduced voluntary running distance by SH3BP2 ^fl/fl^ mice injected with AAV-Cre. (K) Reduction in maximal speed (K), duration of running (L), and distance (M) run in a treadmill experiment. Results are mean ± SEM; n = 10 mice per group; *p < 0.05 and ***p < 0.001; two-tailed Student’s t-test.

## References

1 Fambrough, D. M. Control of acetylcholine receptors in skeletal muscle. Physiol Rev 59, 165–227 (1979). 10.1152/physrev.1979.59.1.165

2 Fertuck, H. C. & Salpeter, M. M. Localization of acetylcholine receptor by 125I-labeled alpha-bungarotoxin binding at mouse motor endplates. Proc Natl Acad Sci U S A 71, 1376–1378 (1974). 10.1073/pnas.71.4.1376

3 Flucher, B. E. & Daniels, M. P. Distribution of Na+ channels and ankyrin in neuromuscular junctions is complementary to that of acetylcholine receptors and the 43 kd protein. Neuron 3, 163–175 (1989). 10.1016/0896-6273(89)90029-9

4 Lin, W. et al. Distinct roles of nerve and muscle in postsynaptic differentiation of the neuromuscular synapse. Nature 410, 1057–1064 (2001). 10.1038/35074025

5 Apel, E. D., Roberds, S. L., Campbell, K. P. & Merlie, J. P. Rapsyn may function as a link between the acetylcholine receptor and the agrin-binding dystrophin-associated glycoprotein complex. Neuron 15, 115–126 (1995). 10.1016/0896-6273(95)90069-1

6 Belhasan, D. C. & Akaaboune, M. The role of the dystrophin glycoprotein complex on the neuromuscular system. Neurosci Lett 722, 134833 (2020). 10.1016/j.neulet.2020.134833

7 Bhat, H. F. et al. ABC of multifaceted dystrophin glycoprotein complex (DGC). J Cell Physiol 233, 5142–5159 (2018). 10.1002/jcp.25982

8 Ervasti, J. M. & Campbell, K. P. A role for the dystrophin-glycoprotein complex as a transmembrane linker between laminin and actin. J Cell Biol 122, 809–823 (1993). 10.1083/jcb.122.4.809

9 Grady, R. M. et al. Skeletal and cardiac myopathies in mice lacking utrophin and dystrophin: a model for Duchenne muscular dystrophy. Cell 90, 729–738 (1997). 10.1016/s0092-8674(00)80533-4

10 Grady, R. M. et al. Maturation and maintenance of the neuromuscular synapse: genetic evidence for roles of the dystrophin--glycoprotein complex. Neuron 25, 279–293 (2000). 10.1016/s0896-6273(00)80894-6

11 Kim, M. J., Whitehead, N. P., Bible, K. L., Adams, M. E. & Froehner, S. C. Mice lacking alpha-, beta1- and beta2-syntrophins exhibit diminished function and reduced dystrophin expression in both cardiac and skeletal muscle. Hum Mol Genet 28, 386–395 (2019). 10.1093/hmg/ddy341

12 Zaccaria, M. L., De Stefano, M. E., Gotti, C., Petrucci, T. C. & Paggi, P. Selective reduction in the nicotinic acetylcholine receptor and dystroglycan at the postsynaptic apparatus of mdx mouse superior cervical ganglion. J Neuropathol Exp Neurol 59, 103–112 (2000). 10.1093/jnen/59.2.103

13 Eckler, S. A., Kuehn, R. & Gautam, M. Deletion of N-terminal rapsyn domains disrupts clustering and has dominant negative effects on clustering of full-length rapsyn. Neuroscience 131, 661–670 (2005). 10.1016/j.neuroscience.2004.11.035

14 Ramarao, M. K., Bianchetta, M. J., Lanken, J. & Cohen, J. B. Role of rapsyn tetratricopeptide repeat and coiled-coil domains in self-association and nicotinic acetylcholine receptor clustering. J Biol Chem 276, 7475–7483 (2001). 10.1074/jbc.M009888200

15 Gautam, M., DeChiara, T. M., Glass, D. J., Yancopoulos, G. D. & Sanes, J. R. Distinct phenotypes of mutant mice lacking agrin, MuSK, or rapsyn. Brain Res Dev Brain Res 114, 171–178 (1999). 10.1016/s0165-3806(99)00013-9

16 Xing, G. et al. Membraneless condensates by Rapsn phase separation as a platform for neuromuscular junction formation. Neuron 109, 1963–1978 e1965 (2021). 10.1016/j.neuron.2021.04.021

17 Li, L. et al. Enzymatic Activity of the Scaffold Protein Rapsyn for Synapse Formation. Neuron 92, 1007–1019 (2016). 10.1016/j.neuron.2016.10.023

18 Dobbins, G. C., Luo, S., Yang, Z., Xiong, W. C. & Mei, L. alpha-Actinin interacts with rapsyn in agrin-stimulated AChR clustering. Mol Brain 1, 18 (2008). 10.1186/1756-6606-1-18

19 Oury, J. et al. MACF1 links Rapsyn to microtubule- and actin-binding proteins to maintain neuromuscular synapses. J Cell Biol 218, 1686–1705 (2019). 10.1083/jcb.201810023

20 Mihailovska, E. et al. Neuromuscular synapse integrity requires linkage of acetylcholine receptors to postsynaptic intermediate filament networks via rapsyn-plectin 1f complexes. Mol Biol Cell 25, 4130–4149 (2014). 10.1091/mbc.E14-06-1174

21 Case, L. B., Ditlev, J. A. & Rosen, M. K. Regulation of Transmembrane Signaling by Phase Separation. Annu Rev Biophys 48, 465–494 (2019). 10.1146/annurev-biophys-052118-115534

22 Kummer, T. T., Misgeld, T. & Sanes, J. R. Assembly of the postsynaptic membrane at the neuromuscular junction: paradigm lost. Curr Opin Neurobiol 16, 74–82 (2006). 10.1016/j.conb.2005.12.003

23 Gingras, J. et al. Alpha-Dystrobrevin-1 recruits Grb2 and alpha-catulin to organize neurotransmitter receptors at the neuromuscular junction. J Cell Sci 129, 898–911 (2016). 10.1242/jcs.181180

24 Grady, R. M. et al. Tyrosine-phosphorylated and nonphosphorylated isoforms of alpha-dystrobrevin: roles in skeletal muscle and its neuromuscular and myotendinous junctions. J Cell Biol 160, 741–752 (2003). 10.1083/jcb.200209045

25 Pawlikowski, B. T. & Maimone, M. M. Formation of complex AChR aggregates in vitro requires alpha-dystrobrevin. Dev Neurobiol 69, 326–338 (2009). 10.1002/dneu.20703

26 Schmidt, N. et al. Neuregulin/ErbB regulate neuromuscular junction development by phosphorylation of alpha-dystrobrevin. J Cell Biol 195, 1171–1184 (2011). 10.1083/jcb.201107083

27 Chen, G. et al. The 3BP2 adapter protein is required for chemoattractant-mediated neutrophil activation. J Immunol 189, 2138–2150 (2012). 10.4049/jimmunol.1103184

28 Deckert, M., Tartare-Deckert, S., Hernandez, J., Rottapel, R. & Altman, A. Adaptor function for the Syk kinases-interacting protein 3BP2 in IL-2 gene activation. Immunity 9, 595–605 (1998). 10.1016/s1074-7613(00)80657-3

29 Foucault, I. et al. The adaptor protein 3BP2 associates with VAV guanine nucleotide exchange factors to regulate NFAT activation by the B-cell antigen receptor. Blood 105, 1106–1113 (2005). 10.1182/blood-2003-08-2965

30 Hatani, T. & Sada, K. Adaptor protein 3BP2 and cherubism. Curr Med Chem 15, 549–554 (2008). 10.2174/092986708783769795

31 Jevremovic, D., Billadeau, D. D., Schoon, R. A., Dick, C. J. & Leibson, P. J. Regulation of NK cell-mediated cytotoxicity by the adaptor protein 3BP2. J Immunol 166, 7219–7228 (2001). 10.4049/jimmunol.166.12.7219

32 Levaot, N. et al. 3BP2-deficient mice are osteoporotic with impaired osteoblast and osteoclast functions. J Clin Invest 121, 3244–3257 (2011). 10.1172/JCI45843

33 Levaot, N. et al. Loss of Tankyrase-mediated destruction of 3BP2 is the underlying pathogenic mechanism of cherubism. Cell 147, 1324–1339 (2011). 10.1016/j.cell.2011.10.045

34 Sada, K. et al. Regulation of FcepsilonRI-mediated degranulation by an adaptor protein 3BP2 in rat basophilic leukemia RBL-2H3 cells. Blood 100, 2138–2144 (2002). 10.1182/blood-2001-12-0340

35 Ueki, Y. et al. Mutations in the gene encoding c-Abl-binding protein SH3BP2 cause cherubism. Nat Genet 28, 125–126 (2001). 10.1038/88832

36 Aittaleb, M., Martinez-Pena, Y. V. I. & Akaaboune, M. Spatial distribution and molecular dynamics of dystrophin glycoprotein components at the neuromuscular junction in vivo. J Cell Sci 130, 1752–1759 (2017). 10.1242/jcs.198358

37 Kummer, T. T., Misgeld, T., Lichtman, J. W. & Sanes, J. R. Nerve-independent formation of a topologically complex postsynaptic apparatus. J Cell Biol 164, 1077–1087 (2004). 10.1083/jcb.200401115

38 Bruneau, E., Sutter, D., Hume, R. I. & Akaaboune, M. Identification of nicotinic acetylcholine receptor recycling and its role in maintaining receptor density at the neuromuscular junction in vivo. J Neurosci 25, 9949–9959 (2005). 10.1523/JNEUROSCI.3169-05.2005

39 Bruneau, E. G., Macpherson, P. C., Goldman, D., Hume, R. I. & Akaaboune, M. The effect of agrin and laminin on acetylcholine receptor dynamics in vitro. Dev Biol 288, 248–258 (2005). 10.1016/j.ydbio.2005.09.041

40 Proszynski, T. J., Gingras, J., Valdez, G., Krzewski, K. & Sanes, J. R. Podosomes are present in a postsynaptic apparatus and participate in its maturation. Proc Natl Acad Sci U S A 106, 18373–18378 (2009). 10.1073/pnas.0910391106

41 Colledge, M. & Froehner, S. C. Tyrosine phosphorylation of nicotinic acetylcholine receptor mediates Grb2 binding. J Neurosci 17, 5038–5045 (1997). 10.1523/JNEUROSCI.17-13-05038.1997

42 Hall, Z. W. & Sanes, J. R. Synaptic structure and development: the neuromuscular junction. Cell 72 Suppl, 99–121 (1993). 10.1016/s0092-8674(05)80031-5

43 Lee, Y., Rudell, J. & Ferns, M. Rapsyn interacts with the muscle acetylcholine receptor via alpha-helical domains in the alpha, beta, and epsilon subunit intracellular loops. Neuroscience 163, 222–232 (2009). 10.1016/j.neuroscience.2009.05.057

44 Boeynaems, S. et al. Protein Phase Separation: A New Phase in Cell Biology. Trends Cell Biol 28, 420–435 (2018). 10.1016/j.tcb.2018.02.004

45 Wu, X., Cai, Q., Feng, Z. & Zhang, M. Liquid-Liquid Phase Separation in Neuronal Development and Synaptic Signaling. Dev Cell 55, 18–29 (2020). 10.1016/j.devcel.2020.06.012

46 Krypotou, E. et al. Bacteria require phase separation for fitness in the mammalian gut. Science 379, 1149–1156 (2023). 10.1126/science.abn7229

47 Milovanovic, D., Wu, Y., Bian, X. & De Camilli, P. A liquid phase of synapsin and lipid vesicles. Science 361, 604–607 (2018). 10.1126/science.aat5671

48 Ramarao, M. K. & Cohen, J. B. Mechanism of nicotinic acetylcholine receptor cluster formation by rapsyn. Proc Natl Acad Sci U S A 95, 4007–4012 (1998). 10.1073/pnas.95.7.4007

49 Tallquist, M. D., Weismann, K. E., Hellstrom, M. & Soriano, P. Early myotome specification regulates PDGFA expression and axial skeleton development. Development 127, 5059–5070 (2000). 10.1242/dev.127.23.5059

50 Jones, R. A. et al. NMJ-morph reveals principal components of synaptic morphology influencing structure-function relationships at the neuromuscular junction. Open Biol 6 (2016). 10.1098/rsob.160240

51 Sanes, J. R. & Lichtman, J. W. Induction, assembly, maturation and maintenance of a postsynaptic apparatus. Nat Rev Neurosci 2, 791–805 (2001). 10.1038/35097557

52 Shi, L., Fu, A. K. & Ip, N. Y. Molecular mechanisms underlying maturation and maintenance of the vertebrate neuromuscular junction. Trends Neurosci 35, 441–453 (2012). 10.1016/j.tins.2012.04.005

53 Legay, C. Why so many forms of acetylcholinesterase? Microsc Res Tech 49, 56–72 (2000). 10.1002/(SICI)1097-0029(20000401)49:1<56::AID-JEMT7>3.0.CO;2-R

54 Rossi, S. G. & Rotundo, R. L. Localization of “non-extractable” acetylcholinesterase to the vertebrate neuromuscular junction. J Biol Chem 268, 19152–19159 (1993).

55 Stanco, A. M. & Werle, M. J. Agrin and acetylcholine receptors are removed from abandoned synaptic sites at reinnervated frog neuromuscular junctions. J Neurobiol 33, 999–1018 (1997).

56 Chen, F. et al. Neuromuscular synaptic patterning requires the function of skeletal muscle dihydropyridine receptors. Nat Neurosci 14, 570–577 (2011). 10.1038/nn.2792

57 Nelson, B. R. et al. Skeletal muscle-specific T-tubule protein STAC3 mediates voltage-induced Ca2+ release and contractility. Proc Natl Acad Sci U S A 110, 11881–11886 (2013). 10.1073/pnas.1310571110

58 Eiber, N. et al. Lack of Desmin in Mice Causes Structural and Functional Disorders of Neuromuscular Junctions. Front Mol Neurosci 13, 567084 (2020). 10.3389/fnmol.2020.567084

59 Reichenberger, E. J., Levine, M. A., Olsen, B. R., Papadaki, M. E. & Lietman, S. A. The role of SH3BP2 in the pathophysiology of cherubism. Orphanet J Rare Dis 7 Suppl 1, S5 (2012). 10.1186/1750-1172-7-S1-S5

60 Bernadzki, K. M. et al. Liprin-alpha-1 is a novel component of the murine neuromuscular junction and is involved in the organization of the postsynaptic machinery. Sci Rep 7, 9116 (2017). 10.1038/s41598-017-09590-7

61 Cote, P. D., Moukhles, H., Lindenbaum, M. & Carbonetto, S. Chimaeric mice deficient in dystroglycans develop muscular dystrophy and have disrupted myoneural synapses. Nat Genet 23, 338–342 (1999). 10.1038/15519

62 Deconinck, A. E. et al. Utrophin-dystrophin-deficient mice as a model for Duchenne muscular dystrophy. Cell 90, 717–727 (1997). 10.1016/s0092-8674(00)80532-2

63 Duclos, F. et al. Progressive muscular dystrophy in alpha-sarcoglycan-deficient mice. J Cell Biol 142, 1461–1471 (1998). 10.1083/jcb.142.6.1461

64 Williamson, R. A. et al. Dystroglycan is essential for early embryonic development: disruption of Reichert’s membrane in Dag1-null mice. Hum Mol Genet 6, 831–841 (1997). 10.1093/hmg/6.6.831

65 Grady, R. M., Merlie, J. P. & Sanes, J. R. Subtle neuromuscular defects in utrophin-deficient mice. J Cell Biol 136, 871–882 (1997). 10.1083/jcb.136.4.871

66 Kameya, S. et al. alpha1-syntrophin gene disruption results in the absence of neuronal-type nitric-oxide synthase at the sarcolemma but does not induce muscle degeneration. J Biol Chem 274, 2193–2200 (1999). 10.1074/jbc.274.4.2193

67 Oak, S. A., Russo, K., Petrucci, T. C. & Jarrett, H. W. Mouse alpha1-syntrophin binding to Grb2: further evidence of a role for syntrophin in cell signaling. Biochemistry 40, 11270–11278 (2001). 10.1021/bi010490n

68 Yang, B. et al. SH3 domain-mediated interaction of dystroglycan and Grb2. J Biol Chem 270, 11711–11714 (1995). 10.1074/jbc.270.20.11711

69 Banks, G. B., Chamberlain, J. S. & Froehner, S. C. Truncated dystrophins can influence neuromuscular synapse structure. Mol Cell Neurosci 40, 433–441 (2009). 10.1016/j.mcn.2008.12.011

70 Martinez-Martinez, P. et al. Silencing rapsyn in vivo decreases acetylcholine receptors and augments sodium channels and secondary postsynaptic membrane folding. Neurobiol Dis 35, 14–23 (2009). 10.1016/j.nbd.2009.03.008

71 Huebsch, K. A. & Maimone, M. M. Rapsyn-mediated clustering of acetylcholine receptor subunits requires the major cytoplasmic loop of the receptor subunits. J Neurobiol 54, 486–501 (2003). 10.1002/neu.10177

72 Marchand, S., Bignami, F., Stetzkowski-Marden, F. & Cartaud, J. The myristoylated protein rapsyn is cotargeted with the nicotinic acetylcholine receptor to the postsynaptic membrane via the exocytic pathway. J Neurosci 20, 521–528 (2000). 10.1523/JNEUROSCI.20-02-00521.2000

73 Alvarez-Suarez, P. et al. Drebrin Regulates Acetylcholine Receptor Clustering and Organization of Microtubules at the Postsynaptic Machinery. Int J Mol Sci 22 (2021). 10.3390/ijms22179387

74 Antolik, C. et al. The actin binding domain of ACF7 binds directly to the tetratricopeptide repeat domains of rapsyn. Neuroscience 145, 56–65 (2007). 10.1016/j.neuroscience.2006.11.047

75 Feng, Z., Chen, X., Zeng, M. & Zhang, M. Phase separation as a mechanism for assembling dynamic postsynaptic density signalling complexes. Curr Opin Neurobiol 57, 1–8 (2019). 10.1016/j.conb.2018.12.001

76 Zeng, M. et al. Phase Transition in Postsynaptic Densities Underlies Formation of Synaptic Complexes and Synaptic Plasticity. Cell 166, 1163–1175 e1112 (2016). 10.1016/j.cell.2016.07.008

77 Matsumoto, Y. et al. RANKL coordinates multiple osteoclastogenic pathways by regulating expression of ubiquitin ligase RNF146. J Clin Invest 127, 1303–1315 (2017). 10.1172/JCI90527

78 Madisen, L. et al. A robust and high-throughput Cre reporting and characterization system for the whole mouse brain. Nat Neurosci 13, 133–140 (2010). 10.1038/nn.2467

79 Grady, R. M. et al. Role for alpha-dystrobrevin in the pathogenesis of dystrophin-dependent muscular dystrophies. Nat Cell Biol 1, 215–220 (1999). 10.1038/12034

80 Shyu, Y. J., Liu, H., Deng, X. & Hu, C. D. Identification of new fluorescent protein fragments for bimolecular fluorescence complementation analysis under physiological conditions. Biotechniques 40, 61–66 (2006). 10.2144/000112036

81 Tanaka, M., Gupta, R. & Mayer, B. J. Differential inhibition of signaling pathways by dominant-negative SH2/SH3 adapter proteins. Mol Cell Biol 15, 6829–6837 (1995). 10.1128/MCB.15.12.6829

82 Pezinski, M., Daszczuk, P., Pradhan, B. S., Lochmuller, H. & Proszynski, T. J. An improved method for culturing myotubes on laminins for the robust clustering of postsynaptic machinery. Sci Rep 10, 4524 (2020). 10.1038/s41598-020-61347-x

83 Doss, S. V. et al. Expression and Roles of Lynx1, a Modulator of Cholinergic Transmission, in Skeletal Muscles and Neuromuscular Junctions in Mice. Front Cell Dev Biol 10, 838612 (2022). 10.3389/fcell.2022.838612

84 Xing, G. et al. A mechanism in agrin signaling revealed by a prevalent Rapsyn mutation in congenital myasthenic syndrome. Elife 8 (2019). 10.7554/eLife.49180

85 Bernadzki, K. M. et al. Arhgef5 Binds alpha-Dystrobrevin 1 and Regulates Neuromuscular Junction Integrity. Front Mol Neurosci 13, 104 (2020). 10.3389/fnmol.2020.00104

86 Kobayashi, Y. M. et al. Sarcolemma-localized nNOS is required to maintain activity after mild exercise. Nature 456, 511–515 (2008). 10.1038/nature07414

87 Zhao, K. et al. Muscle Yap Is a Regulator of Neuromuscular Junction Formation and Regeneration. J Neurosci 37, 3465–3477 (2017). 10.1523/JNEUROSCI.2934-16.2017

88 Wupper, S. et al. Effects of dietary gamma-cyclodextrin on voluntary activity and muscle strength in mice. J Physiol Pharmacol 71 (2020). 10.26402/jpp.2020.3.08

89 Guo, S. et al. Impacts of exercise interventions on different diseases and organ functions in mice. J Sport Health Sci 9, 53–73 (2020). 10.1016/j.jshs.2019.07.004

90 Eiber, N. et al. Loss of Protein Kinase Csnk2b/CK2beta at Neuromuscular Junctions Affects Morphology and Dynamics of Aggregated Nicotinic Acetylcholine Receptors, Neuromuscular Transmission, and Synaptic Gene Expression. Cells 8 (2019). 10.3390/cells8080940

